# Multiplexing information flow through dynamic signalling systems

**DOI:** 10.1101/863159

**Authors:** Giorgos Minas, Dan J. Woodcock, Louise Ashall, Claire V. Harper, Michael R. H. White, David A Rand

## Abstract

We consider how a signalling system can act as an information hub by multiplexing information arising from multiple signals. We formally define multiplexing, mathematically characterise which systems can multiplex and how well they can do it. While the results of this paper are theoretical, to motivate the idea of multiplexing, we provide experimental evidence that tentatively suggests that the NF-*κ*B transcription factor can multiplex information about changes in multiple signals. We believe that our theoretical results may resolve the apparent paradox of how a system like NF-*κ*B that regulates cell fate and inflammatory signalling in response to diverse stimuli can appear to have the low information carrying capacity suggested by recent studies on scalar signals. In carrying out our study, we introduce new methods for the analysis of large, nonlinear stochastic dynamic models, and develop computational algorithms that facilitate the calculation of fundamental constructs of information theory such as Kullback–Leibler divergences and sensitivity matrices, and link these methods to new theory about multiplexing information. We show that many current models such as those of the NF-*κ*B system cannot multiplex effectively and provide models that overcome this limitation using post-transcriptional modifications.

## 1 Introduction

Signalling systems provide a very important example of cellular information systems since they transmit information arising from inside and outside the cell to the cell’s processing units. For example, it is generally believed that the NF-*κ*B system uses the information from a large number of input signals (see Fig. 1(a)) to regulate gene transcription of more than 500 genes in a highly versatile way [9, 38]. NF-*κ*B regulates cell fate and inflammatory signalling in response to diverse stimuli, including changes in temperature [10], viral and bacterial pathogens, free radicals, cytokines, and growth factors [38]. Thus, we have a situation where both the input signal that encodes information about the cell’s environment, and the gene response are multi-dimensional. Such a system is often referred to as an information hub.

**Figure 1:**
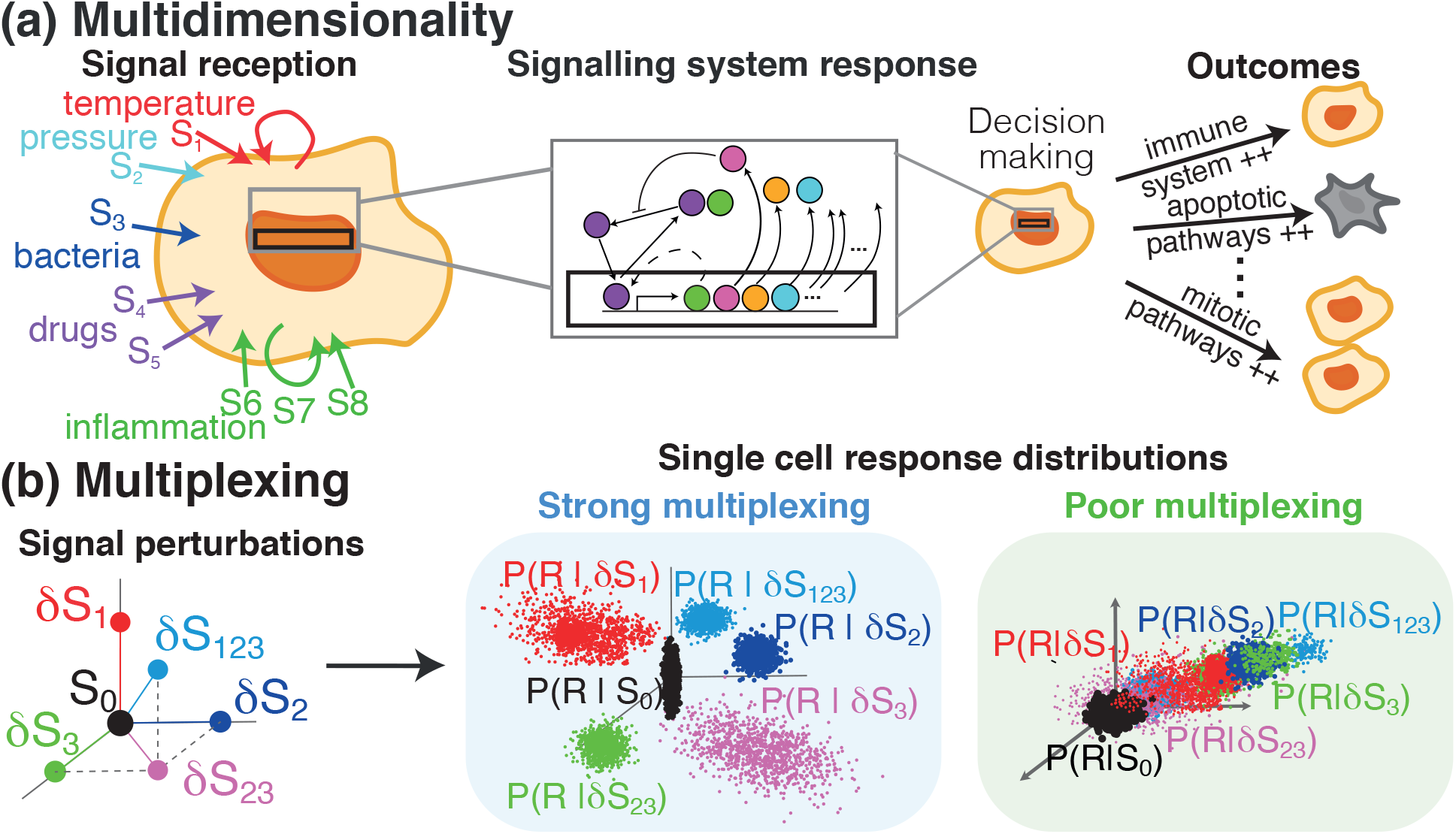
Multiplexing signals through signalling systems. (a) Cells constantly receive a multitude of different signals in which signalling systems respond by (directly or indirectly) modulating the expression of a number of target genes. These target genes activate or not various pathways of the cell which leads to completely different cell outcomes from cell survival to apoptosis or mitosis. In order for this decision making to be done reliable and robust, signalling systems need to have the capacity to multiplex a variety of simultaneously arising signals. (b) Multiplexing is defined as the ability of the signalling system response to identify which of the input signals have changed. In broad terms, strong multiplexing is evident by the probability distributions of the signalling system response in a population of single cells being significantly different for the different regimes of the multi-dimensional signal. On contrary, poor multiplexing leads to response distribution that are very similar for different signals.

This raises the question of to what extent this process is mediated through the signalling system itself which may have a single transcription factor, rather than through multiple other parallel pathways that also provide information to the genome. Can such a signalling system on its own effectively regulate a relationship between multidimensional inputs and responses that can robustly and reliably modulate decision-making of the claimed versatility without using other pathways? This is the central question we consider here.

We consider this question in terms of multiplexing which we define as follows. We suppose that our system has multiple input signals *S*_1_,… *S_s_* and consider how the system responds to changes in them. These signals might, for instance, be changes in temperature or other physical parameters (e.g. pressure or humidity), changes in the level and/or timing pattern of an activator (e.g. tumor necrosis factor-*α* (TNF*α*), interleukin 1*β* (IL-1*β*), and Lipopolysaccharides (LPS) for NF-*κ*B), and/or drug treatments (e.g. Diclofenac for NF-*κ*B). We say that a signalling system can multiplex these input signals *S*_1_,… *S_s_* if one can reliably determine which of these input signals have changed using only the multidimensional response of the target genes (see Fig. 1(B)). It is this response that will regulate downstream responses of the cell and therefore the multiplexing capacity is directly measuring a key aspect of how effectively the cell can respond to the multiple inputs.

So far as we are aware this is a new approach and consequently an immediate question is whether there is any experimental evidence that this is the case. We address this below and present some tentative evidence for it. This is useful because it provides useful context to our discussion and give some helpful insights but we must emphasise that this evidence is very far from proving such a point even though it is highly suggestive.

To address the question of what aspects of the system enable such multiplexing we will introduce a quantity, called the *multiplexing capacity*, that measures the ability of a noisy signalling system to multiplex a set of signals. Using this we demonstrate that while current models cannot multiplex effectively, biologically natural modifications of them can. We believe this indicates general principles behind biological design.

The underlying concepts that we use are connected to important tools involved in systems identification, sensitivity analysis and information geometry such as the Fisher Information Matrix (FIM) and the Kullback-Leibler divergence [6] and we introduce and calculate a new, but related, sensitivity matrix **<u>s</u>** that characterises the multiplexing capacity. While sensitivity analysis [3, 26, 27] is an extensive area for deterministic dynamics (e.g. [54, 8]) it is less well developed for stochastic systems (e.g. [51, 17, 55, 52, 53, 56, 57]). Since it is crucial in our discussion that we take account of realistic levels of stochasticity for the systems we consider, calculating the relevant quantities for complex high-dimensional stochastic systems such as those considered here is therefore a significant mathematical challenge. To overcome this, in the numerical computations we use the pcLNA method [18] that allows fast and accurate computation of key information theoretic quantities, such as Kullback-Leibler divergences and the Fisher Information matrix, for stochastic dynamical systems.

Given that NF-*κ*B has complex oscillatory dynamics, an obvious hypothesis is that it is this dynamical behavior of the system that allows it to act as an information hub. However, we will use our theoretical tools to provide evidence that this is not the case and show that the NF-*κ*B system described by current models cannot multiplex effectively even though it has complex oscillatory dynamics. On the other hand, we will demonstrate how to modify a stochastic model of NF-*κ*B so as to overcome this inability to multiplex. In particular, we show that additional regulated states of NF-*κ*B, which might include differential post-translational modifications and/or differential hetero- and homo-dimerisation, can enable such multiplexing and that the oscillatory dynamics can greatly enrich the multiplexing capacity in this modified model.

Recent important papers studied the information flow through biochemical systems such as the nuclear factor-*κ*B (NF-*κ*B), calcium (Ca^2+^), and extracellular signal-regulated kinase (ERK). The focus was on measuring how much information is being carried by the signalling systems in terms of the mutual information *I*(*S, R*) and the capacity of the channel *S* → *P*(*R|S*) [6] where *S* is the input signal and *P*(*R|S*) the probability distribution of a response *R*. For example, for NF-*κ*B the signal *S* was the level of TNF*α* stimulation and *R* was the level of transcription factor in the nucleus at one or more timepoints. In summary, the channel capacity was estimated to be around 1 bit for static scalar observations in response to one-dimensional stimuli [2, 5, 21, 28, 34], about 1.5 bits when the dynamical behaviour of the system response is considered [28] and up to 1.7 bits when cell-to-cell heterogeneity is accounted for [37]. A number of recent studies support this core observation and report similar low channel capacities [12, 13, 15, 16, 33, 49]. The stochastic models that we use reproduce these relatively low levels of mutual information between TNF*α* level and total transcription factor abundance and also agree with that of the stochastic model in [31]. On the other hand, the modified versions allow significantly greater mutual information for multidimensional inputs.

The logic of our discussion is as follows: Firstly, we discuss some tentative experimental evidence suggesting that NF-*κ*B does multiplex. Then we introduce a method for quantifying the ability of a given stochastic model to multiplex and show that current models are poor at this. We suggest how one can modify the models so as to enable better multiplexing and relate this to known mechanisms in signalling systems. In particular, we provide a modified model that is able to reproduce the behaviour of the simpler of the two multiplexing experimental systems we discuss. We finally discuss how these results relate to the low information capacity found in previous studies. Our analysis gives important insight into how multiplexing can work in a signalling system, however we are not claiming that our model is a true representation of the real biology of those systems.

### Does NF-*κ*B multiplex?

To illustrate the above characterisation of multiplexing we consider some experimental evidence. We ask if, by monitoring the response of a set of genes that are direct NF-*κ*B targets (SI Sect. 5), we can reliably determine the state of a multidimensional input signal. We firstly consider the response of three important genes, EGR1, COX-2 (PTGS2) and IL-8 (CXCL-8), to pulses of varying length repeated every 100 minutes at two temperatures, 37°C and 40°C and ask if, from the response of these genes, we can determine the temperature and pulsing length. EGR1 regulates the response to growth factors, DNA damage, and ischemia, preventing tumor formation by activating p53/TP53 and TGFB1. COX-2 is responsible for production of inflammatory prosta-glandins. We include the chemokine gene IL-8 (CXCL-8) to distinguish temperature at 30mins but there are a small number of other NF-*κ*B target genes such as NUAK2, NFKBIA, TNFAIP3 (A20) that could have been used instead (see [10]). We use microarrays and RT-qPCR data to monitor the expression of these genes around the peak times of nuclear NF-*κ*B at 0, 30, 130, 230 and 430 minutes (Fig. 2a). We see that, if we know the expression levels of these genes at these times we can determine which of these multiple experiments was carried out (Fig. 2b). Monitoring the gene expression at 30 minutes enables the identification of 5 distinct input signal combinations (unstimulated and the four combinations in the table) and also one additional one if observations at 130 minutes are included. This suggests that just from monitoring these three genes we obtain at least 2 bits of information.

**Figure 2:**
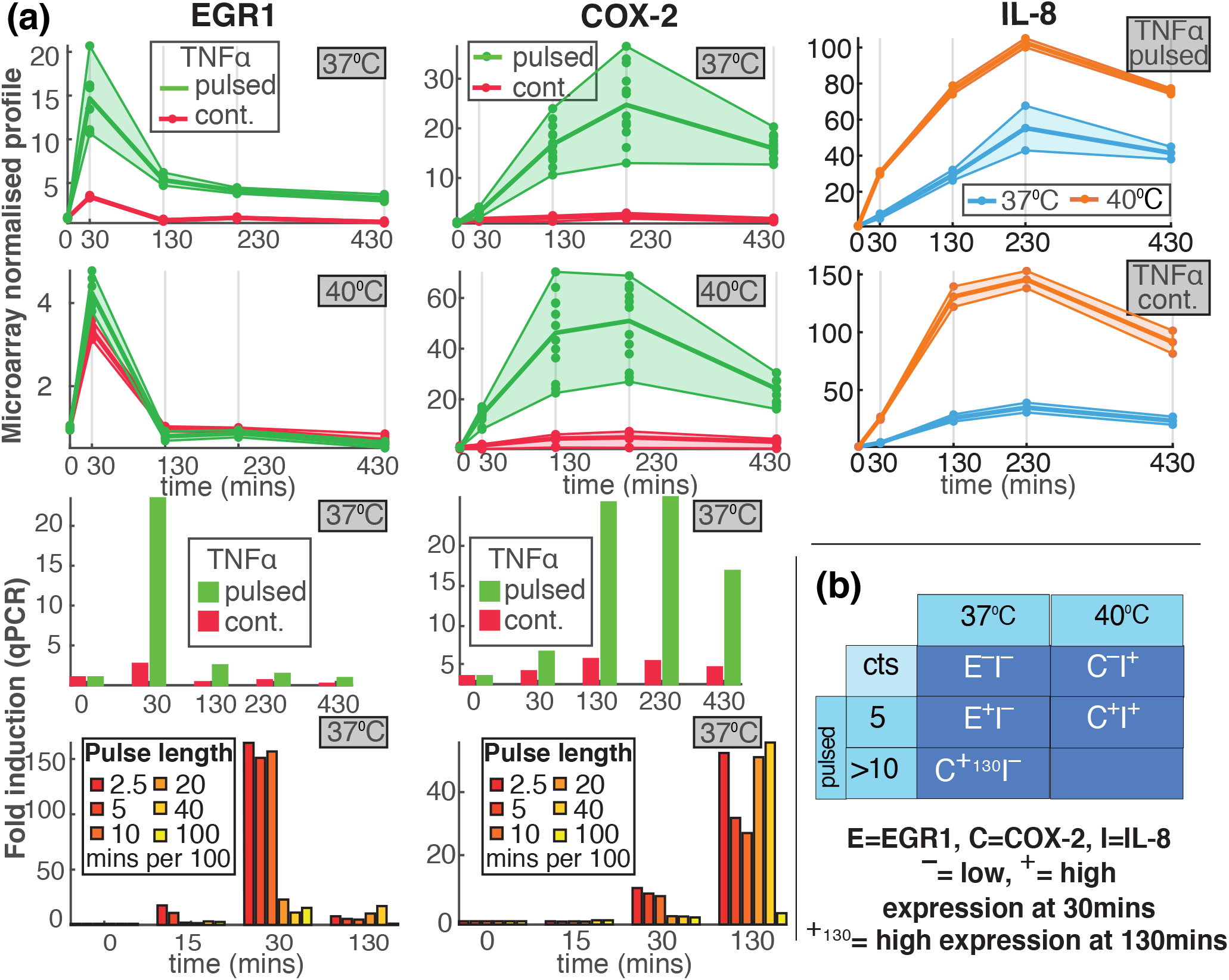
Gene expression can identify different experimental conditions. (a) (Left Column) The expression of the gene EGR1 in normal (37°C) and high (40°C) temperature and under continuous or pulsed TNF*α* treatment with pulses repeated every 100 minutes. The pulse length is 5 minutes except in the bottom row where the pulse length is indicated in the legend. (Middle Column) As Left Column except that the gene is COX-2 (PTGS2). (Right Column) As the first two rows of the Left Column except that the gene is IL-8 (CXCL-8). (b) A table showing which gene expression combinations identify which pairs of the input signal. The letters E, C and I indicate the genes EGR1, COX-2 and IL-8 respectively. The plus symbol (+) indicates high expression at 30 minutes and the minus symbol (-) indicates low expression at this time. Thus *E*^+^*I*^−^ indicates that EGR1 is highly expressed at 30 minutes and IL-8 is then at a low level, which implies that the system is pulsed with 5 minute pulses and the temperature is 37°C. The symbol +_130_ indicates high expression at 130 minutes.

There are two substantial caveats to this observation. Firstly, a potential criticism is that the genes discussed might also be regulated by independent parallel pathways. However, we note that in Fig. 2(b) we have restricted to very early observations at 30 minutes when the involvement of other pathways is unlikely. Secondly, we are using data from cell population assays such as microarrays rather than single cells where stochastic effects are important. However, the expression difference in the table are more than two logs so the overlap of the corresponding expression distribution should be small.

Despite these two caveats these results are highly suggestive and motivate a careful consideration of single cell multiplexing. Further information supporting these ideas is contained in the SI (Sect. 5 and Fig. S11).

## Results

### Decision-making and KL divergence

We now develop a mathematical approach that enables us to quantify multiplexing. We use this to show why current tightly coupled models of NF-*κ*B cannot multiplex effectively and then explain how to modify these so that multiplexing is enabled.

Suppose we have *s* signals *S*_1_,…, *S_s_* which in turn define the vector signal ***S*** = (*S*_1_,…, *S_s_*). Consider a change in the signal from a base value ***S***_0_ to ***S*** = ***S***_0_ + ***δS*** where the change has size *η* = ||***δS***||. We ask whether, the response ***R*** has the capacity to distinguish which components of the signal have significantly changed i.e. to identify which components of ***δS*** are ≥ O(*η*). If it can we say the system can *multiplex*.

Mathematically the question of using the stochastic response ***R***, which has probability distribution *P*_***S***_(***R***) = *P*(***R|S***), to distinguish input signals is related to hypothesis testing. If a change in input signal occurs (say from ***S***_0_ to ***S*** = ***S***_0_ + ***δS***) and we wish to determine if the *i*th component was changed using only ***R***, we need to be able to evaluate the hypothesis that ***R*** comes from *P*_***S***_ rather than from a distribution of the form *P*_***S′***_ where ***S′*** is any perturbation of ***S***_0_ with the same *i*th component as ***S***_0_. By the Neyman-Pearson lemma, the most powerful test of this hypothesis for a given false-positive error rate *α* is a test of the form λ(***R***) ≥ *u_α_* where

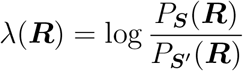

is the log-likelihood ratio and the choice of *α* determines what threshold *u_α_* to use. The *P*_***S***_-mean of the log-likelihood ratio is by definition the Kullback-Leibler (KL) divergence, *D*_KL_(*P*_***S***_||*P*_***S′***_), of *P*_***S***_ and *P*_***S′***_ distributions. The larger is the likelihood ratio, the more evidence we have in favour of signal ***S*** and against ***S′***.

### Multiplexing capacity

If *D*_KL_ is too small then the most powerful test is expected to fail and hence other tests will not fair any better. Furthermore, as we wish to check whether the response ***R*** has the capacity to distinguish ***S*** from *any* signal ***S′*** that has the *i*-th signal unchanged, we study how large is the 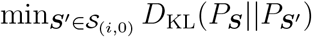 where 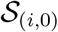 is the set of *all* such ***S′*** signals. However, as ***S*** tends towards ***S***_0_ thus decreasing the length *l* = *l*(***S***) = ||***S*** – ***S***_0_||, this quantity decreases like *l*^2^, and therefore we scale it and define

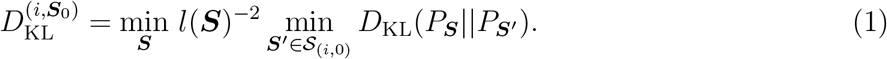

The larger 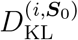 is, the easier it is to detect the change in the *i*th component.

To apply this so as to detect changes in any component of the signal ***S*** we consider

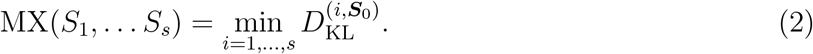

The larger this *multiplexing capacity* MX(*S*_1_,… *S_s_*) is, the better the system at multiplexing the signals *S*_1_,… *S_s_* (see also SI Sects. 2.1-2.4).

### Characterising multiplexing via the sensitivity matrix

While we cannot calculate this quantity in general, we can find an elegant solution in terms of the Fisher Information Matrix (FIM) when the changes in the signal are small, that is the third order terms and above are negligible. That is, we will calculate *D*_KL_(*P*_***S***_||*P*_***S′***_) up to terms that are O(max{||***S*** – ***S′***||^3^, ||***S*** – ***S′***||^3^, ||**S**′ – **S**_0_||^3^, }).

In our context, the FIM 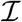 at ***S***_0_ has entries

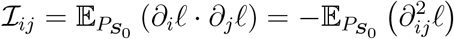

where *ℓ*(***S; R***) = log *P*(***R|S***) is the log–likelihood function, *∂_i_ℓ* denotes the partial derivative with respect to the *i*th component *S_i_* and 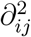 is the corresponding second derivative. These derivatives are evaluated at ***S***_0_.

The FIM measures the sensitivity of *P*_***S***_(***R***) to a change ***δS*** in the signal ***S*** because, up to terms that are O(||***δS***||^3^) (see SI Sect. 2.1),

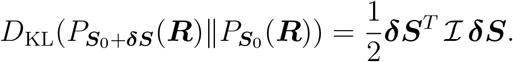

### Multiplexing sensitivity matrix

One can associate to the FIM 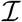 an *s* × *s* matrix **<u>s</u>** that satisfies 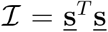 and certain optimality properties described in [18] (SI Sect. 2.5.2). For *j* = 1, 2,…, *s*, we call the entries *<u>s</u>_ij_* of the matrix **<u>s</u>**, the *principal coefficients of sensitivity* of the response ***R*** to the *j*-th signal *S_j_*.

The multiplexing sensitivity matrix **<u>s</u>** describes the ability of the signalling system to multiplex at least locally in the following way. If **<u>s</u>**_*j*_, *j*=1,…, *s*, are the columns of **<u>s</u>**, then

a. **<u>s</u>**_*i*_1_,…,*i_k_*_ denotes the linear subspace of 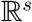 spanned by the vectors **<u>s</u>**_*i*_1_,…, **<u>s</u>***i_k_*_, and
b. **n** = **n**(*i*|*i*_1_,…, *i_k_*) denotes the component of **<u>s</u>**_*i*_ normal to the linear subspace **<u>s</u>***_i_1_,…, i_k__* i.e. **<u>s</u>**_*i*_ = **u** + **n** with **u** in **<u>s</u>**_*i*_1_,…, *i_k_*_ and **n** orthogonal to **<u>s</u>**_*i*_1_,…, *i_k_*_. If *i*_1_,…, *i_k_* include all indices except *j* we use the notation **n**(*i|j* ≠ *i*) for **n**(*i*|*i*_1_,…, *i_k_*).

Firstly, up to third order terms (see SI section 2.4),

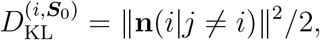

and therefore the length of the normal component, **n**(*i|j* ≠ *i*), determines, at least locally, the capacity of the response ***R*** to distinguish the i-th from the rest of the considered signals. Secondly, there is an essentially unique reordering of the signal components as *S_i_1__,…, S_i_s__* so that if *v_k_* = ||**n**(*i_k_*|*i*_1_,…, *i*_*k*−1_)|| then *v*_1_ ≥ ⋯ ≥ *v_s_* and the multiplexing capacities up to third order terms are

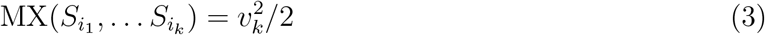

for all *k* = 1,…, *s*. All of these quantities can be rapidly calculated using the QR decompositions of submatrices of **<u>s</u>** made up from the relevant columns of **<u>s</u>** (see SI section 2.4).

This ordering of the set of signals provides a way to choose an optimal subset that can multiplex. That is, we can use the ordering *i*_1_,…, *i_s_* and the associated multiplexing capacities MX(*S_i_1__,…, S_i_k__*), *k* = 1,…, *s*, to identify the subset of signals with the largest number of elements, *k*, that has multiplexing capacity MX(*S_i_1__,…, S_i_k__*) ≥ *m*, for *m* an appropriate threshold (e.g. the minimum *D*_KL_ level for the change to be detectable in a given system of interest, see also SI section 2.4.1).

### Multiplexing capacities of a model

In regulatory and signalling systems, the values of two parameters, say 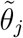 and 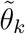, may differ by an order of magnitude or more. Therefore, when discussing sensitivities it is usually not appropriate to consider the absolute changes in the parameters 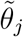, but instead to consider the relative changes. A good way to do this is to introduce new parameters 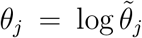 because absolute changes in *θ_j_* correspond to relative changes in 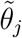. Then, for small changes 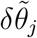 to the parameters, 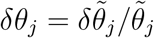 and so the changes *δθ_j_* are scaled and non-dimensional. When discussing the multiplexing capacities of a model below we will always use these scaled parameters *θ_j_* and henceforth when we refer to parameters *θ_i_* we mean these scaled parameters and we drop the tilde.

As mentioned above the typical situation is where the signals *S_i_* change the parameters *θ_j_*, *j* = 1,…, *s*, of the model so that *θ* = *θ*(***S***). In this case we show in Sect. 2.5.3 of the SI how one can relate the multiplexing capacity of the signals to the multiplexing capacity of the parameters. For the latter we regard the parameters as signals and calculate their multiplexing capacities 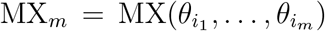 where the parameters *θ_i_* have been reordered so that for all 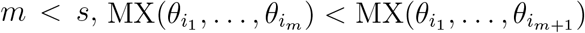. We call these the *multiplexing capacities of the model*.

Knowing the multiplexing capacities of a model is important because, using equation (15)of SI Sect. 2.5.3, we can tell how well the system can multiplex any signals changing these parameters. In particular,

a. if the MX_*m*_ very rapidly decrease with *m* then the system is not able to multiplex through these parameters; and
b. if 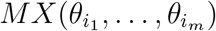 is large and there are signals *S*_1_,…, *S_r_* which change the parameters 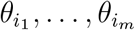 in that 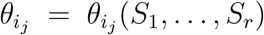, then these signals will have good multiplexing properties provided the matrix *d**θ***(***S***) (which is the derivative of ***θ*** with respect to ***S*** evaluated at the base value of ***S***) is well conditioned (e.g. if det *d**θ***(***S***) is not too small or big).

### Tightly coupled models of NF-*κ*B cannot multiplex effectively

One might expect that a dynamical system with many parameters, such as NF-*κ*B (see Fig. 3(a)), would have the flexibility to multiplex effectively. However, it has been observed that for a large class of deterministic models of regulatory and signalling systems of the sort that we are considering, the deterministic analogue of the FIM for the model parameters has rapidly decreasing eigenvalues 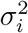 [8, 23, 24, 25, 32, 35]. A similar result was shown for stochastic models of the circadian clock in [18, 58]. This implies that the effects of changing different parameters are highly correlated making it hard to recognise which parameter was changed.

**Figure 3:**
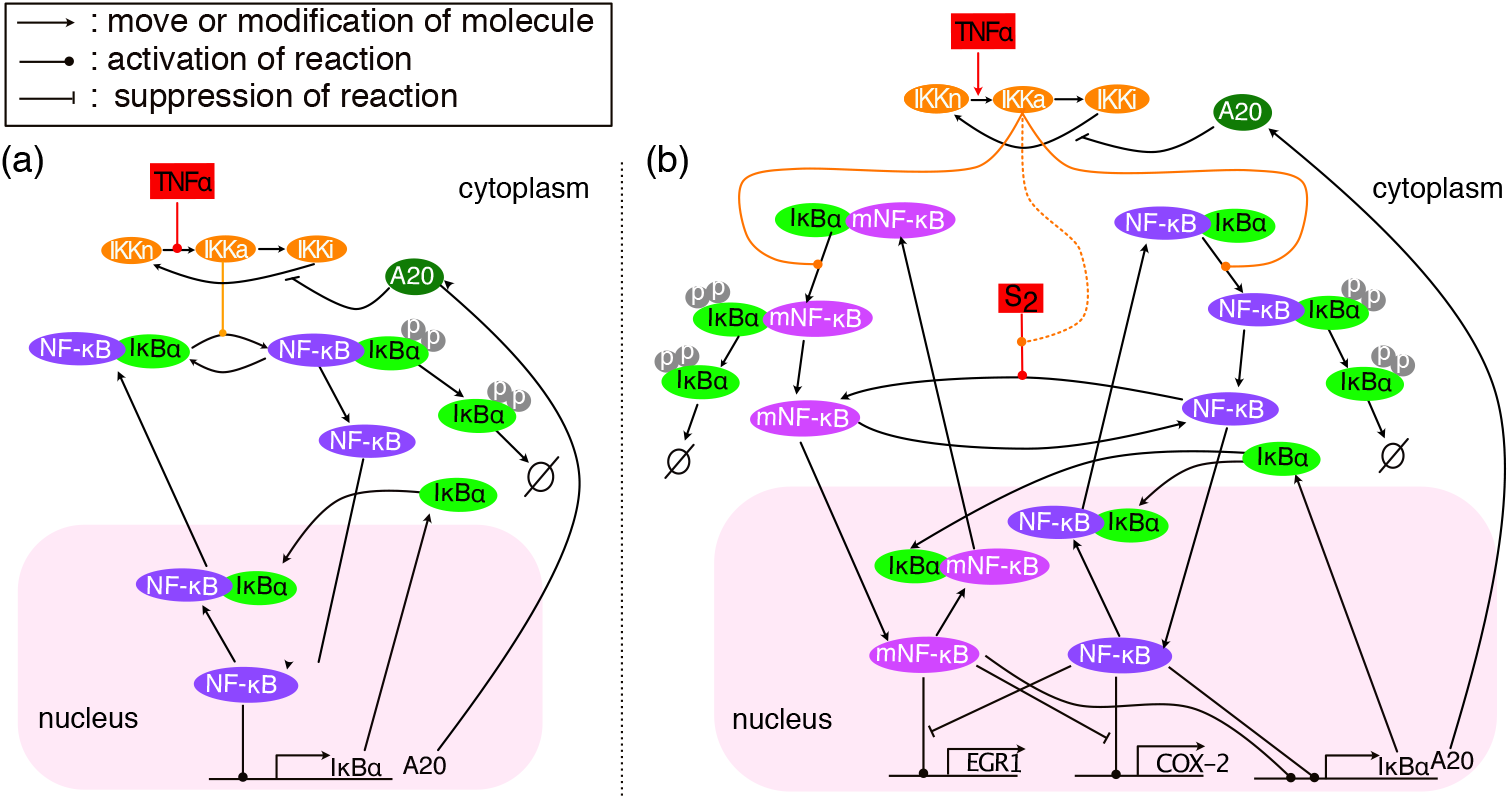
Diagrams of the base NF-*κ*B model in [1] and its modification mNF-*κ*B model. (a) The main reactions following a TNF*α* stimulus according to the model in [1] (base model); (b) The main reactions of the mNF-*κ*B model following a TNF*α* stimulus, and when the modification signal *S*_2_ is constantly transmitted. For the *m*_2_NF-*κ*B model, the *S*_2_ signal controls NF-*κ*B modification jointly (AND logic) with the TNF*α* signal through the active IKK molecules (dashed line).

We see in Fig. 4(a) that such a rapid decline in the eigenvalues of the FIM 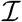 is the case for the base model considered here. They decay with an exponential rate and the second singular value is already less than 1% of the first one. But 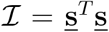 and therefore the eigenvalues of the FIM 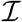 are the squares of the singular values *σ_i_* of **<u>s</u>** (SI Sect. 2.3). Consequently the singularvalues *σ_i_* of **<u>s</u>** rapidly decrease and, since *v*_1_ ⋯ *v_k_* ≤ *σ*_1_ ⋯ *σ_k_* for all *k* ≤ *s* with equality when *k* = *s* (Theorem 3.3.2 of [11], see SI section 1) the same is true for the multiplexing capacities 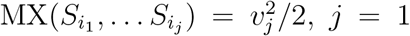, *j* = 1,…, *k*. Using equation (3) we see that the number of signals that can multiplex well must be very small. Fig. 4(b) shows how fast the multiplexing capacities decrease for our base model and identifies the parameters through which signal can multiplex more effectively i.e. these are the parameters that the signals should move if the signals are to be effective.

**Figure 4:**
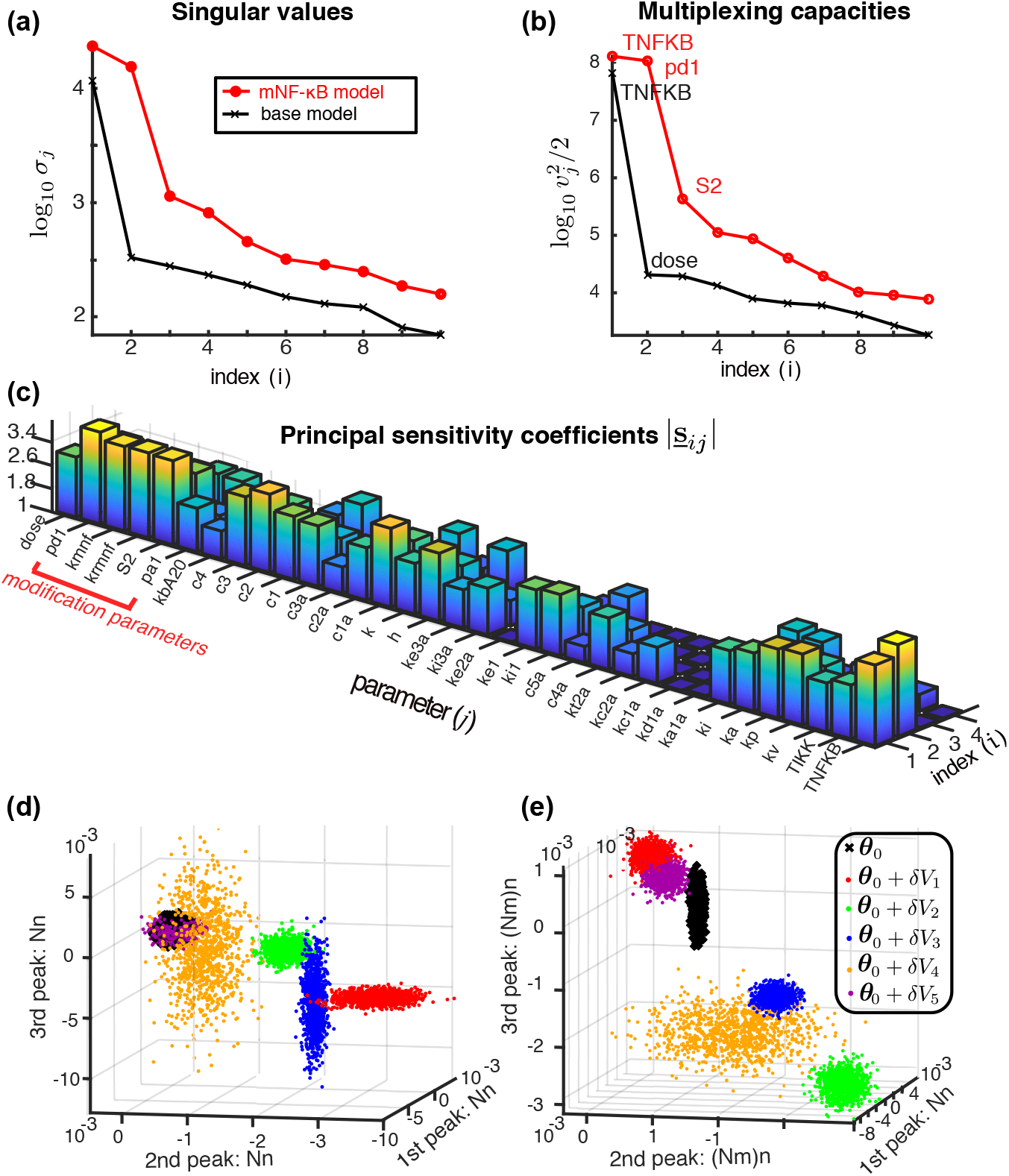
Comparisons of the multiplexing capacities and sensitivities of the base NF-*κ*B and mNF-*κ*B models. (a) The singular values *σ_i_* of the FIM for the base model and the much larger singular values for the mNF-*κ*B model. (b) The multiplexing capacities 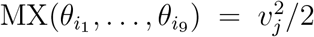 of the base and mNF-*κ*B models. The parameters with the largest multiplexing capacities correspond to the scaled version of the parameters TNFKB (total amount of NF-*κ*B molecules) and TNF*α* dose for the base model and TNFKB, *p*_*d*1_ (reverse modification rate of NF-*κ*B) and *S*_2_ (signal but treated as parameter here) for the mNF-*κ*B model. (c) The principal sensitivity coefficients of the mNF-*κ*B model. Larger values indicate higher sensitivity of the mNF-*κ*B model to changes in the value of the corresponding parameter. (d,e) Realisations (*n* = 1000) of the pcLNA distributions of the base and mNF-*κ*B model respectively at three times chosen to correspond to the first three peaks of the deterministic limit (Ω → ∞) of the model. In each case the 6 clusters correspond to different scaled parameter values which are either the scaled base value *θ*_0_ or the scaled parameter vector *θ*_0_ + *δV_j_*, *j* = 1,…, 5 (*δ* = 0.1) where *V*_1_,…, *V*_5_ are the eigendirections of the FIM corresponding to the 5 largest singular values of each model. Thus each cluster of points corresponds to a different principal component. Notice how much better the clusters are separated in the mNF-*κ*B model compared to the base NF-*κ*B model. For the mNF-*κ*B model the base and first three principal component perturbations (black, red, green and blue) are effectively completely separated.

### Increased multiplexing via additional regulated NF-*κ*B states

Regulation of the NF-*κ*B pathway is enabled by multiple post-translational modifications that control the activity of the core components of NF-*κ*B signaling. In particular, the RelA NF-*κ*B subunit undergoes reversible modifications such as phosphorylation, ubiquitination, and acetylation that can affect its transcriptional functions [20, 40, 41, 43, 44, 45, 46]. Indeed many modification sites in RelA have been identified as having either an enhancing, inhibitory, or modulatory effect on NF-*κ*B transcriptional activity in a gene-specific manner [20, 40, 41, 43, 45]. A further potentially regulated step that could differentially control individual gene expression is the hetero- and homo-dimerisation of the NF-*κ*B Rel proteins [48, 47]. Therefore, in considering the nature of biological mechanisms that could underlie multiplexing of information by the NF-*κ*B system, it is natural to consider modifications that create additional regulated NF-*κ*B states that can affect the transcription of NF-*κ*B target genes.

We consider one of the simplest modifications of the base model that can enable more effective multiplexing. In this modified model, which we call mNF-*κ*B, the cytoplasmic NF-*κ*B is reversibly modified by an input signal *S*_2_ that is independent to the TNFα signal. For example, *S*_2_ might be an environmental signal such as temperature or pressure that strongly affects the activity of a kinase or other molecular process. Such temperature effects have been studied extensively for the circadian clock in the context of temperature compensation [42] and more recently for the NF-*κ*B system [10]. Moreover, in Fig. 2 we see substantial temperature effects on the genes considered there.

The modified form of NF-*κ*B, mNF-*κ*B, competes with the unmodified form for binding of I*κ*B*α* but otherwise is subject to the same reactions (see Fig. 3(b) and SI Sect. 3.3). Importantly, mNF-*κ*B can activate, inhibit, or modulate the transcription of target genes and their differential expression can potentially reveal the levels of the *S*_2_ signal. The mathematical analysis of the stochastic version of the mNF-*κ*B model confirms this in the following ways.

Firstly, the singular values of the FIM are overall increased, and, importantly, there are now two large singular values rather than one (Fig. 4(a)). Secondly, the multiplexing capacities are increased substantially in the mNF-*κ*B model with three of then significantly above the second for the NF-*κ*B model. As explained above in the section ‘‘Multiplexing capacities of a model” and Sect. 2.5.1 of the SI this means that this model supports multiplexing of three signals through these three parameters. That the multiplexing capacities are large for the parameters related to the modification confirm that the extra sensitivity arises from the addition of this modification (see Fig. 4(d)).

Note that the results presented in Fig. 4 are derived for the probability distributions of stochastic trajectories of the system observed at 9 timepoints (see SI section 4.9). If instead only two time-points are considered, that is 10 mins before and at the expected time of the first peak of nuclear NF-*κ*B concentration, the base NF-*κ*B model is not largely affected, but the mNF-*κ*B presents a clearly less prominent increase of the singular values (see SI Fig. S8). This suggests that while the dynamical behaviour of the system does not in itself enable higher multiplexing capacity, it can greatly enhance multiplexing in a system that has the ability to multiplex.

The greater sensitivity of the mNF-*κ*B model compared to the base model is also reflected in Fig. 4(c). We see that the nuclear concentrations of NF-*κ*B are much more affected in the mNF-*κ*B model by changes in the signal. This clearly provides much greater ability in modulating gene expression according to different signals (see next section).

In this example we used one of the most simple and generic modifications where an external environmental signal such as temperature affects an internal parameter. In the SI (Sect. 3.5) we also consider another modification where the variation in the internal parameter is caused by a noisy pathway. Clearly, in this case the multiplexing capacity depends on the mechanism of this modification pathway and its information carrying effectiveness. Despite the increased noise levels, the multiplexing capacity of this alternative model can also be significantly larger than the base model.

### Reproducing multiplexing in the EGR1-COX-2 example

To further illustrate multiplexing, we now consider how to modify the signalling system so as to be able to reproduce the multiplexing behaviour seen in the EGR1-COX-2 gene expression data in Fig. 2. The same principle can be extended to reproduce the expression of the 8 genes presented in SI Fig. S10, but this is beyond our scope, and more data will be necessary for validation. Furthermore, we are not claiming that this is the true underlying biological mechanism but are using this example to illustrate how the NF-*κ*B signalling system can multiplex different signals through gene regulation. This is clearly not possible under the structural constraints of the base model because: (a) the base NF-*κ*B model reacts to pulses of TNF*α* stimuli by nearly identical (forced) oscillations and therefore it cannot explain the difference between early and late expression of EGR1 and COX-2, and (b) the differences in the base model between the response to short and long pulse are extremely small and can hardly explain the differences in EGR1 early response between the different pulse lengths.

The system is modified to include a reversible modification of NF-*κ*B molecules in the cytoplasm. The NF-*κ*B modification is jointly promoted by the TNF*α* stimulus through the IKK module and the independent signal *S*_2_ (see Fig. 3(b)). Pulses of TNF*α* cause bursts of NF-*κ*B nuclear translocations, but also higher levels of the modified NF-*κ*B. The reverse modification is independent of *S*_2_ and TNF*α*. Apart from TNF*α* promoting the NF-*κ*B modification, this model which we call m_2_NFKB is the same as the mNF-*κ*B model (see Fig. 3(b) and SI section 3). The m_2_NFKB model postulates that NF-*κ*B activates the transcription of EGR1, which is inhibited by the mNF-*κ*B, while the reverse regulation is imposed on COX-2. Using our approach to stochastic simulation outlined next, we can calculate the confidence limits for COX-2 and EGR1 under the various pulsing protocols (see Fig. 5) using *n* = 1000 trajectories simulated as described in the next section (see also SI section 4.6). Fig. 5(a) provides the mean time-trajectories (and 10 samples) at the same times observed using microarray and qPCR in Fig. 2. The introduction of the additional regulatory states of NF-*κ*B allows us to reproduce the experimentally observed profile.

**Figure 5:**
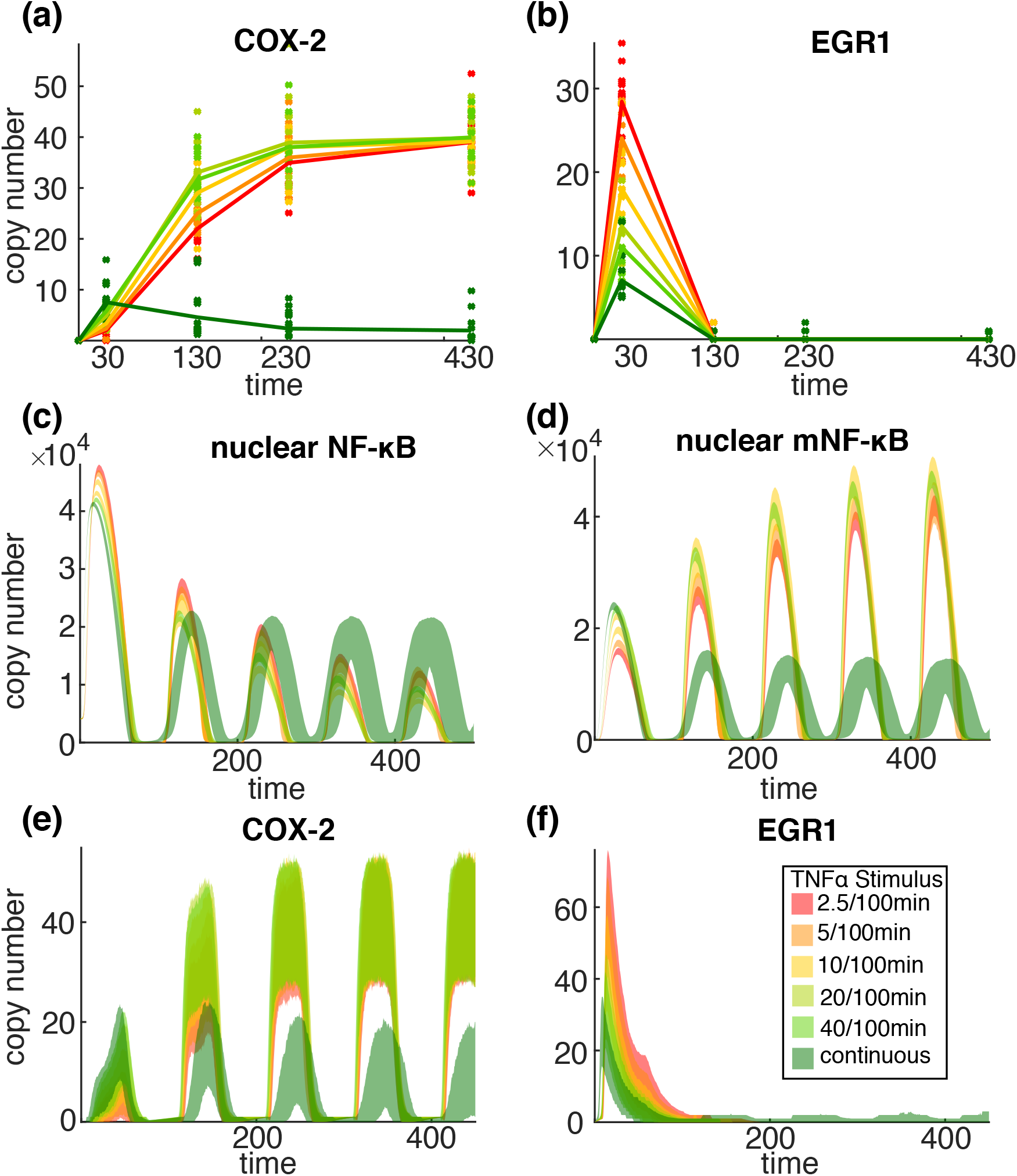
Stochastic simulation of the mNF-*κ*B model II describing the regulation of EGR1 and COX-2 genes for the same types of TNF*α* stimulation as in Fig. 2. (a) A sample of *n* =10 realisations (dots) and the sample mean (straight line) of simulated trajectories under the different TNF*α* stimuli (see legend) at the same times (*t* = 0, 30, 130, 230, 430*min*) as observations in Fig. 2; (b) 95% confidence envelopes of the copy number of (unbound) nuclear NF-*κ*B and mNF-*κ*B molecules, and EGR1, COX-2 mRNA copies under the different TNF*α* stimuli derived using stochastic simulations (n = 1000) under the different TNF*α* stimuli (see legend). The base model cannot reproduce the observed sensitivity to the different pulse lengths.

### Stochastic dynamics of NF-*κ*B

The base model used in our analysis is a stochastic reaction network that describes the oscillatory response of the NF-*κ*B system under stimulation by TNF*α*. It is a slight modification of the system model in [1]. In our version of the model, after adjustments to the rate equations, concentrations are all expressed in terms of the same volume Ω, taken to be Avogadro’s number in the appropriate molar units multiplied by the volume of the cell in appropriate units so that Ω has units L/nM (SI Sect. 3.3). The original model is written in terms of nuclear and cytoplasmic concentrations. Clearly, it is straightforward to convert between the two models (see SI Sect. 3.3).

We use the pcLNA stochastic version of this model [18] that allows us to derive analytical expressions for the FIM and system sensitivity matrix **<u>s</u>** and to rapidly simulate the system with high accuracy (see Fig. 6 and SI Section 4.2). The stochastic model considered here converges to the published deterministic model of [1] as Ω → ∞. We believe that the ability of our method to calculate important information-theoretic quantities such as these for a large fully stochastic model is a significant new development in itself.

**Figure 6:**
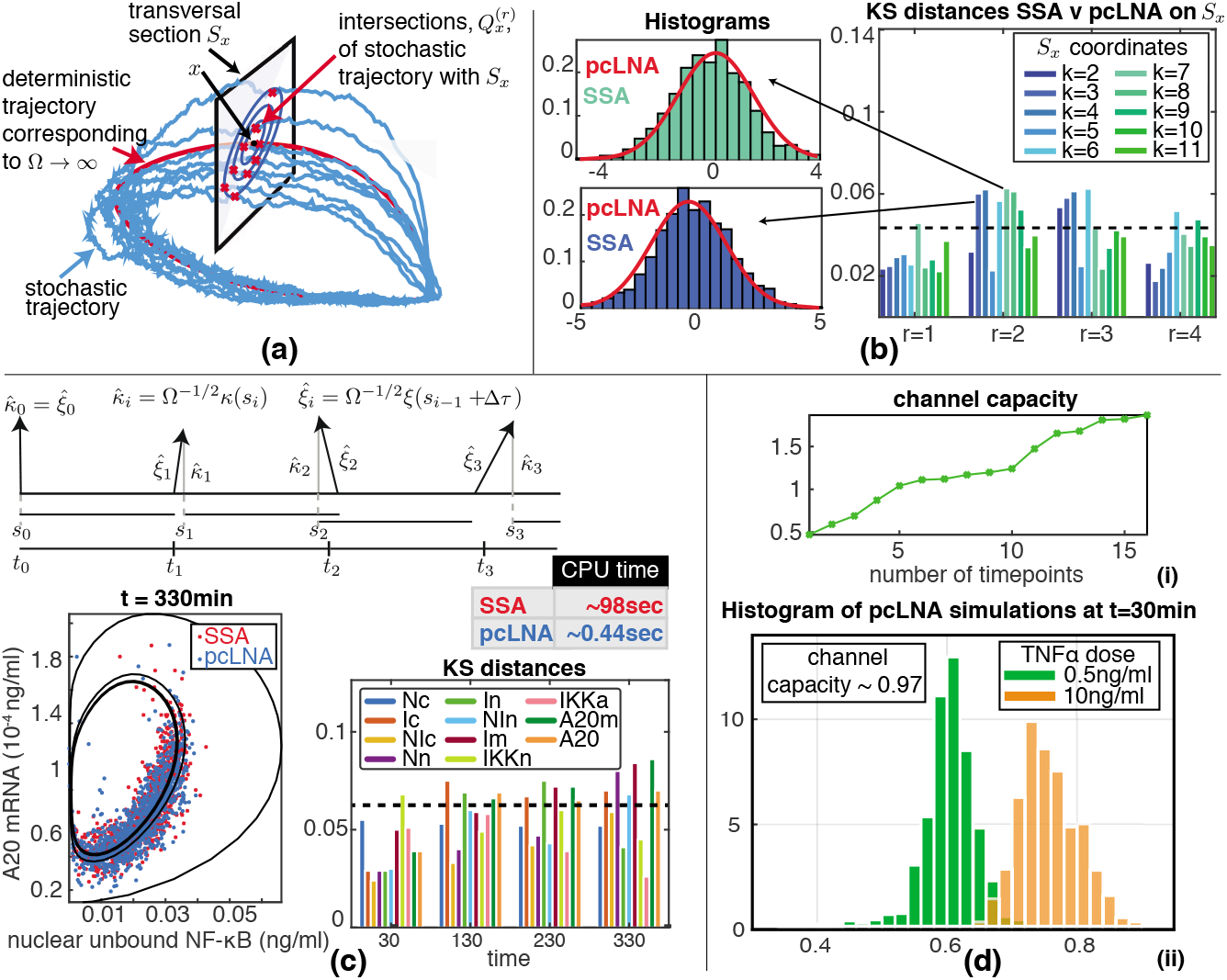
The pcLNA stochastic model and the channel capacity of the NF-*κ*B model. (a) The pcLNA model uses the stability of the probability distributions of stochastic oscillatory systems on the transversal sections, *S_x_*, of a given phase, *x*, of the system’s deterministic solution. (b) The pcLNA probability distributions on those transversal sections match very well the empirical distributions derived by SSA. Here the comparison is done using the Kolmogorov-Smirnov (KS) test at the first 4 peaks of NF-*κ*B model. The corresponding histograms for two of the largest observed KS values are also displayed to illustrate the nearly perfect match of the two distributions even in the case of the largest KS distances recorded here. (c) The pcLNA simulations also match very well the SSA simulations of the NF-*κ*B model which is much slower (see CPU (average) time for a single simulation). (d) Estimation of the channel capacity using the pcLNA simulation algorithm with added noise on the Total number of NF-*κ*B molecules.

The mNF-*κ*B model that includes NF-*κ*B modification is also simulated and analysed using pcLNA (see SI Section 3.4). For the simulation of downstream genes that are regulated by NF-*κ*B (see next section) we use the Stochastic Simulation Algorithm (SSA) [7]. This is because the relevant distribution for the gene expression is far from being Gaussian and therefore it is not appropriate to apply the pcLNA directly to this subsystem. Since this part of the system involves relatively few molecules the combined system can be simulated rapidly. The SSA is also used for comparisons to pcLNA in Fig. 6. In Fig. 6, we show that pcLNA accurately approximates the SSA simulations of the model in [1], which was calibrated to experimental data.

### Capacity of scalar channels

The results above raise the question of whether our models are compatible with the channel capacity seen in previous publications. We can use the base model to compare its behaviour with that discussed in [5, 28]. In these papers there was no attempt to control cell size or consideration of the total amount of NF-*κ*B (see also [37] which discusses such issues). We therefore allow these quantities to vary with the variation being drawn from a log-Normal distribution as described in SI Sect. 4.4.

We study the case where *S* is the level of the continuous TNF*α* stimulation (the parameter *dose*) and the response *R* is the level of nuclear NF-*κ*B at *q* different phases including its first peaks and troughs. Fig. 6(d)(i) shows the estimated capacity as a function of *q*. We also estimate the channel capacity for response *R* the nuclear concentration at *t* = 30*min* after initiating continuous TNF*α* stimulation (Fig. 6(d)(ii)).

The model reproduces the rather limited channel capacity seen in [5, 28] with estimated carrying capacities in the region of one bit. The exact value is not important because this is subject to our estimates of Ω, the total concentration of NF-*κ*B molecules and other parameters derived in [1]. A similar result can be obtained using the model in [31] (see SI section 4.5). It is worth emphasizing that the limited channel capacity is observed for a scalar signal and scalar response.

## Discussion

Cells present a very different context from that of traditional communications channels. The genetic and epigenetic information contained in the genome is translated by molecular interactions into dynamical processes. Described by dynamical interaction networks, these stochastic dynamical processes effectively move information from one system to another by regulating the probability distributions of their component molecules. Therefore, it is unclear whether the classical tools are always the most appropriate and it is likely that a much more extensive information toolbox is needed. New ideas about stochasticity and information are needed to understand how cells respond to dynamic environments so as to ensure appropriate cellular responses with high probability when they are using biochemistry that itself is very noisy.

Using such information theoretic tools we suggest a new insight into the way in which signalling systems transmit information. We mentioned above that recent research [5, 28] has shown that the channel TNF*α* level → nuclear NFKB abundance has a relatively low channel capacity. This raises the question of how our results fit with this. To some extent this is answered by the results in the section entitled “Capacity of scalar channels” where we show that our systems are tuned so as to reproduce this. Clearly, if we ignore noise and use a deterministic system we can make any such channel have as large a capacity as we want so it is important in our work to use reasonable levels of stochasticity.

We suggest that there is coherent picture emerging here where although signalling systems may be rather limited in the way that they transmit any scalar signal (as above for TNF*α* level), they are well designed to transmit multi-dimensional signals. There are two main reasons why when considering information flows in signalling one wishes to consider gene responses that are multidimensional. The first is that transmitting a signal via multiple receivers enables one to reduce the effects of noise. The second which is of central concern here is that it enables complex non-binary decisions. However, to make use of all these dimensions it is necessary that the input signal *S* has multiple dimensions because otherwise, if *R* is d-dimensional, the mean of *P*(*R|S*) is constrained to a 1*d* curve in d-dimensional space. This would mean that to obtain multiplexing or higher channel capacity one would have to use changes in the variance of *P*(*R|S*) with *S* to detect changes which seems very unlikely to be effective. Indeed, to use all d dimensions one needs dim *S* ≥ *d*.

We envisage that in this multidimensional situation it may well be the case that the scalar channels *S_i_* → *R* each have very low capacity as is the case in [5, 28] but that the full system *S* → *R* is able to multiplex so as to enable complex decisions and has a significantly higher capacity. Thus, by using multi-dimensionality the system can use multiple low-capacity components to produce a high capacity system.

A related issue concerns the role of dynamics in information transfer including the suggestion that dynamic systems such as oscillating ones can transmit greater amounts of information compared to static/equilibrium systems [1, 4, 14, 19, 22, 29, 36]. Our examples, also suggest why an oscillating system can use multiplexing to transmit more information than equilibrium systems. In these we see that signals that affect protein modification states or other aspects such as dimerization or binding partners can be good for multiplexing. In an equilibrium system the probability distribution describing how these states are distributed will be stationary in time. On the other hand in an oscillatory system these states can have a non-trivial temporal structure (e.g. oscillating) as catalysts of modifications can be activated and deactivated by interaction with the oscillations. This suggests a clear advantage for oscillating systems for information transfer.

## Acknowledgements

The work was supported by Biotechnology and Biological Sciences Research Council (BBSRC) grants BBF0059382/BB/F0059381/ BBF00561X1, BB/F005318/1, and BB/K003097/1. D.J.W. and D.A.R. were partly funded by the European Union Seventh Framework Programme (FP7/2007-2013) under Grant Agreement 305564. D.A.R. and G.M. were supported by Engineering and Science Research Council Grant EP/P019811/1.

## Supporting information

### Mathematical and computational details

Details of the mathematical analysis, description of the computational algorithms, and the models used, and additional figures.

