## Supplementary Information for "Multiplexing information flow through dynamic signalling systems"

### Supplementary Information for the paper "Multiplexing information flow through dynamic signalling systems"

April 28, 2020

**5 NF-$\kappa$B target genes** **31**

In the following, we provide technical details and various supplementary illustrations to those provided in the main paper that hereafter is referred as **I**.

#### 1 Mathematical details and notation

**Transpose of a vector or matrix and inner product** We denote $\boldsymbol{a}^T$ and $\boldsymbol{A}^T$ the transpose of the vector $\boldsymbol{a} \in \mathbb{R}^n$ and matrix $\boldsymbol{A} \in \mathbb{R}^{n \times n}$, respectively. We use the convention to write vectors as columns, i.e. $\boldsymbol{a} \in \mathbb{R}^{n \times 1}$, and therefore the transpose, $\boldsymbol{a}^T$, is a row vector, i.e. $\boldsymbol{a}^T \in \mathbb{R}^{1 \times n}$. The inner or dot product of two vectors $\boldsymbol{a}, \boldsymbol{b} \in \mathbb{R}^n$ is $\boldsymbol{a}^T\boldsymbol{b} = a_1b_1 + \cdots + a_nb_n$.

**Vectorisation of a matrix and the Kronecker product** The vectorisation operator $vec(\cdot)$ applied to an $m \times n$ matrix $A = (a_{ij})$, maps $A$ to an $mn \times 1$ vector, $vec(A) = (a_{11}, \ldots, a_{1n}, \ldots, a_{m1}, \ldots, a_{mn})^T$.

The Kronecker product of an $m \times n$ matrix $\boldsymbol{A} = (a_{ij})$ and an $s \times t$ matrix $B$ is the $ms \times nt$ matrix $\boldsymbol{A} \otimes \boldsymbol{B} = (a_{ij}\boldsymbol{B})_{ij}$. Then we have the following very useful equality: $\mathrm{vec}(\boldsymbol{ABC}) = (\boldsymbol{C}^T \otimes \boldsymbol{A})\mathrm{vec}(\boldsymbol{B})$ (see [7]).

**Probability density functions** For a stochastic process $\{\boldsymbol{X}(t), t \geq 0\}$ with continuous state space, we write the probability density function (pdf) at time $t \geq 0$ as $P(\boldsymbol{X}(t))$. The corresponding conditional pdf of $\boldsymbol{X}(t)$, $t > t_0$, given a fixed initial state $\boldsymbol{X}(t_0) = \boldsymbol{x}_0$ or an initial distribution $\boldsymbol{X}(t_0) \sim P_0$ are respectively denoted as $P(\boldsymbol{X}(t)|\boldsymbol{X}(t_0) = \boldsymbol{x}_0)$ and $P(\boldsymbol{X}(t)|\boldsymbol{X}(t_0) \sim P_0)$. The indicator $LNA$, as in $P_{\text{LNA}}$, is used in probability functions such as the above to denote that these are derived under the linear noise approximation (LNA).

**Multivariate Normal distribution** We write $\boldsymbol{\zeta} \sim MVN(\boldsymbol{m}, \boldsymbol{V})$ or $P$ is $MVN(\boldsymbol{m}, \boldsymbol{V})$ to mean that the random variable $\boldsymbol{\zeta}$ or the probability distribution $P$, respectively, is multivariate normal with mean vector $\boldsymbol{m}$ and covariance matrix $\boldsymbol{V}$.

**Singular Value Decomposition (SVD)** Singular Value Decomposition (SVD) gives a decomposition of the matrix $\boldsymbol{A} \in \mathcal{F}_{m \times k}$ into a product of the form $\boldsymbol{A} = \boldsymbol{UDV}^T$ where $\boldsymbol{U}$ is a $m \times k$ column-orthogonal matrix ($\boldsymbol{U}^T\boldsymbol{U} = \boldsymbol{I}_k$), $\boldsymbol{V}$ is a $k \times k$ orthogonal matrix ($\boldsymbol{V}^T\boldsymbol{V} = \boldsymbol{V}\boldsymbol{V}^T = \boldsymbol{I}_k$) and $\boldsymbol{D} = \mathrm{diag}(\sigma_1, \ldots, \sigma_k)$ is a diagonal matrix. This is the version of SVD that is often called thin SVD (see [14] for more details). The SVD applies for matrices defined in arbitrary fields $\mathcal{F}$, but henceforth we only consider real matrices, i.e. $\mathcal{F} = \mathbb{R}$. The ordered elements $\sigma_1 \geq \cdots \geq \sigma_k$ are the *singular values* of $\boldsymbol{A}$. We write the $i$th singular value $\sigma_i = \sigma_i(\boldsymbol{A})$. We note that the columns $\boldsymbol{V}_j$, $j = 1, \ldots k$ of $\boldsymbol{V}$ form an orthonormal basis for $\mathbb{R}^k$. If the last $n$ of the singular values are zero then the last $n$ columns are an orthonormal basis for the kernel of $\boldsymbol{A}$. If $m < k$ then $n \geq k - m$.

In what follows we will refer to the column vectors $\boldsymbol{V}_j = (V_{1j}, \ldots, V_{kj})^T$ and $\boldsymbol{U}_j = (U_{1j}, \ldots, U_{mj})^T$, $j = 1, 2, \ldots, k$ as the right and left singular vectors respectively, for which $\boldsymbol{A}\boldsymbol{V}_j = \sigma_j\boldsymbol{U}_j$ and $\boldsymbol{U}_j^T\boldsymbol{A} = \sigma_j\boldsymbol{V}_{j\cdot}^T$, with $\boldsymbol{V}_{j\cdot}^T$ the $j$th row of $\boldsymbol{V}$.

**QR-decomposition (QR)** The QR-decomposition with column pivoting produces a decomposition of any $m \times k$ matrix $\boldsymbol{M}$ as $\boldsymbol{MP} = \boldsymbol{QR}$ where $P$ is a permutation matrix, which is unitary ($\boldsymbol{P}^T\boldsymbol{P} = \boldsymbol{P}\boldsymbol{P}^T = \boldsymbol{I}$), $\boldsymbol{Q}$ is an $k \times k$ orthogonal matrix with orthonormal columns $\boldsymbol{Q}_j$, $j = 1, \ldots, k$, i.e. $\boldsymbol{Q}_j^T\boldsymbol{Q}_i = 0$, if $j \neq i$ and $\boldsymbol{Q}_i^T\boldsymbol{Q}_i = 1$, and $\boldsymbol{R}$ is an upper-triangular $m \times k$ matrix. If the row rank of $\boldsymbol{M}$ is $q$ then only the first $q$ of these entries are positive.

The permutation matrix $\boldsymbol{P}$ simply reorders the columns $\boldsymbol{M}_i$ of $\boldsymbol{M}$ so that the matrix $\boldsymbol{MP} = [\boldsymbol{M}_{i_1} \cdots \boldsymbol{M}_{i_k}]$ with the index set $\{i_1, \ldots, i_k\} = \{1, \ldots, k\}$. It is easy to check that $\boldsymbol{M}_{i_j} = \boldsymbol{Q}\boldsymbol{R}_j$ where $\boldsymbol{R}_j$ the $j$th column of $\boldsymbol{R}$. The entries $R_{ij}$ of $\boldsymbol{R}$ are $R_{ij} = \boldsymbol{Q}_i^T\boldsymbol{M}_{i_j}$, when

$i \leq j$, and 0 otherwise. The columns $\boldsymbol{Q}_i$ of $\boldsymbol{Q}$ are

$$\begin{aligned} \boldsymbol{U}_1 &= \boldsymbol{M}_{i_1}, & \boldsymbol{Q}_1 &= \boldsymbol{U}_1/\|\boldsymbol{U}_1\| \\ \boldsymbol{U}_2 &= \boldsymbol{M}_{i_2} - (\boldsymbol{M}_{i_2}^T\boldsymbol{Q}_1)\boldsymbol{Q}_1, & \boldsymbol{Q}_2 &= \boldsymbol{U}_2/\|\boldsymbol{U}_2\| \\ &\vdots & &\vdots \\ \boldsymbol{U}_k &= \boldsymbol{M}_{i_k} - \sum_{l=1}^{k-1}(\boldsymbol{M}_{i_k}^T\boldsymbol{Q}_l)\boldsymbol{Q}_l, & \boldsymbol{Q}_k &= \boldsymbol{U}_k/\|\boldsymbol{U}_k\| \end{aligned}$$

and thus

$$R_{jj}^2 = (\boldsymbol{M}_{i_j}^T\boldsymbol{Q}_j)^2 = \boldsymbol{M}_{i_j}^T\boldsymbol{M}_{i_j} - \sum_{l=1}^{j-1}(\boldsymbol{M}_{i_j}^T\boldsymbol{Q}_l)^2.$$

**Theorem 3.3.2 in [7]** Let $\boldsymbol{A} \in M_n$ have singular values $\sigma_1(\boldsymbol{A}) \geq \cdots \geq \sigma_n(\boldsymbol{A}) \geq 0$ and eigenvalues $\{\lambda_1(\boldsymbol{A}), \dots, \lambda_n(\boldsymbol{A})\} \subset \mathbb{C}$ ordered so that $|\lambda_1(\boldsymbol{A})| \geq \cdots \geq |\lambda_n(\boldsymbol{A})|$. Then

$$|\lambda_1(\boldsymbol{A}) \cdots \lambda_k(\boldsymbol{A})| \leq \sigma_1(\boldsymbol{A}) \cdots \sigma_k(\boldsymbol{A}) \quad \text{for } k = 1, \dots, n, \text{ with equality for } k = n.$$

#### 2 Fisher Information, Sensitivity Matrix & Multiplexing Capacity

We next outline an approach to sensitivity analysis and multiplexing. The key point will be that the sensitivity matrix $\underline{\mathbf{s}}$, previously introduced in [13], can be used to characterise the ability of the system to multiplex. We consider the stochastic response $\boldsymbol{R} = (R_1, \dots, R_n)^T$, that has probability distribution $P_{\boldsymbol{S}}$ which depends on the signal $\boldsymbol{S} = (S_1, S_2, \dots, S_s)^T \in \mathcal{S}$.

##### 2.1 Fisher Information Matrix & KL Divergence

Let $\ell(\boldsymbol{S}; r) = \log p_{\boldsymbol{S}}(r)$ be the log-likelihood function of $P_{\boldsymbol{S}}$ that has pdf $p_{\boldsymbol{S}}(r)$, $r \in \mathcal{R}$. The Fisher Information Matrix (FIM), $\mathcal{I}(\boldsymbol{S})$, of the probability distribution $P_{\boldsymbol{S}}$ at $\boldsymbol{S} \in \mathcal{S}$ is a symmetric positive-(semi)definite $s \times s$ matrix with entries

$$\mathcal{I}_{ij}(\boldsymbol{S}) = \mathbb{E}_{P_{\boldsymbol{S}}}\left(\partial_i \ell \cdot \partial_j \ell\right) = -\mathbb{E}_{P_{\boldsymbol{S}}}\left(\partial_{ij}^2 \ell\right)$$

Here $\mathbb{E}_{P_{\boldsymbol{S}}}$ denotes the expectation function under the distribution $P_{\boldsymbol{S}}$, $\partial_i$ the partial derivative with respect to the $i$th component $S_i$ evaluated at $\boldsymbol{S} \in \mathcal{S}$ and $\partial_{ij}^2$ the corresponding second order derivative.

If the signals $S_i$, $i = 1, \dots, s$, act through changing parameters $\theta_j$ of the system (i.e. so that $\boldsymbol{\theta} = \boldsymbol{\theta}(\boldsymbol{S})$) and $\mathcal{I}_\theta$ the FIM with respect to $\boldsymbol{\theta} = (\theta_1, \dots, \theta_m)'$ we have that

$$\mathcal{I}(\boldsymbol{S}) = d\boldsymbol{\theta}(\boldsymbol{S})^T \mathcal{I}_\theta d\boldsymbol{\theta}(\boldsymbol{S}) \tag{1}$$

where $d\boldsymbol{\theta}(\boldsymbol{S})$ is the derivative of $\boldsymbol{\theta}$ with respect to $\boldsymbol{S}$ at $\boldsymbol{S} \in \mathcal{S}$ . This well-known property of the FIM follows from the chain rule for derivatives.

The Kullback–Leibler (KL) divergence of two probability distributions $P$ and $Q$ with density functions $p(x)$ and $q(x)$, $x \in \mathcal{X}$, is given by

$$D_{\mathrm{KL}}(P\|Q) = \int_{\mathcal{X}} p(x) \log \frac{p(x)}{q(x)}\, dx.$$

That is, the KL divergence $D_{\mathrm{KL}}(P\|Q)$ is the expected value of the likelihood ratio $\lambda(x) = \log p(x)/q(x)$ with the expectation taken with respect to $P$ with the usual conventions when $p(x) = 0$ or $q(x) = 0$. If $P = P_{\boldsymbol{S}+\delta\boldsymbol{S}}$ and $Q = P_{\boldsymbol{S}}$, then (see [3]),

$$D_{\mathrm{KL}}(P_{\boldsymbol{S}+\delta\boldsymbol{S}}\|P_{\boldsymbol{S}}) = \frac{1}{2}\delta\boldsymbol{S}^T\mathcal{I}(\boldsymbol{S})\delta\boldsymbol{S} + \mathrm{O}(\|\delta\boldsymbol{S}\|^3).$$

That is, the FIM is the hessian matrix of the above KL divergence at $\boldsymbol{S} \in \mathcal{S}$.

##### 2.2 Information geometry

If the Fisher information matrix $\mathcal{I}(\boldsymbol{S}_0)$, $\boldsymbol{S}_0 \in \mathcal{S}$, is positive definite, it can also be used to define a Riemannian metric over the space of probability distributions. Firstly, we use it to define a metric on the signal space $\mathcal{S}$ at $\boldsymbol{S}_0$ by

$$\langle \delta\boldsymbol{S}, \delta\boldsymbol{S}'\rangle_{\boldsymbol{S}_0} = \sum_{i,j} \delta\boldsymbol{S}_i \delta\boldsymbol{S}'_j \mathcal{I}_{ij}(\boldsymbol{S}_0) = \delta\boldsymbol{S}^T \mathcal{I}(\boldsymbol{S}_0)\delta\boldsymbol{S}'.$$

This inner product is interesting to us because,

$$D_{\mathrm{KL}}(P_{\boldsymbol{S}_0+\delta\boldsymbol{S}}||P_{\boldsymbol{S}_0}) = \frac{1}{2}\langle \delta\boldsymbol{S}, \delta\boldsymbol{S}\rangle_{\boldsymbol{S}_0} + \mathrm{O}(||\delta\boldsymbol{S}||^3).$$

Therefore, assuming that $\mathcal{I}(\boldsymbol{S})$ depends in a $\mathrm{C}^1$ fashion on $\boldsymbol{S}$, if $\boldsymbol{S}$ and $\boldsymbol{S}'$ are perturbations of $\boldsymbol{S}_0$

$$D_{\mathrm{KL}}(P_{\boldsymbol{S}}||P_{\boldsymbol{S}'}) = \frac{1}{2}\langle \boldsymbol{S}-\boldsymbol{S}', \boldsymbol{S}-\boldsymbol{S}'\rangle_{\boldsymbol{S}_0} + \mathrm{O}(\max\{\|\boldsymbol{S}-\boldsymbol{S}'\|^3, \|\boldsymbol{S}-\boldsymbol{S}_0\|^3, \|\boldsymbol{S}'-\boldsymbol{S}_0\|^3\}. \quad (2)$$

If this is the case we say $D_{\mathrm{KL}}(P_{\boldsymbol{S}}||P_{\boldsymbol{S}'}) = \frac{1}{2}\langle \boldsymbol{S}-\boldsymbol{S}', \boldsymbol{S}-\boldsymbol{S}'\rangle_{\boldsymbol{S}}$ up to third order.

##### 2.3 The multiplexing sensitivity matrix $\underline{\mathbf{s}}$

Since the FIM $\mathcal{I} = \mathcal{I}(\boldsymbol{S}_0)$ is symmetric and positive semi-definite its SVD is of the form $\boldsymbol{V}\boldsymbol{D}\boldsymbol{V}^T$ where $\boldsymbol{V}$ is orthogonal and $\boldsymbol{D}$ is diagonal with entries $\sigma_1^2 \geq \cdots \geq \sigma_s^2 \geq 0$. The *sensitivity matrix* $\underline{\mathbf{s}}$ associated with $\mathcal{I}$ is $\underline{\mathbf{s}} = \boldsymbol{D}^{1/2}\boldsymbol{V}^T$. We explain the reason for this name below.

##### 2.4 Multiplexing Capacity

Recall the definition of $D_{\mathrm{KL}}^{(i,\boldsymbol{S}_0)}$ as $\min_{\boldsymbol{S}} \eta_i^{-2} \min_{\boldsymbol{S}'} D_{\mathrm{KL}}(P_{\boldsymbol{S}}||P_{\boldsymbol{S}'})$ where the $i$th component of $\boldsymbol{S}-\boldsymbol{S}_0$ has magnitude $\eta_i = |S_i - S_{0,i}| > 0$ and the minimum is taken over all perturbations $\boldsymbol{S}'$ of $\boldsymbol{S}_0$ where the $i$th component $S_i$ has not changed, i.e. $S'_i - S_{0,i} = 0$. Note that $D_{\mathrm{KL}}^{(i,\boldsymbol{S}_0)}$ depends upon the set of signals $(S_1, \ldots, S_k)$, $k \leq s$, being considered. If we need to stress this we will write $D_{\mathrm{KL}}^{(i,\boldsymbol{S}_0)}(S_1, \ldots, S_k)$ instead. Note that in general $D_{\mathrm{KL}}^{(i,\boldsymbol{S}_0)}(S_1, \ldots, S_k) \neq D_{\mathrm{KL}}^{(i,\boldsymbol{S}_0)}(S_1, \ldots, S_{k-1})$.

Given any set of integers $1 \leq i_1 < \cdots < i_r < s$ distinct from $i$ consider the columns $\underline{\mathbf{s}}_{i_j}$ of $\underline{\mathbf{s}}$. Denote by $n(\underline{\mathbf{s}}_i|\underline{\mathbf{s}}_{i_1}, \ldots, \underline{\mathbf{s}}_{i_r}) = \mathbf{n}(i|i_1, \ldots, i_r)$ the unique vector $\underline{\mathbf{n}}$ normal to the vectors $\underline{\mathbf{s}}_{i_1}, \ldots, \underline{\mathbf{s}}_{i_r}$ such that $\underline{\mathbf{s}}_i - \underline{\mathbf{n}}$ lies in the span of $\underline{\mathbf{s}}_{i_1}, \ldots, \underline{\mathbf{s}}_{i_r}$. That is,

$$\underline{\mathbf{n}}^T\underline{\mathbf{s}}_{i_j} = 0, \quad j = 1, \ldots, r, \quad \text{and} \quad \underline{\mathbf{n}}^T(\underline{\mathbf{s}}_i - \underline{\mathbf{n}}) = 0. \quad (3)$$

In the case where $1 \leq i_1 < \cdots < i_r < s$ consists of all $j$ with $j \neq i$ we use the notation $n(\underline{\mathbf{s}}_i|\underline{\mathbf{s}}_j, j \neq i)$. We now prove that, up to third order terms,

$$D_{\mathrm{KL}}^{(i,\boldsymbol{S}_0)} = \frac{1}{2}||n(\underline{\mathbf{s}}_i|\underline{\mathbf{s}}_j, j \neq i)||^2. \quad (4)$$

We first prove (4) for the case where $i = s$. Let $\underline{\mathbf{s}} = \boldsymbol{Q}\boldsymbol{R}$ be the $QR$ decomposition of $\underline{\mathbf{s}}$. By definition $\underline{\mathbf{n}} = n(\underline{\mathbf{s}}_s|\underline{\mathbf{s}}_j, j \neq s) = R_{ss}\boldsymbol{Q}_s$ and therefore $\|\underline{\mathbf{n}}\| = |R_{ss}|$. To check this, one can write $\underline{\mathbf{n}} = \sum_i c_i\boldsymbol{Q}_i$ and use (3) to derive the coefficients $c_j = 0$, $j < s$ and $c_s = R_{ss}$.

Then, see that $\mathcal{I}(\boldsymbol{S}_0) = \underline{\mathbf{s}}^T\underline{\mathbf{s}} = \boldsymbol{R}^T\boldsymbol{R} = \boldsymbol{V}\boldsymbol{D}\boldsymbol{V}^T$ since $\boldsymbol{Q}^T\boldsymbol{Q}$ is the identity matrix. But, by (2), up to third order,

$$2D_{\mathrm{KL}}(P_{\boldsymbol{S}}||P_{\boldsymbol{S}'}) = \langle \delta\boldsymbol{S}, \delta\boldsymbol{S}\rangle_{\boldsymbol{S}_0} = \delta\boldsymbol{S}^T\mathcal{I}(\boldsymbol{S}_0)\delta\boldsymbol{S} = \delta\boldsymbol{S}^T\boldsymbol{R}^T\boldsymbol{R}\delta\boldsymbol{S} = \|\boldsymbol{R}\delta\boldsymbol{S}\|^2. \quad (5)$$

where $\delta \boldsymbol{S} = \boldsymbol{S} - \boldsymbol{S}'$. Because $\boldsymbol{R}$ is upper triangular and $\delta \boldsymbol{S}_s = S_s$ the last entry of the vector $\boldsymbol{R}\delta\boldsymbol{S}$ is $R_{ss}S_s$. It follows that, up to third order terms,

$$2D_{\mathrm{KL}}(P_{\boldsymbol{S}}||P_{\boldsymbol{S}'}) = R_{ss}^2 \boldsymbol{S}_s^2 + \sum_{j=1}^{s-1} (\boldsymbol{R}\delta\boldsymbol{S})_j^2 \geq |R_{ss}\boldsymbol{S}_s|^2 = \|\underline{\mathbf{n}}\|^2 \eta_s^2,$$

where $(\boldsymbol{R}\delta\boldsymbol{S})_j$ the $j$th entry of the vector $\boldsymbol{R}\delta\boldsymbol{S}$ and the minimum is achieved when $S_j'$ is such that $(\boldsymbol{R}\delta\boldsymbol{S})_j = 0$, for $j = 1, \ldots, s-1$. These $S_j'$'s can be found if we let $\boldsymbol{R}'$ be the $(s-1) \times (s-1)$ upper triangular matrix obtained by deleting the last row and column from $\boldsymbol{R}$ and let $\boldsymbol{R}_s$ be the $(s-1)$-dimensional vector obtained by deleting the last element from the last column of $\boldsymbol{R}$. Then, if $\delta\boldsymbol{S}'$ is the $(s-1)$-dimensional vector solving $\boldsymbol{R}'\delta\boldsymbol{S}' = -\boldsymbol{R}_k$ and $\delta\boldsymbol{S}$ is the $s$-dimensional vector obtained by adding $\delta\boldsymbol{S}_s$ to the end of $\delta\boldsymbol{S}'$ we have that $\|R\delta\boldsymbol{S}\|^2 = |\delta\boldsymbol{S}_s|^2$. Therefore, up to third order terms, $2D_{\mathrm{KL}}^{(i,\boldsymbol{S}_0)} = R_{ss}^2$.

In case $i < s$, we can simply use a permutation matrix, $\boldsymbol{P}$, such that the last column of the matrix $\underline{\mathbf{s}}\boldsymbol{P}$ is $\underline{\mathbf{s}}_i$, and follow the same steps as above to show that, up to third order terms, $2D_{\mathrm{KL}}^{(i,\boldsymbol{S}_0)} = ||n(\underline{\mathbf{s}}_i|\underline{\mathbf{s}}_j, j \neq i)||^2 = R_{ss}^2$, where $R_{ss}$ the bottom right entry of the $\boldsymbol{R}$ matrix that satisfies $\underline{\mathbf{s}}\boldsymbol{P} = \boldsymbol{Q}\boldsymbol{R}$. This proves (4).

##### 2.4.1 Finding optimal multiplexing sets

In view of the above result we ask the following question, given a set of signals $S_1, \ldots, S_s$, for which subsets $S_{i_1}, \ldots, S_{i_r}$, $r \leq s$, can we effectively multiplex in the sense that all the multiplexing capacities for $S_{i_1}, \ldots, S_{i_r}$ are above a given threshold.

We first choose the column pivoting matrix $\boldsymbol{P}^*$ so that the diagonal entries, $R_{jj}$, of the $\boldsymbol{R}$ matrix in the QR decomposition $\underline{\mathbf{s}}\boldsymbol{P}^* = [\underline{\mathbf{s}}_{i_1} \cdots \underline{\mathbf{s}}_{i_s}] = \boldsymbol{Q}\boldsymbol{R}$ give the minimum multiplexing capacities $\mathrm{MX}(S_{i_1}, \ldots, S_{i_k})$. That is, up to third order,

$$\mathrm{MX}(S_{i_1}, \ldots, S_{i_k}) = \min_{i=i_1,\ldots,i_k} D_{\mathrm{KL}}^{(i,\boldsymbol{S}_0)}(S_{i_1}, \ldots, S_{i_k}) = R_{kk}^2/2, \quad k = 1, \ldots, s. \tag{6}$$

This QR decomposition can be easily derived by an algorithm such as this provided in Table S1. The idea is to start with the full set of signals and use QR to find and remove from the current set of signals, say $\mathcal{S}_k = \{S_{i_1}, \ldots, S_{i_k}\}$, $k = s, s-1, \ldots, 2$, the signal that gives the minimum multiplexing capacity $\min_{i_1,\ldots,i_l} \|n(\underline{\mathbf{s}}_i|\underline{\mathbf{s}}_{i_l} \in \mathcal{S}_k \setminus \underline{\mathbf{s}}_i)\|^2$. These multiplexing capacities can be derived by the following steps: (i) pivoting through an appropriate $\boldsymbol{P}$ the $\underline{\mathbf{s}}_{i_l}$ columns, $l = 1, \ldots, k$, to get $\underline{\mathbf{s}}\boldsymbol{P}$ with last column equal to $\underline{\mathbf{s}}_i$, (ii) performing QR decomposition to get $\underline{\mathbf{s}}\boldsymbol{P} = \boldsymbol{Q}\boldsymbol{R}$ and (iii) using that $R_{kk}^2 = \|n(\underline{\mathbf{s}}_i|\underline{\mathbf{s}}_{i_l} \in \mathcal{S}_k \setminus \underline{\mathbf{s}}_i)\|^2$ to compute the minimum $D_{\mathrm{KL}}$ (see Table S1). Note that this is a backward reduction QR decomposition as opposed to the standard (forward addition) column pivoting QR decomposition provided by software such as MATLAB [11].

It is easy to see that, as in the standard column pivoting QR decomposition, the diagonal entries $R_{ii}$ of $\boldsymbol{R}$ in $\underline{\mathbf{s}}\boldsymbol{P}^* = \boldsymbol{Q}\boldsymbol{R}$ are in decreasing order

$$R_{11}^2 \geq \cdots \geq R_{ss}^2.$$

This is because, for $l \leq s$,

$$|R_{ll}| = \|n(\underline{\mathbf{s}}_{i_l}|\underline{\mathbf{s}}_1, \ldots, \underline{\mathbf{s}}_{i_{l-1}})\| \leq \|n(\underline{\mathbf{s}}_{i_{l-1}}|\underline{\mathbf{s}}_1, \ldots, \underline{\mathbf{s}}_{i_{l-2}}, \underline{\mathbf{s}}_{i_l})\| \leq \|n(\underline{\mathbf{s}}_{i_{l-1}}|\underline{\mathbf{s}}_{i_1}, \ldots, \underline{\mathbf{s}}_{i_{l-2}})\| = |R_{l-1,l-1}|.$$

However, the backward reduction QR, unlike the forward addition QR, ensures that the diagonal entries of $\boldsymbol{R}$ can be used to derive the minimum multiplexing capacities as in (6).

Once the appropriate ordering of signals $S_{i_1}, \ldots, S_{i_s}$ is identified by the backward reduction QR, we can perform a QR decomposition without pivoting to the sensitivity matrix $\underline{\mathbf{s}}^* = [\underline{\mathbf{s}}_{i_1} \ldots \underline{\mathbf{s}}_{i_s}] = \boldsymbol{Q}\boldsymbol{R}$ and by examining the diagonal entries of $\boldsymbol{R}$, we can see which signals $S_{i_1}, S_{i_2}, \dots, S_{i_k}$, $k = 1, \dots, s$ can multiplex before the multiplexing capacity $\mathrm{MX}(S_{i_1}, S_{i_2}, \dots, S_{i_k}) = R^2_{kk}/2$ falls below a threshold. Equivalently, one can start from the full set of signals $S_1, \dots, S_s$ and find the minimum subset of signals, $S_{i_{s-r+1}}, \dots, S_{i_s}$, $r \geq 0$, that needs to be removed (i.e. keep unperturbed) to derive a multiplexing subset, i.e. a subset that has $\mathrm{MX}(S_{i_1}, S_{i_2}, \dots, S_{i_{s-r}}) = R^2_{s-r,s-r}/2$ larger than a threshold.

| |
|---|
| 1. Compute the SVD, $\mathcal{I} = VD\boldsymbol{V}^T$, of the FIM $\mathcal{I}$ and set $\underline{\mathbf{s}} = D^{1/2}\boldsymbol{V}^T$ and $s' = s$ |
| 2. For $i = 1, \dots, s'$: |
| (a) Use a column pivoting matrix $P$ to get $\underline{\mathbf{s}}' = \underline{\mathbf{s}}P$ so that its last column $\underline{\mathbf{s}}'_{s'} = \underline{\mathbf{s}}_i$, where $\underline{\mathbf{s}}_i$ the $i$th column of $\underline{\mathbf{s}}$. |
| (b) Perform QR decomposition to compute $QR = \underline{\mathbf{s}}'$ |
| (c) Set $r_i = \lvert R_{s's'} \rvert$ |
| 3. Find $i_{s'} = \arg\min_{i=1,\dots,s'} r_i$ |
| 4. Set $s' := s' - 1$ and $\underline{\mathbf{s}} := [\underline{\mathbf{s}}_1 \cdots \underline{\mathbf{s}}_{i_{s'}-1}\, \underline{\mathbf{s}}_{i_{s'}+1} \cdots \underline{\mathbf{s}}_s]$ |
| 5. If $s' > 1$ return to step 2. |

**Table S1:** *The backward reduction QR decomposition used in section 2.4.1.*

#### 2.5 Sensitivity analysis for multivariate normal distributions

In this section we assume that $P_{\boldsymbol{S}}$ is $\mathrm{MVN}(\boldsymbol{\mu}, \boldsymbol{\Sigma})$, with mean $\boldsymbol{\mu} = \boldsymbol{\mu}(S)$, and covariance $\Sigma = \Sigma(S)$. Then the entries of the FIM are

$$\mathcal{I}_{ij}(\boldsymbol{S}_0) = (\partial_i \boldsymbol{\mu})^T \boldsymbol{\Sigma}^{-1} (\partial_j \boldsymbol{\mu}) + \frac{1}{2} \mathrm{tr}(\boldsymbol{\Sigma}^{-1} (\partial_i \boldsymbol{\Sigma}) \boldsymbol{\Sigma}^{-1} (\partial_j \boldsymbol{\Sigma})) \tag{7}$$

where all derivatives are taken at $\boldsymbol{S} = \boldsymbol{S}_0$. This can also be written using vec notation as

$$\mathcal{I}_{ij}(\boldsymbol{S}_0) = (\partial_i \boldsymbol{\mu})^T \boldsymbol{\Sigma}^{-1} (\partial_j \boldsymbol{\mu}) + \frac{1}{2} vec(\partial_i \boldsymbol{\Sigma})(\boldsymbol{\Sigma}^{-1} \otimes \boldsymbol{I})(\boldsymbol{I} \otimes \boldsymbol{\Sigma}^{-1}) vec(\partial_j \boldsymbol{\Sigma}).$$

Now consider the $N \times s$ matrix ($N = n + n^2$)

$$\mathcal{L} = \begin{pmatrix} \partial \boldsymbol{\mu} \\ \partial vec(\boldsymbol{\Sigma}) \end{pmatrix} = \begin{pmatrix} \partial_1 \boldsymbol{\mu} & \dots & \partial_k \boldsymbol{\mu} \\ \partial_1 vec(\boldsymbol{\Sigma}) & \dots & \partial_k vec(\boldsymbol{\Sigma}) \end{pmatrix}, \tag{8}$$

which is the derivative (linearisation matrix) of the mapping $\boldsymbol{S} \mapsto \Theta(\boldsymbol{S}) = (\boldsymbol{\mu}(\boldsymbol{S}), \mathrm{vec}\boldsymbol{\Sigma}(S))$ at $\boldsymbol{S}_0$. That is, if we let $\delta\boldsymbol{\mu} = (\delta\mu_1, \dots, \delta\mu_n)^T$ and $\delta vec(\Sigma) = (\delta\Sigma_{11}, \dots, \delta\Sigma_{nn})^T$, with $\delta\mu_i = \mu_i(\boldsymbol{S} + \delta\boldsymbol{S}) - \mu_i(\boldsymbol{S})$ and $\delta\Sigma_{ij} = \Sigma_{ij}(\boldsymbol{S} + \delta\boldsymbol{S}) - \Sigma_{ij}(\boldsymbol{S})$, then

$$\delta\Theta = (\delta\boldsymbol{\mu}, \delta\mathrm{vec}(\boldsymbol{\Sigma}))^T = \mathcal{L}\, \delta\boldsymbol{S} + \mathrm{O}(\lVert \delta\boldsymbol{S} \rVert^2). \tag{9}$$

If we also define the $N \times N$ matrix $\mathcal{F}$ as the Cholesky decomposition of the positive-definite matrix $\boldsymbol{F}$ (i.e. $\mathcal{F}^T\mathcal{F} = \boldsymbol{F}$), where

$$\boldsymbol{F} = \begin{pmatrix} \boldsymbol{\Sigma}^{-1} & 0 \\ 0 & (\boldsymbol{\Sigma}^{-1} \otimes \boldsymbol{I})(\boldsymbol{I} \otimes \boldsymbol{\Sigma}^{-1})/2 \end{pmatrix}$$

then we can write the Fisher information in (7) as

$$\mathcal{I}(\boldsymbol{S}_0) = (\mathcal{F}\mathcal{L})^T (\mathcal{F}\mathcal{L}).$$

Therefore, $\mathcal{F}\mathcal{L}$ is a linear map from $\mathcal{S}$ to $\mathbb{R}^N$ which sends the $\langle \cdot, \cdot \rangle_{\boldsymbol{S}_0}$ metric to the standard one in $\mathbb{R}^N$:

$$\langle \delta\boldsymbol{S}, \delta\boldsymbol{S}' \rangle_{\boldsymbol{S}_0} = \delta\boldsymbol{S}^T \mathcal{I}(\boldsymbol{S}_0) \delta\boldsymbol{S}' = (\mathcal{F}\mathcal{L}\delta\boldsymbol{S})^T (\mathcal{F}\mathcal{L}\delta\boldsymbol{S}') = \delta\boldsymbol{S}^T (\mathcal{F}\mathcal{L})^T (\mathcal{F}\mathcal{L}) \delta\boldsymbol{S}'$$

and relates the FI metric in $\mathcal{S}$ to the standard one in $\mathbb{R}^{n+n^2}$:

$$\lVert \delta\boldsymbol{S} \rVert^2_S = \lVert \mathcal{F}\mathcal{L}\delta\boldsymbol{S} \rVert^2.$$

##### 2.5.1 The matrix $\underline{\mathbf{s}}$ characterises sensitivity

The sensitivity to changes $\delta \boldsymbol{S}$ in $S$ of the probability distribution $MVN(\boldsymbol{\mu}(\boldsymbol{S}), \boldsymbol{\Sigma}(\boldsymbol{S}))$ can therefore be studied using the vector $\mathcal{F}\mathcal{L}\,\delta\boldsymbol{S}$. Equation (9) shows that

$$\mathcal{F}\,\delta\Theta = \mathcal{F}(\delta\boldsymbol{\mu}, \delta\mathrm{vec}(\boldsymbol{\Sigma}))^T = \mathcal{F}\mathcal{L}\delta\boldsymbol{S} + \mathrm{O}(\|\delta\boldsymbol{S}\|^2).$$

We now consider the (thin) SVD $\mathcal{F}\mathcal{L} = \boldsymbol{W}\boldsymbol{D}\boldsymbol{V}^T$ (as described in section 1) to deduce some useful facts about the vector $\mathcal{F}\mathcal{L}\delta\boldsymbol{S}$. Firstly, note that the $N \times 1$ orthogonal column vectors $\boldsymbol{W}_i$ of $\boldsymbol{W}$ and the $s \times 1$ column vectors $\boldsymbol{V}_i$ of $\boldsymbol{V}$, satisfy $\mathcal{F}\mathcal{L}\boldsymbol{V}_i = \sigma_i \boldsymbol{W}_i$, where $\sigma_i$ the singular vectors of $\mathcal{F}\mathcal{L}$. If we define $\boldsymbol{U}_i = \mathcal{F}^{-1}\boldsymbol{W}_i$, $i = 1, 2, \ldots, s$ then

$$\mathcal{L}\boldsymbol{V}_i = \sigma_i \boldsymbol{U}_i \tag{10}$$

where the $N = (n + n^2)$-dimensional vectors $\boldsymbol{U}_i$ can be written as $\boldsymbol{U}_i = (\boldsymbol{U}_i^{\boldsymbol{\mu}}, \boldsymbol{U}_i^{\boldsymbol{\Sigma}})'$ to reflect the correspondence of the first $n$ entries to the mean vector $\boldsymbol{\mu}$ and the last $n^2$ entries to the covariance matrix $\boldsymbol{\Sigma}$.

By (10), up to terms that are $\mathrm{O}(\|\delta\boldsymbol{S}\|^2)$,

$$\delta\Theta = \mathcal{L}\cdot\delta\boldsymbol{S} = \mathcal{L}\cdot\sum_i (V_i^T \cdot \delta\boldsymbol{S})V_i = \sum_i (\sigma_i \boldsymbol{V}_i^T \cdot \delta\boldsymbol{S})\boldsymbol{U}_i = \sum_i (\sum_j \underline{\mathbf{s}}_{ij}\delta\boldsymbol{S}_j)\boldsymbol{U}_i = \boldsymbol{U}\cdot\underline{\mathbf{s}}\cdot\delta\boldsymbol{S}$$

i.e.

$$\mathcal{L} = \boldsymbol{U}\cdot\underline{\mathbf{s}} \tag{11}$$

and therefore

$$\|\delta\Theta\|_{\boldsymbol{S}_0} = \|\underline{\mathbf{s}}\cdot\delta\boldsymbol{S}\| + \mathrm{O}(\|\delta\boldsymbol{S}\|^2)$$

since the $\boldsymbol{U}_i$ are orthonormal in the $\langle\cdot,\cdot\rangle_{\boldsymbol{S}_0}$ metric and the coefficient of $\boldsymbol{U}_i$ in $\boldsymbol{U}\cdot\underline{\mathbf{s}}\cdot\delta\boldsymbol{S}$ is the $i$th coordinate of $\underline{\mathbf{s}}\cdot\delta\boldsymbol{S}$.

These equations are the reason we call $\underline{\mathbf{s}} = D^{1/2}\boldsymbol{V}^T$ the *sensitivity matrix*. It describes how the distribution changes as the signal $S$ is changed.

Similarly, up to terms that are $\mathrm{O}(\|\delta\boldsymbol{S}\|^2)$,

$$\mathcal{F}(\delta\Theta) = \sum_{i=1}^{s}\sum_{j=1}^{s} \boldsymbol{W}_i \underline{\mathbf{s}}_{ij}\delta\boldsymbol{S}_j = \boldsymbol{W}\cdot\underline{\mathbf{s}}\cdot\delta\boldsymbol{S} \tag{12}$$

i.e. $\mathcal{F}\mathcal{L} = \boldsymbol{W}\cdot\underline{\mathbf{s}}$. We can now make a few useful observations:

1. Since the FIM is $\mathcal{I}(\boldsymbol{S}_0) = \underline{\mathbf{s}}^T\underline{\mathbf{s}}$, the eigenvalues of $\mathcal{I}(\boldsymbol{S}_0)$ are the squares of the singular values $\sigma_i$ of $\underline{\mathbf{s}}$ and the right singular vectors of $\underline{\mathbf{s}}$, $\boldsymbol{V}_i$, are the eigenvectors of $\mathcal{I}$. Further, since the column vectors $\boldsymbol{V}_i$ of $\boldsymbol{V}$ are unit vectors the squared norm of the $i$th row of $\underline{\mathbf{s}}$, $\sum_j \underline{\mathbf{s}}_{ij}^2$ is $\sigma_i^2$.

2. Equation (12) shows that the change in the probability distribution $MVN(\boldsymbol{\mu}(\boldsymbol{S}), \boldsymbol{\Sigma}(\boldsymbol{S}))$ produced by a change $\boldsymbol{S} \to \boldsymbol{S} + \delta\boldsymbol{S}$, according to the Fisher information metric, is a weighted sum of the vectors $\mathcal{F}(\boldsymbol{U}_j^{\boldsymbol{\mu}}, \boldsymbol{U}_j^{\boldsymbol{\Sigma}})^T$ with weights-coefficients $\sigma_j \boldsymbol{V}_j^T\delta\boldsymbol{S}$. The change in the probability distribution is reflected to the mean through the $\boldsymbol{U}_j^{\boldsymbol{\mu}}$ directions and the covariance matrix through the $\boldsymbol{U}_j^{\boldsymbol{\Sigma}}$ directions.

3. The coefficients $\sigma_j \boldsymbol{V}_j^T\delta\boldsymbol{S}$ are proportional to the singular values $\sigma_j$ and the inner products $\boldsymbol{V}_j^T\delta\boldsymbol{S}$. The latter are the coordinates of $\delta\boldsymbol{S}$ in the orthonormal basis of $\mathcal{S}$ defined by the columns of the matrix $\boldsymbol{V}$. Therefore, if the singular values are chosen in non-increasing order, i.e. $\sigma_1 \geq \cdots \geq \sigma_s$, the largest change in the probability distribution, subject to fixed $\|\delta\boldsymbol{S}\|$, occurs when the change $\delta\boldsymbol{S}$ is parallel to $\boldsymbol{V}_1$ that corresponds to the largest singular value $\sigma_1$. If the singular values decay fast, there are only a few directions of the signal space that can produce a relatively large change in the $\mathrm{MVN}(\boldsymbol{\mu}(S), \boldsymbol{\Sigma}(S))$ distribution (subject to fixed $\|\delta\boldsymbol{S}\|$).

4. Furthermore, the overall contribution of each coordinate $\delta \boldsymbol{S}_i$ of $\delta \boldsymbol{S}$ in the change of the probability distribution is measured, according to the Fisher information metric, by $\sum_{j=1}^{s} \boldsymbol{W}_j \underline{\mathbf{s}}_{ij} = \boldsymbol{W} \underline{\mathbf{s}}_i$. That is, if $\delta \boldsymbol{S} = \epsilon \boldsymbol{e}_i$ with $\boldsymbol{e}_i \in \mathbb{R}^s$ the usual unit vector with only non-zero entry $e_{ii} = 1$, then the corresponding change in the probability distribution $\mathrm{MVN}(\boldsymbol{\mu}(\boldsymbol{S}), \boldsymbol{\Sigma}(\boldsymbol{S}))$ according to the Fisher information metric is

$$\epsilon \mathcal{F}\mathcal{L} \boldsymbol{e}_i = \epsilon \boldsymbol{W} \underline{\mathbf{s}}_i. \tag{13}$$

##### 2.5.2 Optimality of $\underline{\mathbf{s}}$

The above sensitivity matrix is optimal for capturing as much sensitivity as possible in the low order principal components of $\delta\Theta = (\delta\boldsymbol{\mu}, \delta \mathrm{vec}(\boldsymbol{\Sigma}))^T$. That is, for any (sensitivity) matrix $\underline{\mathbf{s}}'$ which for some orthogonal matrix $\boldsymbol{U}'$ satisfies

$$\delta\Theta = \boldsymbol{U}'(\underline{\mathbf{s}}')^T \delta \boldsymbol{S} + \mathrm{O}(\|\delta \boldsymbol{S}\|^2),$$

the sensitivity matrix $\underline{\mathbf{s}}$ satisfies the following inequalities for all $\ell < k$,

$$\sum_{i \le \ell} \sum_j \underline{\mathbf{s}}_{ij}^2 \ge \sum_{i \le \ell} \sum_j \underline{\mathbf{s}}_{ij}'^2$$

and

$$\sum_{i \ge \ell} \sum_j \underline{\mathbf{s}}_{ij}^2 \le \sum_{i \ge \ell} \sum_j \underline{\mathbf{s}}_{ij}'^2 \tag{14}$$

i.e. among all such sensitivity matrices $\underline{\mathbf{s}}$ squeezes as much of the sensitivity effect as possible into the lower $i$ components.

For the above reasons, we call $\underline{\mathbf{s}}_{ij}$, for $j = 1, \dots, k$, the principal coefficients of sensitivity of $\mathrm{MVN}(\boldsymbol{\mu}(\boldsymbol{S}), \boldsymbol{\Sigma}(\boldsymbol{S}))$ to changes in the $i$th component of the signal $S_i$, $i = 1, \dots, k$.

##### 2.5.3 The multiplexing capacities of a model

So far as modelling is concerned the typical situation is where the signals $S_i$ change some of the parameters $\theta_j$, $j = 1, \dots, m$ of the model so that $\boldsymbol{\theta} = \boldsymbol{\theta}(\boldsymbol{S})$. Assume that the parameters that are changed by the signals have been reordered so that $\nu_j^2 = 2 \cdot \mathrm{MX}(\theta_1, \dots, \theta_j)$ satisfy $\nu_1 \ge \nu_2 \ge \dots \ge \nu_s$ as can be done using the results of Sect. 2.4.1. Assume also that the signals have also been reordered so that $\nu_j'$, where $\nu_j'^2 = 2 \cdot \mathrm{MX}(S_1, \dots, S_j)$, similarly decrease. We show that then

$$\prod_{i=1}^{s} \nu_j' = (\det d\boldsymbol{\theta})^2 \prod_{i=1}^{s} \nu_j. \tag{15}$$

where $d\boldsymbol{\theta} = d\boldsymbol{\theta}(\boldsymbol{S})$ is the derivative of $\boldsymbol{\theta}$ with respect to $\boldsymbol{S}$ at $\boldsymbol{S} \in \mathcal{S}$ as in Sect. 2.4.1. It follows from (15) that if these parameters have a large multiplexing capacity then signals for which $d\boldsymbol{\theta}$ is well-conditioned will also multiplex well.

To show eq. (15) we note that in this case the FIM of the parameters $\mathcal{I}_\theta$ and signals $\mathcal{I}(\boldsymbol{S})$ are related by equation (1). Consider the corresponding sensitivity matrices $\underline{\mathbf{s}}_\theta$ and $\underline{\mathbf{s}}(\boldsymbol{S})$, respectively. By equation (1)

$$(\underline{\mathbf{s}}(\boldsymbol{S}))^T \underline{\mathbf{s}}(\boldsymbol{S}) = d\boldsymbol{\theta}^T \mathcal{I}_\theta d\boldsymbol{\theta} = (\underline{\mathbf{s}}_{\boldsymbol{\theta}} d\boldsymbol{\theta})^T (\underline{\mathbf{s}}_{\boldsymbol{\theta}} d\boldsymbol{\theta}).$$

Therefore, the matrices $\underline{\mathbf{s}}(\boldsymbol{S})$ and $\underline{\mathbf{s}}_{\boldsymbol{\theta}}$ have the same singular values and the same determinant.

Let $\boldsymbol{Q}\boldsymbol{R} = \boldsymbol{P}\underline{\mathbf{s}}_\theta$ and $\boldsymbol{Q}'\boldsymbol{R}' = \boldsymbol{P}'\underline{\mathbf{s}}(\boldsymbol{S})$ be the $QR$ decompositions from Sect. 2.4.1. Then $\nu_k = \mathrm{MX}(\theta_1, \dots, \theta_k) = R_{kk}^2/2, \quad k = 1, \dots, s.$ and $\nu_k' = \mathrm{MX}(S_1, \dots, S_k) = R_{kk}'^2/2, \quad k = 1, \dots, s.$ Thus,

$$\prod_{i=1}^{s} \nu_j' / \prod_{i=1}^{s} \nu_j = \prod_{i=1}^{s} R_{jj}'^2 / \prod_{i=1}^{s} R_{jj}^2 = \det \underline{\mathbf{s}}(\boldsymbol{S})^2 / \det \underline{\mathbf{s}}_\theta^2 = (\det d\boldsymbol{\theta})^2$$

#### 3 Reaction Networks & Stochastic NF-$\kappa$B model

Stochastic models of cellular processes in signaling and regulatory systems are usually described in terms of reaction networks. A system of multiple different molecular subpopulations has *state vector*, $\boldsymbol{Y}(t) = (Y_1(t), \ldots, Y_n(t))^T$ where $Y_i(t)$, $i = 1, \ldots, n$, denotes the number of molecules of each species at time $t$. These molecules undergo a number of possible reactions (e.g. transcription, translation, degradation) where the reaction of index $j$ changes $\boldsymbol{Y}(t)$ to $\boldsymbol{Y}(t) + \boldsymbol{\nu}_j$, $\boldsymbol{\nu}_j \in \mathbb{R}^n$. The vectors $\boldsymbol{\nu}_j$ are called *stoichiometric* vectors. Each reaction occurs randomly at a rate $w_j(\boldsymbol{Y}(t))$ (often called the intensity of the reaction), which is a function of $\boldsymbol{Y}(t)$ but may also depend on $t$.

Therefore the state of the system at some time $t$ is

$$\boldsymbol{Y}(t) = \boldsymbol{Y}(0) + \sum_j \boldsymbol{\nu}_j Z_j \left( \int_0^t w_j(\boldsymbol{Y}(s))ds \right), \tag{16}$$

with $Z_j$ independent unit Poisson processes corresponding to the $j$-th reaction channel. The Stochastic Simulation algorithm (SSA) [4] use the properties of the above stochastic process to generate stochastic trajectories. These have exactly the same distribution with $\boldsymbol{Y}(t)$. More specifically, the SSA, often referred as Gillespie algorithm, generates the time to the first next reaction, $\Delta t$, at a given time $t$ that has an exponential distribution with rate $w_0(\boldsymbol{Y}(t)) = \sum_j w_j(\boldsymbol{Y}(t))$ and the type of the next reaction that is of type $j'$ with probability $w_{j'}(\boldsymbol{Y}(t))/w_0(\boldsymbol{Y}(t))$. Then the time $t$ is updated to $t + \Delta t$ and the state is updated to become $\boldsymbol{Y}(t) + \boldsymbol{\nu}_{j'}$. The same steps are repeated until $t > T$, for some time $T > 0$.

##### 3.1 System size parameter

It is common in studying stochastic systems to introduce a system size $\Omega$ which is a parameter that occurs in the intensities of the reactions $w_j(\boldsymbol{Y}(t))$. The precise description of this parameter depends on the system. In population models it might be considered to be of the same order of magnitude as the total population size while in chemical systems and cellular biology a natural choice is to use molar concentrations and therefore regard $\Omega$ as Avogadro's number in the appropriate molar units (e.g. $\mathrm{nM}^{-1}$) multiplied by the volume of the reacting solution (e.g. the cell) in appropriate units (e.g. in litres (L)). In the NF-$\kappa$B system, we consider it has units L/$\mu$M.

The system size governs the size of the state fluctuations and therefore the size of the jumps. Larger system sizes generally imply less fluctuations and vice versa. In a certain sense the system size parameter is just a mathematical convenience to enable the study of the dependence of stochastic fluctuations upon system size. Indeed, the methods we develop in this paper can and should be applied to systems that do not involve a system size parameter but then it will be necessary to ensure that the population sizes achieved in the given system are large enough.

While having a system size parameter is not necessary to apply our methods, it allows one to study the dependence of stochastic fluctuations upon system size and to calculate the deterministic equations that describe the evolution of the concentration vector $\boldsymbol{X}(t) = \boldsymbol{Y}(t)/\Omega$ in the limit of $\Omega \to \infty$ (see next section). A sufficient condition to derive this limit is that the rates $w_j(\boldsymbol{Y}(t))$ depend upon $\Omega$ (cf. [8, 9, 10]) as

$$w_j(\boldsymbol{Y}) = \Omega u_j(\boldsymbol{Y}/\Omega). \tag{17}$$

where $u_j(\boldsymbol{x})$ the macroscopic ($\Omega \to \infty$) rates that generally depend on the concentration vector $\boldsymbol{x} = \boldsymbol{x}(t)$. For example, linear degradation rates have the form $w_{\text{deg}}(N) = cN$, where $N$ stands for the relevant population number. The rate $w_{\text{deg}}(N)$ is equal to $\Omega u_{deg}(n)$, where $u_{deg}(n) = cn$ the macroscopic rate and $n = N/\Omega$ the corresponding concentration level. Similarly binding reaction rates have the form $w_{\text{bind}}(N_1, N_2) = cN_1N_2 = \Omega u_{\text{bind}}(n_1, n_2) = \tilde{c}n_1n_2$ where $\tilde{c} = c\Omega$, and $n_1 = N_1/\Omega$, $n_2 = N_2/\Omega$.

#### 3.2 The classical deterministic approximation

Assuming condition (17), the equation (16) describing the time evolution of $\boldsymbol{Y}(t)$ can be written in terms of $\boldsymbol{X} = \boldsymbol{Y}/\Omega$ as

$$\boldsymbol{X}(t) = \boldsymbol{X}(0) + \sum_j \boldsymbol{\nu}_j \Omega^{-1} Z_j \left( \int_0^t \Omega u_j(\boldsymbol{X}(s)) ds \right). \tag{18}$$

Using the law of large numbers, as $\Omega \to \infty$, we have that

$$\boldsymbol{x}(t) = \boldsymbol{x}(0) + \sum_j \boldsymbol{\nu}_j \int_0^t u_j(\boldsymbol{x}(s)) ds,$$

where here $\boldsymbol{x}(t)$ is the limit of convergence in probability for $\boldsymbol{X}(t)$ as $\Omega \to \infty$. Consequently, $\boldsymbol{x}(t)$ satisfies the ordinary differential equation (ODE)

$$\dot{\boldsymbol{x}} = \boldsymbol{F}(\boldsymbol{x}(t)), \quad \boldsymbol{F}(\boldsymbol{x}) = \sum_j \boldsymbol{\nu}_j u_j(\boldsymbol{x}), \tag{19}$$

which is the classical macroscopic-deterministic approximation of the time-evolution of the concentration levels of the system. In matrix form, $\boldsymbol{F}(\boldsymbol{x}) = \boldsymbol{M}\boldsymbol{u}(\boldsymbol{x})$ where $\boldsymbol{M} = [\boldsymbol{\nu}_1 \, \boldsymbol{\nu}_2 \, \ldots \, \boldsymbol{\nu}_r]$ the $n \times r$ stoichiometry matrix and $\boldsymbol{u}(\boldsymbol{x}(t)) = (\boldsymbol{u}_1(\boldsymbol{x}(t)), \boldsymbol{u}_2(\boldsymbol{x}(t)), \ldots, \boldsymbol{u}_r(\boldsymbol{x}(t)))^T$ the $r \times 1$ macroscopic rates vector. Herein we use $\dot{\boldsymbol{x}}$ to denote the time-derivative $d\boldsymbol{x}/dt$.

Throughout we consider autonomous systems (i.e. $\boldsymbol{F}(t) = \boldsymbol{F}(\boldsymbol{x}(t))$, but the method applies to non-autonomous systems with appropriate simple re-parameterisation (see for example [6]).

#### 3.3 Stochastic NF-$\kappa$B model

The model of NF-$\kappa$B system used in this SI describes the oscillatory response of the system following stimulation by tumor necrosis factor alpha (TNF$\alpha$). The system is initiated at its stable equilibrium state reached under the parameter settings of no stimulation (see Table S2). This is followed by a period of TNF$\alpha$ stimulation either continuous or in the form of repeated pulses (e.g. 5 min stimulation every 100 minutes). The stimulation cause a transient oscillation, which in the case of pulses is repeated after every pulse, and in the case of continuous stimulation relaxes to a stable limit cycle.

The model of NF-$\kappa$B considered here is a slight modification of that in [2]. In particular, the redundant (no feedback) variable of the phosphorylated I$\kappa$B$\alpha$ is removed as well as the phosphorylated complex of NF-$\kappa$B and I$\kappa$B$\alpha$ and the inactive IKK that are also redundant because the sum of IKK and NF-$\kappa$B are fixed in the model in [2]. These changes ensure that there are no rank-deficiencies in the covariance matrices of the LNA of the system.

We also write the system in a form where concentrations are in terms of the same volume. The original system in [2] is written in cytoplasmic concentrations for all species except the nuclear NF-$\kappa$B, I$\kappa$B$\alpha$ and their nuclear complex that are written in nuclear concentrations and are set to be 3.3 larger than cytoplasmic concentrations. Here, the necessary adjustments to the system equations are made so that all concentrations are of the same scale and no further conversions are necessary.

The ODE model for the NF-$\kappa$B system is therefore given in Table S3. The solution of the ODE system is provided in Fig. S1. The reactions and their rates used for the SSA are provided in Table **??**. The parameter values used in our simulations are provided in Table S4.

| | name | description | initial value |
|---|---|---|---|
| 1 | $N_c$ | Free Cytoplasmic NF-$\kappa$B | 0.0036 |
| 2 | $I_c$ | Free Cytoplasmic I$\kappa$B$\alpha$ | 0.016 |
| 3 | $(N\text{-}I)_c$ | Cytoplasmic complex of NF-$\kappa$B and I$\kappa$B$\alpha$ | 0.072 |
| 4 | $N_n$ | Free nuclear NF-$\kappa$B | 0.0042 |
| 5 | $I_n$ | Free nuclear I$\kappa$B$\alpha$ | 0.0013 |
| 6 | $(N\text{-}I)_n$ | Nuclear complex NF-$\kappa$B and I$\kappa$B$\alpha$ | 2.62e-4 |
| 7 | $I_m$ | I$\kappa$B$\alpha$ mRNA | 2.04e-05 |
| 8 | $K_n$ | IKK in neutral state | 0.08 |
| 9 | $K_a$ | IKK in active state | 2.22e-16 |
| 10 | $A_m$ | A20 mRNA | 1.27e-05 |
| 11 | $A$ | A20 | 0.0014 |

**Table S2:** *The variables of the base NF-$\kappa$B system and the initial conditions (in $\mu M$, concentration per cell) used to derive the ODE solution. The initial conditions are set to the equilibrium fixed point derived under no stimulation (i.e. the state of the system before activation).*

| | |
|---|---|
| 1 | $\dot{N}_c = k_{d1a}(N\text{-}I)_c - k_{a1a}N_cI_c - k_{i1}N_c + c_{5a}(N\text{-}I)_c + k_vk_{e1}N_n + k_{t2a}(\text{TNFKB} - N_c - (N\text{-}I)_c - N_n + (N\text{-}I)_n)$ |
| 2 | $\dot{I}_c = k_{d1a}(N\text{-}I)_c - k_{a1a}N_cI_c - k_{i3a}I_c + k_vk_{e3a}I_n - c_{4a}I_c + c_{2a}I_m - k_{c1a}K_aI_c$ |
| 3 | $(\dot{N\text{-}I})_c = k_{a1a}N_cI_c - k_{d1a}(N\text{-}I)_c + k_vk_{e2a}(N\text{-}I)_n - c_{5a}(N\text{-}I)_c - k_{c2a}K_a(N\text{-}I)_c$ |
| 4 | $\dot{N}_n = k_{d1a}(N\text{-}I)_n - k_vk_{a1a}N_nI_n + k_{i1}N_c - k_vk_{e1}N_n$ |
| 5 | $\dot{I}_n = k_{d1a}(N\text{-}I)_n - k_vk_{a1a}N_nI_n + k_{i3a}I_c - k_vk_{e3a}I_n - c_{4a}I_n$ |
| 6 | $(\dot{N\text{-}I})_n = k_vk_{a1a}(N\text{-}I)_n - k_{d1a}(N\text{-}I)_n - k_vk_{e2a}(N\text{-}I)_n$ |
| 7 | $\dot{I}_m = c_{1a}(N_n^h/(N_n^h + (k/k_v)^h)) - c_{3a}I_m$ |
| 8 | $\dot{K}_n = k_p(\text{TIKK} - K_n - K_a)(k_{bA20}/(k_{bA20} + A \cdot \text{TNF}\alpha)) - k_a\text{TNF}\alpha K_n$ |
| 9 | $\dot{K}_a = k_a\text{TNF}\alpha K_n - k_iK_a$ |
| 10 | $\dot{A}_m = c_1(N_n^h/(N_n^h + (k/k_v)^h)) - c_3A_m$ |
| 11 | $\dot{A} = c_2A_m - c_4A$ |

**Table S3:** *The ODE system of the base NF-$\kappa$B model used here which is the same as in [2] subject to some modifications explained in Section 3.3.*

| parameter | description | value | measurement unit |
|---|---|---|---|
| $k_v$ | C:N ratio | 3.300000 | – |
| $k_p$ | IKKn production | 0.0006 | $s^{-1}$ |
| $k_a$ | Activation caused by TNFa | 0.004 | $s^{-1}$ |
| $k_i$ | Spontaneous IKK activation | 0.003 | $s^{-1}$ |
| $k_{a1a}$ | NFkB-IkBa association | 0.5 | $\mu M^{-1}s^{-1}$ |
| $k_{d1a}$ | NFkB-IkBa dissociation | 0.0005 | $s^{-1}$ |
| $k_{c1a}$ | Catalysis of IKK-IkBa dimer | 0.074 | $s^{-1}$ |
| $k_{c2a}$ | Catalysis of IKK-IkBa-NFkB trimer | 0.37 | $s^{-1}$ |
| $k_{t2a}$ | degradation of IkBa (IKK dependent from trimer) | 0.1 | $s^{-1}$ |
| $c_{4a}$ | Free IkBa degradation | 0.0005 | $s^{-1}$ |
| $c_{5a}$ | NFkB complexed IkBa degradation | 0.000022 | $s^{-1}$ |
| $k_{i1}$ | NFkB nuclear import | 0.0026 | $s^{-1}$ |
| $k_{e1}$ | NFkB nuclear export | 0.000052 | $s^{-1}$ |
| $k_{e2a}$ | NFkB-IkBa nuclear export | 0.01 | $s^{-1}$ |
| $k_{i3a}$ | IkBa nuclear import | 0.00067 | $s^{-1}$ |
| $k_{e3a}$ | IkBa nuclear export | 0.000335 | $s^{-1}$ |
| $h$ | Order of hill function | 2 | – |
| $k$ | Hill constant | 0.0650 | $\mu$M/L |
| $c_{1a}$ | IkBa mRNA synthesis | 1.400e-07 | $\mu M^{-1}s^{-1}$ |
| $c_{2a}$ | IkBa translation rate | 0.5 | $s^{-1}$ |
| $c_{3a}$ | IkBa mRNA degradation | 0.0003 | $s^{-1}$ |
| $c_1$ | IkBa mRNA synthesis | 1.4e-07 | $\mu M^{-1}s^{-1}$ |
| $c_2$ | A20 mRNA translation | 0.5 | $s^{-1}$ |
| $c_3$ | A20 mRNA degradation | 0.00048 | $s^{-1}$ |
| $c_4$ | A20 degradation | 0.0045 | $s^{-1}$ |
| $k_{bA20}$ | Half-max A20 inhibition concentration | 0.0018 | $\mu$M/L |
| TNF$\alpha$ | Tumor necrosis factor alpha level | 10 | $ng/mL$ |
| TNF-$\kappa$B | total NF-$\kappa$B concentration | 0.08 | $\mu M$ |
| $TIKK$ | total IKK concentration | 0.08 | $\mu M$ |

**Table S4:** *The parameters of the base NF-$\kappa$B system and the values used to derive their ODE solution.*

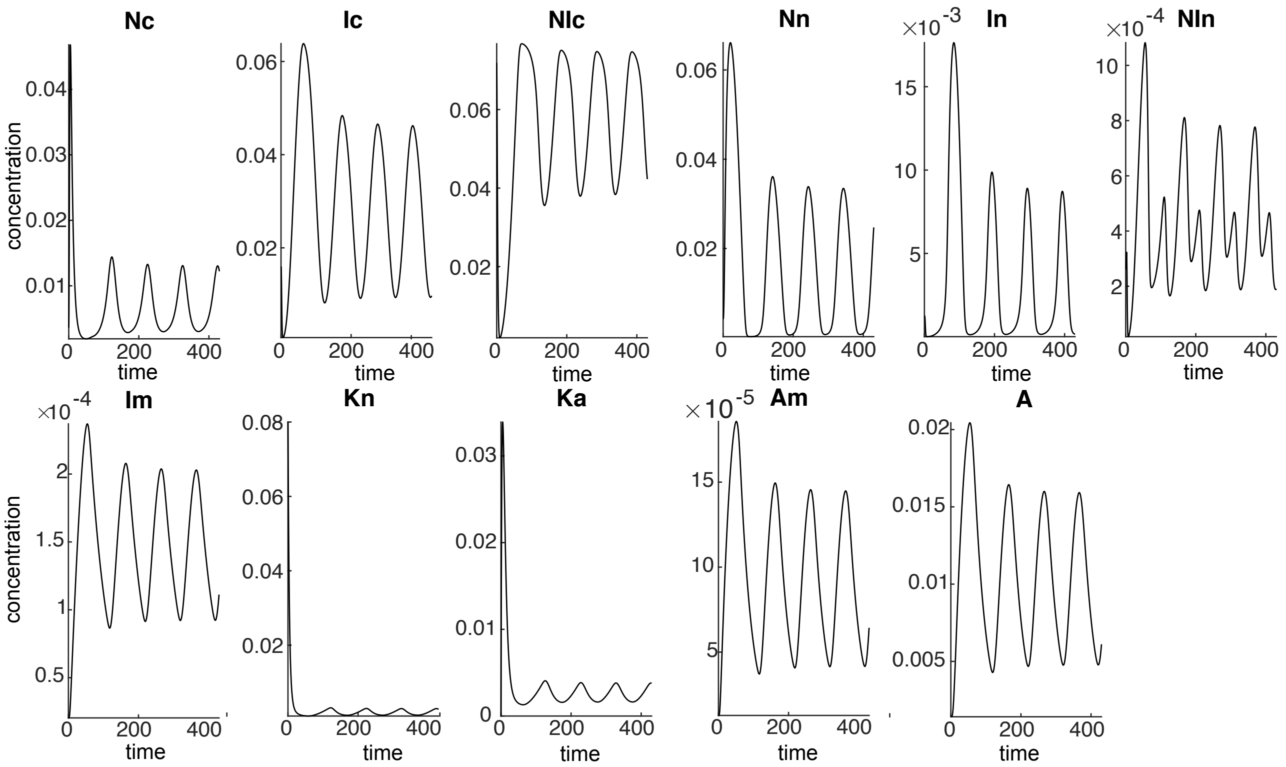

**Figure S1:** *The solution of the variables of the ODE system of the base NF-κB model.*

##### 3.4 The mNF-κB model

In addition to the variables and parameters in the base model, the mNF-κB model has the extra variables and parameters provided in tables S5 and S10, respectively. As you can see in table S6, which includes the ODE system of the mNF-κB model, the rest of the system is unchanged.

| | name | description | initial value |
|---|---|---|---|
| 1 | $N_c$ | Free Cytoplasmic NF-κB | 0.0036 |
| 2 | $I_c$ | Free Cytoplasmic IκBα | 0.016 |
| 3 | $(N\text{-}I)_c$ | Cytoplasmic complex of NF-κB and IκBα | 0.072 |
| 4 | $N_n$ | Free nuclear NF-κB | 0.0042 |
| 5 | $I_n$ | Free nuclear IκBα | 0.0013 |
| 6 | $(N\text{-}I)_n$ | Nuclear complex NF-κB and IκBα | 2.62e-4 |
| 7 | $I_m$ | IκBα mRNA | 2.04e-05 |
| 8 | $K_n$ | IKK in neutral state | 0.08 |
| 9 | $K_a$ | IKK in active state | 2.22e-16 |
| 10 | $A_m$ | A20 mRNA | 1.27e-05 |
| 11 | $A$ | A20 | 0.0014 |
| 12 | $N_m$ | Free Cytoplasmic NF-κB | 0.0018 |
| 13 | $N_m\text{-}I$ | Cytoplasmic complex of modified NF-κB and phosphorylated IκBα | 0.0360 |
| 14 | $N_m\text{-}I_p$ | Cytoplasmic complex of modified NF-κB and phosphorylated IκBα | 0 |
| 15 | $(N_m)_n$ | Free nuclear modified NF-κB | 0.0021 |
| 16 | $(N_m\text{-}I)_n$ | Free complex of modified NF-κB and phosphorylated IκBα | 0.0001 |

**Table S5:** *The variables of mNF-κB model and the initial conditions (in μM, concentration per cell) used to derive the ODE solution. The initial conditions are set to the equilibrium fixed point derived under no stimulation. The extra variables, compared to the base model, are marked in blue.*

| | |
|---|---|
| 1 | $\dot{N}_c = k_{d1a}(N\text{-}I)_c - k_{a1a}N_cI_c - k_{i1}N_c + c_{5a}(N\text{-}I)_c + k_vk_{e1}N_n$ $+k_{t2a}(\text{TNFKB} - N_c - (N\text{-}I)_c - N_n - (N\text{-}I)_n) - p_{a1}S_2\frac{N_c}{N_c+k_{pnf}} + p_{d1}\frac{N_m}{N_m+k_{mnf}}$ |
| 2 | $\dot{I}_c = k_{d1a}(N\text{-}I)_c - k_{a1a}N_cI_c - k_{i3a}I_c + k_vk_{e3a}I_n - c_{4a}I_c + c_{2a}I_m - k_{c1a}K_aI_c$ |
| 3 | $(\dot{N\text{-}I})_c = k_{a1a}N_cI_c - k_{d1a}(N\text{-}I)_c + k_vk_{e2a}(N\text{-}I)_n - c_{5a}(N\text{-}I)_c - k_{c2a}K_a(N\text{-}I)_c$ |
| 4 | $\dot{N}_n = k_{d1a}(N\text{-}I)_n - k_vk_{a1a}N_nI_n + k_{i1}N_c - k_vk_{e1}N_n$ |
| 5 | $\dot{I}_n = k_{d1a}(N\text{-}I)_n - k_vk_{a1a}N_nI_n + k_{i3a}I_c - k_vk_{e3a}I_n - c_{4a}I_n$ |
| 6 | $(\dot{N\text{-}I})_n = k_vk_{a1a}(N\text{-}I)_n - k_{d1a}(N\text{-}I)_n - k_vk_{e2a}(N\text{-}I)_n$ |
| 7 | $\dot{I}_m = c_{1a}\frac{(N_n+(N_m)_n)^h}{(N_n+(N_m)_n)^h+(k/k_v)^h} - c_{3a}I_m$ |
| 8 | $\dot{K}_n = k_p(\text{TIKK} - K_n - K_a)(k_{bA20}/(k_{bA20} + A \times \text{TNF}\alpha)) - k_a\text{TNF}\alpha K_n$ |
| 9 | $\dot{K}_a = k_a\text{TNF}\alpha K_n - k_iK_a$ |
| 10 | $\dot{A}_m = c_1\frac{(N_n+(N_m)_n)^h}{(N_n+(N_m)_n)^h+(k/k_v)^h} - c_3A_m$ |
| 11 | $\dot{A} = c_2A_m - c_4A$ |
| 12 | $\dot{N}_m = k_{d1a}(N_m\text{-}I) - k_{a1a}N_mI_c - k_{i1}N_m + k_vk_{e1}(N_m)_n + c_{5a}N_m\text{-}I + k_{t2a}(N_m\text{-}I_p)$ $+p_{a1}S_2\frac{N_c}{N_c+k_{pnf}} - p_{d1}\frac{N_m}{N_m+k_{mnf}}$ |
| 13 | $(\dot{N_m\text{-}I}) = k_{a1a}N_mI_c - k_{d1a}N_m\text{-}I + k_v * k_{e2a}N_m\text{-}I_n - c_{5a}N_m\text{-}I - k_{c2a}K_aN_m\text{-}I)$ |
| 14 | $(\dot{N_m\text{-}I_p}) = k_{c2a}K_aN_m\text{-}I - k_{t2a}N_m\text{-}I_p$ |
| 15 | $(\dot{N_m})_n = k_{d1a}N_m\text{-}I_n - k_vk_{a1a}(N_m)_nI_n + k_{i1}N_m - k_vk_{e1}(N_m)_n$ |
| 16 | $(\dot{N_m\text{-}I})_n = k_vk_{a1a}(N_m)_nI_n - k_{d1a}N_m\text{-}I_n - k_vk_{e2a}N_m\text{-}I_n$ |

**Table S6:** *The ODE system for the mNF-$\kappa$B model. This is an extension of the model provided in [2] to include modification of NF-$\kappa$B (see also Section 3.3). The extra equations, compared to the base model, are marked in blue.*

| parameter | description | value | measurement unit |
|---|---|---|---|
| $S_2$ | signal controlling NF-$\kappa$B modification | 1 | $s^{-1}$ |
| $p_{a1}$ | NF-$\kappa$B modification rate | 0.0175 | $s^{-1}$ |
| $p_{d1}$ | NF-$\kappa$B de-modification rate | 0.0175 | $s^{-1}$ |
| $k_{pnf}$ | half-max NF-$\kappa$B modification level | 0.01 | $\mu$M/L |
| $k_{mnf}$ | half-max NF-$\kappa$B un-modification level | 0.01 | $\mu$M/L |

**Table S7:** *The extra parameters of mNF-$\kappa$B system, compared to the base model, and the values used to derive their ODE solution.*

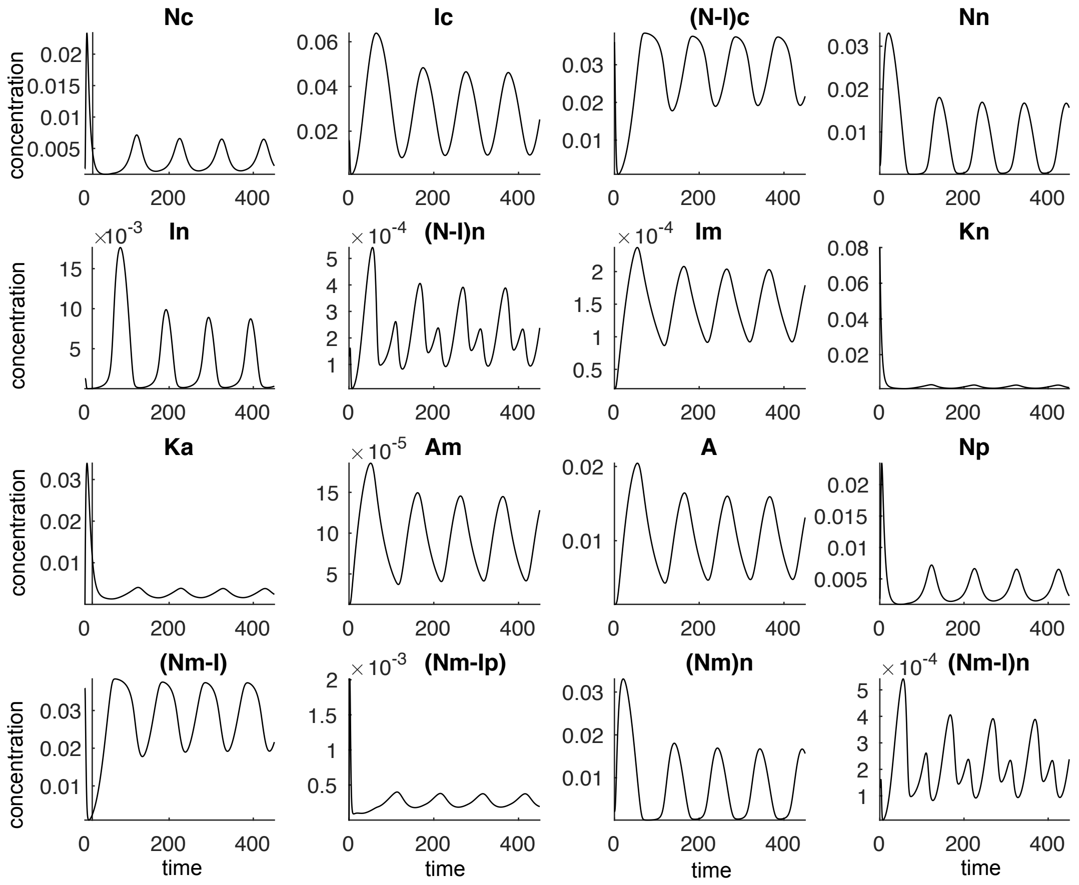

**Figure S2:** *The solution of the variables of the ODE system of the modified NF-$\kappa$B model.*

#### 3.5 Another model for NF-$\kappa$B modification

We now consider another model for NF-$\kappa$B modification where the NF-$\kappa$B molecules that are bound by I$\kappa$B$\alpha$ molecules, are transformed to another complex of modified NF-$\kappa$B molecules bounded by I$\kappa$B$\alpha$ molecules. For the modification to happen, a transition to an intermediate state needs to happen first. This transition is controlled by an independent signal $S_2$. A similar mechanism using an intermediate state is used for the reverse modification. The diagram of the model, which we will call amNF-$\kappa$B model, is depicted in Fig. S4(a), while the solutions of the ODE equations provided in table S9 are provided in Fig. S4. As we can see in Fig. S5(b), the singular values of the FIM of amNF-$\kappa$B are larger than the singular values of the base and mNF-$\kappa$B models. The multiplexing capacities are also increased with parameters related to this modification taking some of the higher values (see Fig. 3.5(c)). The sensitivity coefficients are generally increased with those parameters related to the modification having high sensntiviity (see Fig. 3.5(d)).

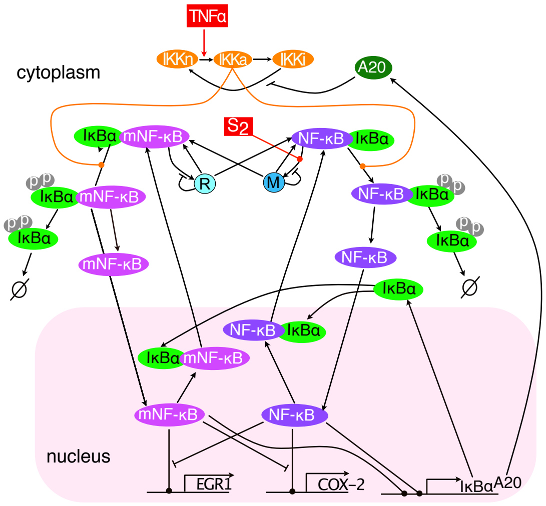

**Figure S3:** *Diagram of the main reactions of the amNF-κB model described in SI section 3.5;*

| | name | description | initial value |
|---|---|---|---|
| 1 | $N_c$ | Free Cytoplasmic NF-κB | 0.0018 |
| 2 | $I_c$ | Free Cytoplasmic IκBα | 0.015 |
| 3 | $(N\text{-}I)_c$ | Cytoplasmic complex of NF-κB and IκBα | 0.0348 |
| 4 | $N_n$ | Free nuclear NF-κB | 0.0029 |
| 5 | $I_n$ | Free nuclear IκBα | 0.0009 |
| 6 | $(N\text{-}I)_n$ | Nuclear complex NF-κB and IκBα | 0.0001 |
| 7 | $I_m$ | IκBα mRNA | $1.90 \times 10^{-5}$ |
| 8 | $K_n$ | IKK in neutral state | 0.08 |
| 9 | $K_a$ | IKK in active state | $2.22 \times 10^{-5}$ |
| 10 | $A_m$ | A20 mRNA | $1.19 \times 10^{-5}$ |
| 11 | $A$ | A20 | $1.19 \times 10^{-5}$ |
| 12 | $N_m$ | Free Cytoplasmic NF-κB | 0.0018 |
| 13 | $N_m\text{-}I$ | Cytoplasmic complex of modified NF-κB and phosphorylated IκBα | 0.0347 |
| 14 | $N_m\text{-}I_p$ | Cytoplasmic complex of modified NF-κB and phosphorylated IκBα | 0 |
| 15 | $(N_m)_n$ | Free nuclear modified NF-κB | 0.0028 |
| 16 | $(N_m\text{-}I)_n$ | Free complex of modified NF-κB and phosphorylated IκBα | 0.0001 |
| 17 | $M$ | Transient modification state | 0.0005 |
| 18 | $R$ | Transient de-modification state | 0.0004 |

**Table S8:** *The variables of the amNF-κB model described in section 3.5 and the initial conditions (in μM, concentration per cell) used to derive the ODE solution. The initial conditions are set to the equilibrium fixed point derived under no stimulation. The extra variables, compared to the base model, are marked in blue.*

| | |
|---|---|
| 1 | $\dot{N}_c = k_{d1a}(N\text{-}I)_c - k_{a1a}N_cI_c - k_{i1}N_c + c_{5a}(N\text{-}I)_c + k_vk_{e1}N_n$ <br> $+k_{t2a}(\text{TNFKB} - N_c - (N\text{-}I)_c - N_n - (N\text{-}I)_n)$ |
| 2 | $\dot{I}_c = k_{d1a}(N\text{-}I)_c - k_{a1a}N_cI_c - k_{i3a}I_c + k_vk_{e3a}I_n - c_{4a}I_c + c_{2a}I_m - k_{c1a}K_aI_c$ <br> $+k_{d1a}(N_m - I) - k_{a1a}N_mI_c$ |
| 3 | $(\dot{N\text{-}I})_c = k_{a1a}N_cI_c - k_{d1a}(N\text{-}I)_c + k_vk_{e2a}(N\text{-}I)_n - c_{5a}(N\text{-}I)_c - k_{c2a}K_a(N\text{-}I)_c$ <br> $+p_{k2}R + p_{d1}M - p_{a1}(N\text{-}I)_c(\text{TM}\times S2 - M)$ |
| 4 | $\dot{N}_n = k_{d1a}(N\text{-}I)_n - k_vk_{a1a}N_nI_n + k_{i1}N_c - k_vk_{e1}N_n$ |
| 5 | $\dot{I}_n = k_{d1a}(N\text{-}I)_n - k_vk_{a1a}N_nI_n + k_{i3a}I_c - k_vk_{e3a}I_n - c_{4a}I_n$ <br> $+k_{d1a}(N_m - I)_n - k_vk_{a1a}(N_m)_n \times I_n$ |
| 6 | $(\dot{N\text{-}I})_n = k_vk_{a1a}(N\text{-}I)_n - k_{d1a}(N\text{-}I)_n - k_vk_{e2a}(N\text{-}I)_n$ |
| 7 | $\dot{I}_m = c_{1a}\left(\frac{N_n^h}{N_n^h + (k/k_v)^h} + \frac{(N_m)_n^h}{(N_m)_n^h + (k/k_v)^h}\right) - c_{3a}I_m$ |
| 8 | $\dot{K}_n = k_p(\text{TIKK} - K_n - K_a)(k_{bA20}/(k_{bA20} + A \times \text{TNF}\alpha)) - k_a\text{TNF}\alpha K_n$ |
| 9 | $\dot{K}_a = k_a\text{TNF}\alpha K_n - k_iK_a$ |
| 10 | $\dot{A}_m = c_1\left(\frac{N_n^h}{N_n^h + (k/k_v)^h} + \frac{(N_m)_n^h}{(N_m)_n^h + (k/k_v)^h}\right) - c_3A_m$ |
| 11 | $\dot{A} = c_2A_m - c_4A$ |
| 12 | $\dot{N}_m = k_{d1a}(N_m\text{-}I) - k_{a1a}N_mI_c - k_{i1}N_m + k_vk_{e1}(N_m)_n + c_{5a}(N_m\text{-}I)$ <br> $+k_{t2a}(N_m\text{-}I_p)$ |
| 13 | $(\dot{N_m\text{-}I}) = k_{a1a}N_mI_c - k_{d1a}(N_m\text{-}I) + k_vk_{e2a}(N_m\text{-}I)_n - c_{5a}(N_m\text{-}I)$ <br> $-k_{c2a}K_a(N_m\text{-}I) + p_{k1}M + p_{d2}R - p_{a2}(N_m\text{-}I_p)(\text{TR} - \text{R})$ |
| 14 | $(\dot{N_m\text{-}I_p}) = k_{c2a}K_a(N_m\text{-}I) - k_{t2a}(N_m\text{-}I_p)$ |
| 15 | $(\dot{N_m})_n = k_{d1a}(N_m\text{-}I)_n - k_vk_{a1a}(N_m)_nI_n + k_{i1}N_m - k_vk_{e1}(N_m)_n$ |
| 16 | $(\dot{N_m\text{-}I})_n = k_vk_{a1a}(N_m)_nI_n - k_{d1a}N_m\text{-}I_n - k_vk_{e2a}(N_m\text{-}I)_n$ |
| 17 | $\dot{M} = p_{a1}(N\text{-}I)_c(\text{TM}\times S2 - M) - p_{d1}M - p_{k1}M$ |
| 18 | $\dot{R} = p_{a2}(N_m\text{-}I_p)(\text{TR-R}) - p_{k2}R - p_{d2}R$ |

**Table S9:** *The ODE system for the amNF-$\kappa$B model described in section 3.5. The extra equations, compared to the base model, are marked in blue.*

| parameter | value | measurement unit |
|---|---|---|
| $TM$ | 0.0008 | $\mu$M/L |
| $TR$ | 0.0004 | $\mu$M/L |
| $p_{a1}$ | 6 | $s^{-1}$ |
| $p_{d1}$ | 0.0035 | $s^{-1}$ |
| $p_{k1}$ | 0.0035 | $s^{-1}$ |
| $p_{a2}$ | 6 | $s^{-1}$ |
| $p_{d2}$ | 0.0035 | $s^{-1}$ |
| $p_{k2}$ | 0.0035 | $s^{-1}$ |
| $S2$ | 0.62 | |

**Table S10:** *The extra parameters of amNF-κB model described in section 3.5, compared to the base model, and the values used to derive their ODE solution.*

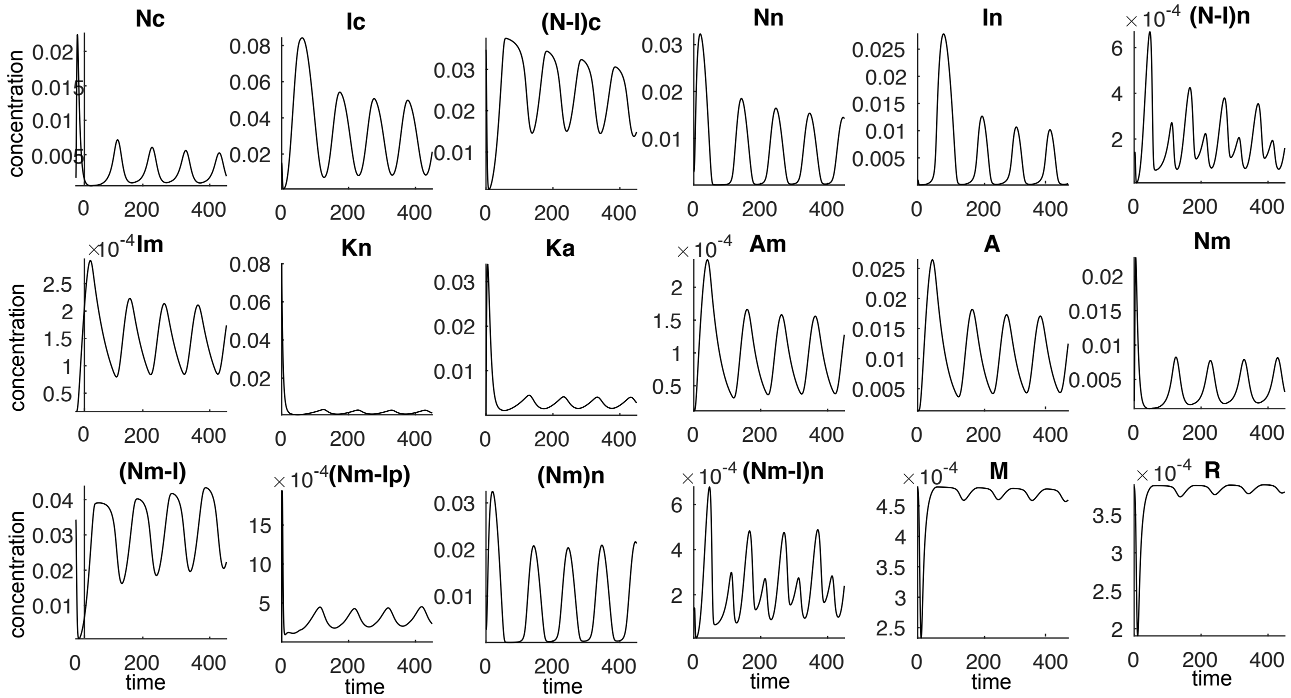

**Figure S4:** *The solution of the variables of the ODE system of the amNF-κB model described in section 3.5.*

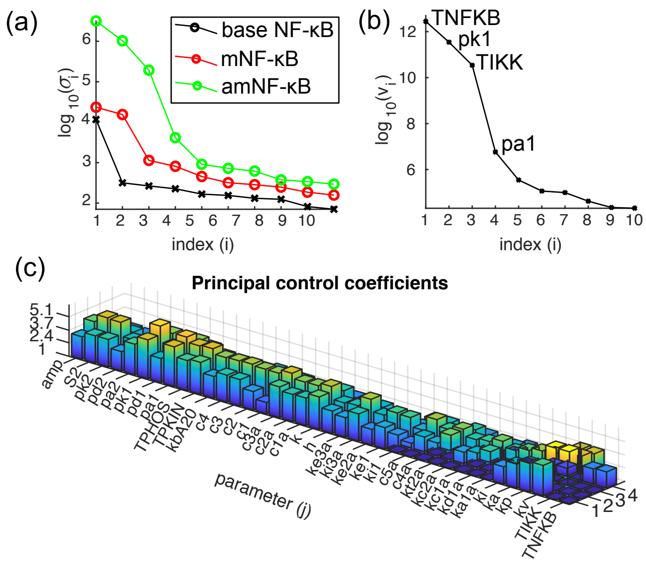

**Figure S5:** *(a) The singular values $\sigma_i$ of the amNF-κB , mNF-κB and base model; (b) The multiplexing capacities $v_j$ of the amNF-κB model. (c) The principal sensitivity coefficients of the amNF-κB model. Larger values indicate higher sensitivity of the amNF-κB model to changes in the value of the corresponding parameter.*

#### 4 LNA, pcLNA and calculating the FIM

We next describe the LNA [1] and pcLNA [13] as well as how to derive the FIM using pcLNA.

##### 4.1 The linear noise approximation (LNA)

The LNA ansantz describes the state $\boldsymbol{X}(t)$ at some time $t$ as the sum of the deterministic solution $\boldsymbol{x}(t)$ of (19) and the noise, $\boldsymbol{\xi}(t)$, scaled by $\Omega^{-1/2}$, i.e.

$$\boldsymbol{X}(t) = \boldsymbol{x}(t) + \Omega^{-1/2}\boldsymbol{\xi}(t). \tag{20}$$

The noise satisfies the linear stochastic differential equation (sde)

$$d\boldsymbol{\xi}(t) = \boldsymbol{J}(\boldsymbol{x}(t))\boldsymbol{\xi}(t)dt + \sqrt{\boldsymbol{M}\boldsymbol{U}(\boldsymbol{x}(t))\boldsymbol{M}^T}d\boldsymbol{W}_t \tag{21}$$

and $\boldsymbol{W}_t$ an $n \times 1$ Wiener process (i.e. $\boldsymbol{W}_t$ is a random vector of independent one-dimensional Wiener processes). Here $\boldsymbol{J}(\boldsymbol{x}(t))$ is the Jacobian matrix of (19), i.e.

$$\boldsymbol{J}(\boldsymbol{x}) = \left(\frac{d\boldsymbol{F}_i}{d\boldsymbol{x}_j}\right)_{ij}, \tag{22}$$

$\boldsymbol{M}$ the stoichiometry matrix, and $\boldsymbol{U}(\boldsymbol{x})$ the diagonal matrix with main diagonal the rates vector $\boldsymbol{u}(\boldsymbol{x})$. The noise sde can be solved and the solution can be written as

$$\boldsymbol{\xi}(t) = \boldsymbol{C}(t_0,t)\boldsymbol{\xi}(t_0) + \boldsymbol{\eta}(t_0,t), \quad \boldsymbol{\eta}(t_0,t) \sim \mathrm{MVN}(0, \boldsymbol{V}(t_0,t)), \tag{23}$$

where $\boldsymbol{C}(t_0,t)$ is the fundamental matrix of (19), i.e. $\boldsymbol{C}(t_0,t)$ is the solution of the initial value problem

$$\dot{\boldsymbol{C}} = \boldsymbol{J}\boldsymbol{C}, \quad \boldsymbol{C}(t_0,t_0) = \boldsymbol{I}, \tag{24}$$

and the symmetric positive-definite matrix $\boldsymbol{V}(t_0,t)$ is the solution of the initial value problem

$$\dot{\boldsymbol{V}}(t_0,t) = \boldsymbol{J}\boldsymbol{V} + \boldsymbol{V}\boldsymbol{J}^T + \boldsymbol{M}\boldsymbol{U}\boldsymbol{M}^T, \quad \boldsymbol{V}(t_0,t_0) = \boldsymbol{0}. \tag{25}$$

Equation (23) implies that if $\boldsymbol{\xi}(t_0) \sim MVN(\boldsymbol{m}_0, \boldsymbol{V}_0)$, then $P_{\text{LNA}}(\boldsymbol{\xi}(t)|\boldsymbol{\xi}(t_0) \sim MVN(\boldsymbol{m}_0, \boldsymbol{V}_0))$ is $MVN(\boldsymbol{m}_t, \boldsymbol{V}_t)$, with

$$\boldsymbol{m}_t = \boldsymbol{C}(t_0,t)\boldsymbol{m}_0, \qquad \boldsymbol{V}_t = \boldsymbol{C}(t_0,t)\boldsymbol{V}_0\boldsymbol{C}(t_0,t)^T + \boldsymbol{V}(t_0,t) \tag{26}$$

and the LNA anstantz in (20) that $P_{\text{LNA}}(\boldsymbol{X}(t)|\boldsymbol{\xi}(t_0) \sim MVN(\boldsymbol{m}_0, \boldsymbol{V}_0))$ is $MVN(\boldsymbol{\mu}_t, \boldsymbol{\Sigma}_t)$, where

$$\boldsymbol{\mu}_t = \boldsymbol{x}(t) + \Omega^{-1/2}\boldsymbol{m}_t, \qquad \boldsymbol{\Sigma}_t = \Omega^{-1}\boldsymbol{V}_t. \tag{27}$$

It is worth noting that the LNA can be derived by applying the Central Limit Theorem, as $\Omega \to \infty$, to the Poisson random variables in (18) (see [1],[8],[9]). It was also derived by van Kampen (see [10]) as an expansion, in powers of $\Omega$, of Kolmogorov's forward equations (also called Master equation) of the Poisson process $\boldsymbol{Y}(t)$ in (16) ignoring terms of order $O(\Omega^{-1/2})$.

#### 4.2 Phase corrected LNA (pcLNA)

The pcLNA method described below is very similar to this in [13]. However, for clarity, we provide a description more appropriate to the current context. Unlike [13], we consider solutions $\gamma$ given by $x = \boldsymbol{g}(t)$, $t \geq 0$, of (19) without assuming that $\gamma$ is a limit cycle. However, especially for low $\Omega$, that is where the fluctuations are large, the method provides better approximation when the underlying deterministic solutions are such that transversal sections are well defined (e.g. oscillatory systems).

We first provide several definitions relevant to the theory developed herein and in **I**.

**Transversal sections** Consider the solution $\gamma$ of the ode in (19) given by $\boldsymbol{x} = \boldsymbol{g}(t)$, $t > 0$. By a transversal section through $\gamma$ we mean an $(n-1)$-dimensional linear hyperplane $\mathcal{S}_x$ containing $\boldsymbol{x}$ and transversal to the tangent vector, $\boldsymbol{F}(\boldsymbol{x})$, to $\gamma$ at $\boldsymbol{x}$. A particular example is the hyperplane normal to $\gamma$ at $\boldsymbol{x}$. A transversal system is a family $\mathcal{S}_{\boldsymbol{g}(t)}$ of transversal sections that vary smoothly with $t$ in the sense that the unit normal vector to $\mathcal{S}_{\boldsymbol{g}(t)}$ varies smoothly with $t$.

A transversal system defines a mapping $\boldsymbol{G}$ of a neighbourhood of $\gamma$ onto $\gamma$ where if $\boldsymbol{X} \in \mathcal{S}_{\boldsymbol{x}}$ then $\boldsymbol{G}(\boldsymbol{X}) = \boldsymbol{x} \in \gamma$. In cases where $\boldsymbol{X}{=}\boldsymbol{X}(t)$ lies in more than one transversal sections, then $\boldsymbol{G}(\boldsymbol{X}(t)) = \boldsymbol{x}(s)$ with $s = \arg\min_{\{s' \colon \boldsymbol{G}(\boldsymbol{X}(t))=\boldsymbol{x}(s')\}} |s' - t|$ the closest time to $t$. We denote this mapping for the normal transversal system by $\boldsymbol{G}_N$.

**Computation of the normal mapping $\boldsymbol{G}_N(X(t))$** To compute the mapping $\boldsymbol{G}_N(\boldsymbol{X}) = \boldsymbol{x} \in \gamma$, of $\boldsymbol{X} = \boldsymbol{X}(t)$, for a given solution $\boldsymbol{x} = \boldsymbol{g}(t)$, $t > 0$, of the ode in (19), we wish to minimise the squared distance

$$\|\boldsymbol{X} - \boldsymbol{g}(s)\|^2$$

with respect to $s > 0$. Note that the minimum $s$ is achieved when

$$\boldsymbol{h}(s) = \boldsymbol{F}(\boldsymbol{g}(s))^T(\boldsymbol{X} - \boldsymbol{g}(s)) = 0,$$

i.e. when the tangent vector $\boldsymbol{F}(\boldsymbol{g}(s))$ is orthogonal to $\boldsymbol{X} - \boldsymbol{g}(s)$ and thus $\boldsymbol{X}$ lies on the normal transversal section $\mathcal{S}_{\boldsymbol{g}(s)}$. To find the appropriate $s$, we first use a Newton's optimisation to derive the approximate limit of the sequence

$$s_{n+1} = s_n - \boldsymbol{h}(s)/\dot{\boldsymbol{h}}(s), \qquad n = 0, 1, \ldots$$

for different $s_0$, stopping the search when the square of the step $\boldsymbol{h}(s)/\dot{\boldsymbol{h}}(s)$ is appropriately small. If different limits are achieved for different $s_0$, we choose the one that is closest to $t$. Note that other methods of optimisation could be used here.

**Adapted coordinate systems** An adapted coordinate system $\mathcal{C}_{\boldsymbol{g}(t)}$ at a point $\boldsymbol{g}(t)$ on $\gamma$ is one determined by a set of orthonormal basis vectors $\boldsymbol{e}_1(t), \dots, \boldsymbol{e}_n(t)$ with $\boldsymbol{e}_1(t)$ the unit normal vector to $\mathcal{S}_{\boldsymbol{g}(t)}$ and the vectors $\boldsymbol{e}_2(t), \dots, \boldsymbol{e}_n(t)$ forming an orthonormal basis of $\mathcal{S}_{\boldsymbol{g}(t)}$. If these are defined for $t$ in some interval in $\mathbb{R}^+$ then we always assume that the $\boldsymbol{e}_i(t)$ have smooth (i.e. $C^2$) dependence upon $t$. It is important that the coordinates are defined by an orthonormal basis in the original coordinates because this effectively preserves the covariance matrix in the sense that a covariance matrix $\boldsymbol{V}$ in the adapted coordinates is $\boldsymbol{R}\boldsymbol{V}\boldsymbol{R}^T$ in the original coordinates with $\boldsymbol{R}$ a real orthogonal matrix. In particular, the eigenvalues are preserved. The state $\boldsymbol{X}$ of the process (written in original coordinates corresponding to concentrations of each reacting molecule) relates to the adapted coordinates $(X^{(1)}, \boldsymbol{X}^{(2)})$ at $\boldsymbol{x} = \boldsymbol{g}(t)$ with $\boldsymbol{X}^{(2)} \in \mathcal{S}_{\boldsymbol{g}(t)}$ through the orthogonal matrix $\boldsymbol{R}_t = [\boldsymbol{R}_t^{(1)}\ \boldsymbol{R}_t^{(2)}]$ by

$$\left(X^{(1)}, \boldsymbol{X}^{(2)}\right)^T = \boldsymbol{R}_t(\boldsymbol{X} - \boldsymbol{x}(t)) \quad \text{and} \quad \boldsymbol{X} - \boldsymbol{x}(t) = \boldsymbol{R}_t^{(1)} X^{(1)} + \boldsymbol{R}_t^{(2)} \boldsymbol{X}^{(2)}. \tag{28}$$

**Transversal points** Consider a stochastic trajectory $\boldsymbol{X}(t_i)$, $i = 0, 1, 2, \dots$ and let $\boldsymbol{g}(t)$, $t \geq 0$, be a solution of (19) with $\boldsymbol{g}(t_0) = \boldsymbol{g}_0$. Suppose that the stochastic trajectory has initial condition $\boldsymbol{X}(0)$ in the neighbourhood of $\boldsymbol{g}_0$. For the SSA trajectories presented in **I**, this is satisfied by taking $\boldsymbol{X}_0 = \boldsymbol{g}_0$, but this is not necessary. Find the transversal section $\mathcal{S}_{\boldsymbol{g}(\boldsymbol{S}_0)}$ for which $\boldsymbol{G}_N(\boldsymbol{X}(t_0)) = \boldsymbol{g}(s_0)$ and $\boldsymbol{S}_0 = \arg\min_{\{s' : \boldsymbol{G}_N(\boldsymbol{X}(t_0)) = \boldsymbol{g}(s')\}} |s' - t_0|$. Then, for $i = 1, 2, \dots$, define the transversal sections $\mathcal{S}_{\boldsymbol{g}(s_i)}$, $i = 1, 2, \dots$, such that

$$\boldsymbol{G}_N(\boldsymbol{X}(t_i)) = \boldsymbol{g}(s_i) \quad \text{and} \quad s_i = \text{argmin}_{\{s' : \boldsymbol{G}_N(\boldsymbol{X}(t_i)) = \boldsymbol{g}(s')\}} |s' - s_{i-1} + (t_i - t_{i-1})|.$$

Suppose that we wish to find the point, $\boldsymbol{Q}_{\boldsymbol{x}}$, at which the stochastic trajectory intersects the transversal section $\mathcal{S}_{\boldsymbol{x} = \boldsymbol{g}(s)}$ where $s \in (s_i, s_{i+1})$ for some $i > 0$. Then some form of interpolation between $\boldsymbol{X}(t_i)$ and $\boldsymbol{X}(t_{i+1})$ needs to be used. For the SSA trajectories presented in **I**, we first use that there are no jumps between $t_i$ and $t_{i+1}$, then write $\boldsymbol{X}(t_i)$ in coordinates $(r_i, q_i)$ adapted at $\boldsymbol{g}(s)$, i.e. use the $\mathcal{C}_{g(s)}$ coordinates, and take the transversal point $Q_x = q_i$.

Note that backward moves and therefore multiple intersections to the same transversal section are possible. However, as we have shown in [13], the distribution of the number of intersections decreases exponentially and the distribution of the transversal points in different intersection is indistinguishable. Therefore, hereafter, we assume that in each trajectory there is only a single intersection to each transversal section, $\mathcal{S}_{\boldsymbol{x}}$, of $\boldsymbol{x} \in \gamma$, called transversal point $\boldsymbol{Q}_{\boldsymbol{x}}$.

##### 4.2.1 The pcLNA distribution

Consider a solution $\boldsymbol{g}(t)$, $t > 0$ of (19) that satisfies $\boldsymbol{g}_0 = \boldsymbol{g}(0)$ and a point $\boldsymbol{x} = \boldsymbol{g}(t)$ of this solution. Let $\mathcal{S}_{\boldsymbol{x} = \boldsymbol{g}(t)}$ be the corresponding transversal section at $\boldsymbol{x} = \boldsymbol{g}(t)$. We wish to approximate the distribution of the transversal point $\boldsymbol{Q}_{\boldsymbol{x} = \boldsymbol{g}(t)}$ as defined in section 4.2. We use the adapted coordinates $\mathcal{C}_x$ and write the state of the system at time $t$ as $\boldsymbol{X}(t) = (X^{(1)}(t), \boldsymbol{X}^{(2)}(t))^T$, with $\boldsymbol{X}^{(2)}(t) \in \mathcal{S}_{\boldsymbol{x}}$. Then the transversal distribution $P(\boldsymbol{Q}_{\boldsymbol{x}} | \boldsymbol{X}(0))$ is estimated in pcLNA by the distribution

$$P_{\text{LNA}}(\boldsymbol{X}^{(2)}(t) | \boldsymbol{X}(0)).$$

This is the LNA distribution of the state of the system on the transversal section $\mathcal{S}_{\boldsymbol{x} = \boldsymbol{g}(t)}$ at time $t$.

To compute this distribution, henceforth called pcLNA distribution and written

$$P_{\text{pcLNA}}(\boldsymbol{Q}_{\boldsymbol{x}} | \boldsymbol{X}(0)),$$

suppose that $\boldsymbol{\xi}(0) \sim MVN(\boldsymbol{m}_0, \boldsymbol{S}_0)$ and use the results in section 4.1 to compute $P(\boldsymbol{X}(t) | \boldsymbol{\xi}(0) \sim MVN(\boldsymbol{m}_0, \boldsymbol{S}_0))$ which is $MVN(\boldsymbol{\mu}_t, \boldsymbol{\Sigma}_t)$ with parameters as in (27) in the original coordinates (that correspond to concentration levels of each of the reacting molecules). Then, find an orthogonal matrix $\boldsymbol{R}_x = [\boldsymbol{R}_x^{(1)}\ \boldsymbol{R}_x^{(2)}]$ such that $\boldsymbol{R}_x^T(\boldsymbol{X}(t) - \boldsymbol{x}) = (X^{(1)}(t), \boldsymbol{X}^{(2)}(t))^T$ with $\boldsymbol{X}^{(2)}(t) \in \mathcal{S}_{x=\boldsymbol{g}(t)}$. The pcLNA distribution $P_{\text{pcLNA}}(\boldsymbol{Q}_{\boldsymbol{x}}|\boldsymbol{\xi}(0) \sim MVN(\boldsymbol{m}_0, \boldsymbol{S}_0))$ is $MVN(\boldsymbol{\mu}_{\boldsymbol{x}}, \boldsymbol{\Sigma}_{\boldsymbol{x}})$, where

$$\boldsymbol{\mu}_{\boldsymbol{x}} = \Omega^{-1/2}\boldsymbol{m}_t^{(2)}, \qquad \boldsymbol{\Sigma}_x = \Omega^{-1}\boldsymbol{V}_t^{(2,2)},$$

and the moments of the noise process on the transversal are

$$\boldsymbol{m}_t^{(2)} = \boldsymbol{R}_x^{(2)^T}\boldsymbol{m}_t \qquad \boldsymbol{V}_t^{(2,2)} = \boldsymbol{R}_x^{(2)^T}\boldsymbol{V}_t\boldsymbol{R}_x^{(2)},$$

with $\boldsymbol{m}_t$, $\boldsymbol{V}_t$ given in (26). This pcLNA distribution is compared with the empirical distribution of transversal points derived from SSA simulations as explained in the previous section. The results are presented in **I** Figure 4 and demonstrate that the two distributions are hardly distinguishable.

Note that the above pcLNA distributions are slightly simpler than those presented in [13]. Here the marginal LNA distributions of $\boldsymbol{X}^{(2)}$ are used instead of the conditional distributions $(\boldsymbol{X}^{(2)}|X^{(1)} = 0)$. The performance of the two distributions compared to the SSA simulations are very similar with differences identified only during the initial transient response of NF-$\kappa$B to TNF$\alpha$ signals. The simpler marginal distributions provided here give slightly better fit during this period due to their extra variability that fits better the SSA simulations. The properties of pcLNA distributions, in terms of accuracy compared to SSA, robustness as $t \to \infty$, and analytical tractability are thoroughly discussed in [13].

##### 4.2.2 The pcLNA joint distribution of multiple transversals

We now wish to approximate the joint distribution of multiple transversal points $P(\mathbf{Q}_{1:k}|\boldsymbol{X}(0) \sim MVN(\boldsymbol{\mu}_0, \boldsymbol{\Sigma}_0)) = P(\boldsymbol{Q}_{\boldsymbol{x}_1}, \boldsymbol{Q}_{\boldsymbol{x}_2}, \ldots, \boldsymbol{Q}_{\boldsymbol{x}_k}|\boldsymbol{X}(0) \sim MVN(\boldsymbol{\mu}_0, \boldsymbol{\Sigma}_0))$ where $\boldsymbol{x}_i = \boldsymbol{g}(t_i)$ and $0 \le t_1 \le \cdots \le t_k$. This can be written as a product of transition probabilities

$$P(\mathbf{Q}_{1:k}|\boldsymbol{X}(0)) = P(\boldsymbol{Q}_{\boldsymbol{x}_1}, \boldsymbol{Q}_{\boldsymbol{x}_2}, \ldots, \boldsymbol{Q}_{\boldsymbol{x}_k}|\boldsymbol{X}(0)) = P(\boldsymbol{Q}_{\boldsymbol{x}_1}|\boldsymbol{X}(0)) \prod_{i=1}^{k-1} P(\boldsymbol{Q}_{\boldsymbol{x}_{i+1}}|\boldsymbol{Q}_{\boldsymbol{x}_i})$$

The first transition probability on the rhs of the above equation, as explained in the previous section, is approximated by the $MVN(\boldsymbol{\mu}_{\boldsymbol{x}_1}, \boldsymbol{\Sigma}_{\boldsymbol{x}_1})$ distribution, by taking $\boldsymbol{\mu}_0 = \boldsymbol{x}_0 + \Omega^{-1/2}\boldsymbol{m}_0$, $\boldsymbol{\Sigma}_0 = \boldsymbol{S}_0/\Omega$ and $\boldsymbol{x} = \boldsymbol{x}_1 = \boldsymbol{g}(t_1)$. To derive the approximation of the transition probabilities from $\boldsymbol{Q}_{\boldsymbol{x}_i}$ to $\boldsymbol{Q}_{\boldsymbol{x}_{i+1}}$ note that

$$\begin{aligned}
\boldsymbol{X}^{(2)}(t_{i+1}) &= \boldsymbol{R}_{\boldsymbol{x}_{i+1}}^{(2)^T}(\boldsymbol{X}(t_{i+1}) - \boldsymbol{x}(t_{i+1})), \quad \text{by (28)} \\
&= \boldsymbol{R}_{\boldsymbol{x}_{i+1}}^{(2)^T}\Omega^{-1/2}\boldsymbol{\xi}(t_{i+1}), \quad \text{by (20)} \\
&= \boldsymbol{R}_{\boldsymbol{x}_{i+1}}^{(2)^T}\Omega^{-1/2}(\boldsymbol{C}(t_i, t_{i+1})\boldsymbol{\xi}(t_i) + \boldsymbol{\eta}(t_i, t_{i+1})), \quad \text{by (23)} \\
&= \Omega^{-1/2}(\boldsymbol{R}_{\boldsymbol{x}_{i+1}}^{(2)^T}\boldsymbol{\eta}(t_i, t_{i+1}) + \boldsymbol{R}_{\boldsymbol{x}_{i+1}}^{(2)^T}\boldsymbol{C}(t_i, t_{i+1})(\boldsymbol{X}(t_i) - \boldsymbol{x}(t_i))), \quad \text{by (20)} \\
&= \Omega^{-1/2}(\boldsymbol{R}_{\boldsymbol{x}_{i+1}}^{(2)^T}\boldsymbol{\eta}(t_i, t_{i+1}) + \boldsymbol{R}_{\boldsymbol{x}_{i+1}}^{(2)^T}\boldsymbol{C}(t_i, t_{i+1})(\boldsymbol{R}_{\boldsymbol{x}_i}^{(1)}X^{(1)}(t_i) + \boldsymbol{R}_{\boldsymbol{x}_i}^{(2)}\boldsymbol{X}^{(2)}(t_i))), \quad \text{by (28).}
\end{aligned}$$

We assume that the term describing the transition through $\boldsymbol{C}(t_i, t_{i+1})$ of the tangental fluctuations at time $t_i$, $\boldsymbol{R}_{\boldsymbol{x}_i}^{(1)}X^{(1)}(t_i)$, to fluctuations on the transversal coordinate, $\boldsymbol{R}_{\boldsymbol{x}_{i+1}}^{(2)}$, at time $t_{i+1}$, i.e.

$$\Omega^{-1/2}\boldsymbol{R}_{\boldsymbol{x}_{i+1}}^{(2)^T}\boldsymbol{C}(t_i, t_{i+1})\boldsymbol{R}_{\boldsymbol{x}_i}^{(1)}X^{(1)}(t_i)$$

is small compared to the transition of transversal fluctuations and ignore it to derive that $P_{\text{pcLNA}}(\boldsymbol{Q}_{\boldsymbol{x}_{i+1}}|\boldsymbol{Q}_{\boldsymbol{x}_i}=q_i) = P_{\text{LNA}}(\boldsymbol{X}^{(2)}(t_{i+1})|\boldsymbol{X}^{(2)}(t_i)=q_i)$ is $MVN\left(\hat{\boldsymbol{\mu}}_{\boldsymbol{x}_{i+1}}, \hat{\boldsymbol{\Sigma}}_{\boldsymbol{x}_{i+1}}\right)$, where

$$\hat{\boldsymbol{\mu}}_{\boldsymbol{x}_{i+1}} = \boldsymbol{C}^{(2,2)}(t_i,t_{i+1})q_i, \qquad \boldsymbol{C}^{(2,2)}(t_i,t_{i+1}) = \boldsymbol{R}^{(2)^T}_{\boldsymbol{x}_{i+1}}\boldsymbol{C}(t_i,t_{i+1})R^{(2)}_{\boldsymbol{x}_i},$$

and

$$\hat{\boldsymbol{\Sigma}}_{\boldsymbol{x}_{i+1}} = \Omega^{-1}\boldsymbol{V}^{(2,2)}(t_i,t_{i+1})), \qquad \boldsymbol{V}^{(2,2)}(t_i,t_{i+1}) = \boldsymbol{R}^{(2)^T}_{\boldsymbol{x}_{i+1}}\boldsymbol{V}(t_i,t_{i+1}))\boldsymbol{R}^{(2)}_{\boldsymbol{x}_{i+1}}.$$

More generally, the pcLNA transition probability with $\boldsymbol{X}^{(2)}(t_i) \sim MVN(\boldsymbol{\mu}_{\boldsymbol{x}_i}, \boldsymbol{\Sigma}_{\boldsymbol{x}_i})$,

$$P_{\text{pcLNA}}\left(\boldsymbol{Q}_{\boldsymbol{x}_{i+1}}|\boldsymbol{Q}_{\boldsymbol{x}_i} \sim MVN(\boldsymbol{\mu}_{\boldsymbol{x}_i}, \boldsymbol{\Sigma}_{\boldsymbol{x}_i})\right) = P_{\text{LNA}}\left(\boldsymbol{X}^{(2)}(t_{i+1})\middle|\boldsymbol{X}^{(2)}(t_i) \sim MVN(\boldsymbol{\mu}_{\boldsymbol{x}_i}, \boldsymbol{\Sigma}_{\boldsymbol{x}_i})\right)$$

is $\text{MVN}(\boldsymbol{\mu}_{\boldsymbol{x}_{i+1}}, \boldsymbol{\Sigma}_{\boldsymbol{x}_{i+1}})$ where (again ignoring the tangental fluctuations transduced on the transversal)

$$\boldsymbol{\mu}_{\boldsymbol{x}_{i+1}} = \boldsymbol{C}^{(2,2)}(t_i,t_{i+1})\boldsymbol{\mu}_{\boldsymbol{x}_i},$$

and

$$\boldsymbol{\Sigma}_{\boldsymbol{x}_{i+1}} = \hat{\boldsymbol{\Sigma}}_{\boldsymbol{x}_{i+1}} + \boldsymbol{C}^{(2,2)}(t_i,t_{i+1})\boldsymbol{\Sigma}_{\boldsymbol{x}_i}\boldsymbol{C}^{(2,2)}(t_i,t_{i+1})^T.$$

Then, the pcLNA joint distribution is

$$P_{\text{pcLNA}}(\boldsymbol{Q}_{\boldsymbol{x}_2}, \boldsymbol{Q}_{\boldsymbol{x}_1}|\boldsymbol{X}(0) \sim \text{MVN}(\boldsymbol{\mu}_0, \boldsymbol{\Sigma}_0)) \propto \exp\left(-\frac{1}{2}\left\{(\boldsymbol{Q}_{\boldsymbol{x}_1} - \boldsymbol{\mu}_{\boldsymbol{x}_1})^T\boldsymbol{\Sigma}^{-1}_{\boldsymbol{x}_1}(\boldsymbol{Q}_{\boldsymbol{x}_1} - \boldsymbol{\mu}_{\boldsymbol{x}_1}) \right.\right.$$
$$\left.\left. + \left(\boldsymbol{Q}_{\boldsymbol{x}_2} - \boldsymbol{C}^{(2,2)}(t_1,t_2)\boldsymbol{Q}_{\boldsymbol{x}_1}\right)^T\hat{\boldsymbol{\Sigma}}_{\boldsymbol{x}_2}\left(\boldsymbol{Q}_{\boldsymbol{x}_2} - \boldsymbol{C}^{(2,2)}(t_1,t_2)\boldsymbol{Q}_{\boldsymbol{x}_1}\right)\right\}\right)$$

and by adding and subtracting the term $\boldsymbol{C}^{(2,2)}(t_1,t_2)\boldsymbol{\mu}_{\boldsymbol{x}_1}$ in the second term of the rhs above, we have that this can be written as the kernel of a $MVN$ distribution with mean vector

$$(\boldsymbol{\mu}_{\boldsymbol{x}_1}, \boldsymbol{\mu}_{\boldsymbol{x}_2})$$

and precision matrix (inverse of covariance matrix)

$$\begin{pmatrix} \boldsymbol{\Sigma}^{-1}_{\boldsymbol{x}_1} + \boldsymbol{C}^{(2,2)}(t_1,t_2)^T\hat{\boldsymbol{\Sigma}}^{-1}_{\boldsymbol{x}_2}\boldsymbol{C}^{(2,2)}(t_1,t_2) & -\boldsymbol{C}^{(2,2)^T}(t_1,t_2))\hat{\boldsymbol{\Sigma}}^{-1}_{\boldsymbol{x}_2} \\ -\hat{\boldsymbol{\Sigma}}^{-1}_{\boldsymbol{x}_2}\boldsymbol{C}^{(2,2)}(t_1,t_2) & \hat{\boldsymbol{\Sigma}}^{-1}_{\boldsymbol{x}_2} \end{pmatrix}.$$

By induction we can derive that the pcLNA joint distributions $P_{\text{pcLNA}}(\boldsymbol{Q}_{\boldsymbol{x}_1}, \boldsymbol{Q}_{\boldsymbol{x}_2}, \dots, \boldsymbol{Q}_{\boldsymbol{x}_k}|\boldsymbol{X}(0) \sim MVN(\boldsymbol{\mu}_0, \boldsymbol{\Sigma}_0))$, for $k = 2, \dots,$ are $MVN(\boldsymbol{\mu}_{\boldsymbol{x}_{1:k}}, A^{-1}_{\boldsymbol{x}_{1:k}})$ with mean vector

$$\boldsymbol{\mu}_{\boldsymbol{x}_{1:k}} = (\boldsymbol{\mu}_{\boldsymbol{x}_1}, \boldsymbol{\mu}_{\boldsymbol{x}_2}, \dots, \boldsymbol{\mu}_{\boldsymbol{x}_k})$$

and precision matrix, $A_{\boldsymbol{x}_{1:k}}$, with a block tridiagonal form where the diagonal blocks, $A^{(ii)}_{\boldsymbol{x}_{1:k}}$, are

$$\boldsymbol{\Sigma}^{-1}_{\boldsymbol{x}_i} + (\boldsymbol{C}^{(2,2)}(t_i,t_{i+1}))^T(\hat{\boldsymbol{\Sigma}}_{\boldsymbol{x}_{i+1}})^{-1}\boldsymbol{C}^{(2,2)}(t_i,t_{i+1}), \qquad \text{for } i = 1, 2, \dots, k-1$$

and

$$\hat{\boldsymbol{\Sigma}}^{-1}_{\boldsymbol{x}_{i+1}}, \qquad \text{for } i = k,$$

upper diagonal entries, $\boldsymbol{A}^{(i,i+1)}_{\boldsymbol{x}_{1:k}}$, $i = 1, 2, \dots, k-1$,

$$-\boldsymbol{C}^{(2,2)}(t_i,t_{i+1})^T\hat{\boldsymbol{\Sigma}}^{(2,2)^{-1}}_{t_{i+1}}$$

and lower diagonal entries, $\boldsymbol{A}^{(i+1,i)}_{\boldsymbol{x}_{1:k}}$, $i = 1, 2, \dots, k-1$,

$$-\hat{\boldsymbol{\Sigma}}^{(2,2)^{-1}}_{t_{i+1}}\boldsymbol{C}^{(2,2)}(t_i,t_{i+1}).$$

To write the pcLNA joint distribution in the original coordinates we can use the transformation $\boldsymbol{x}_{1:k} + \boldsymbol{R}_{1:k}\boldsymbol{Q}_{1:k}$ where $\boldsymbol{x}_{1:k} = (x_1, \boldsymbol{x}_2, \ldots, \boldsymbol{x}_k)^T$, $\boldsymbol{R}_{1:k} = \text{Diag}\left(\boldsymbol{R}^{(2)}_{t_1}, \ldots, \boldsymbol{R}^{(2)}_{t_k})\right)$ to get the mean vector

$$\boldsymbol{\mu}_{1:k} = \boldsymbol{x}_{1:k} + \boldsymbol{R}_{1:k}\boldsymbol{\mu}_{\boldsymbol{x}_{1:k}} \tag{29}$$

and precision matrix

$$\boldsymbol{A}_{1:k} = \boldsymbol{R}_{1:k}\boldsymbol{A}_{\boldsymbol{x}_{1:k}}\boldsymbol{R}^T_{1:k}. \tag{30}$$

As discussed above, the pcLNA joint distributions have been shown to have hardly distinguishable distributions to the empirical transversal distributions derived by SSA in the NF-$\kappa$B system (see Figure 4 in **I** and [13]).

#### 4.3 pcLNA simulation

The approach is to amend the LNA Ansatz $\boldsymbol{X}(t) = \boldsymbol{g}(t) + \Omega^{-1/2}\boldsymbol{\xi}(t)$ to $\boldsymbol{X}(t) = \boldsymbol{g}(s) + \Omega^{-1/2}\boldsymbol{\kappa}(s)$ where $\boldsymbol{g}(s) = \boldsymbol{G}_N(\boldsymbol{X}(t))$ and to use resetting of $t$ to $s$ to cope with the growth in the variance of phase deviations keeping the LNA fluctuation $\boldsymbol{\kappa}(s)$ normal to $\gamma$. While for free-running oscillators the variance of $\boldsymbol{\xi}(t)$ grows without bound as $t$ increases, $\boldsymbol{\kappa}(s)$ has uniformly bounded variance.

The pcLNA simulation algorithm iteratively uses standard LNA steps of length $\Delta\tau$ to move from a state $\boldsymbol{X}(s_{i-1})$ to a new state $\boldsymbol{X}(s_{i-1} + \Delta\tau) = \boldsymbol{X}_i$, $i = 1, 2, \ldots$. After each LNA step, the phase of the system is reset or "corrected" such that $\boldsymbol{g}(s_i) = \boldsymbol{G}_N(\boldsymbol{X}_i)$ and the (global) fluctuations $\boldsymbol{\xi}(s_{i-1} + \Delta\tau) = \Omega^{-1/2}(\boldsymbol{X}_i - \boldsymbol{g}(s_{i-1} + \Delta\tau))$ are replaced by the normally transversal fluctuation $\boldsymbol{\kappa}(s_i) = \Omega^{-1/2}(\boldsymbol{X}_i - \boldsymbol{g}(s_i))$ which are MVN distributed and, as we showed in the previous section, approximate well the transversal fluctuations under the exact Markov Jump process.

The steps of the pcLNA simulation algorithm are described next in more detail (see also Fig 7).

1. Choose a time-step size $\Delta\tau > 0$.
2. Input **initial conditions** $\boldsymbol{\kappa}(\boldsymbol{S}_0)$ and $\boldsymbol{X}_0 = \boldsymbol{g}(\boldsymbol{S}_0) + \Omega^{-1/2}\boldsymbol{\kappa}(\boldsymbol{S}_0)$.
3. **For iteration** $i = 1, 2, \ldots$
   (a) sample $\boldsymbol{\xi}(s_{i-1} + \Delta\tau)$ from MVN$(\boldsymbol{C}_i\boldsymbol{\kappa}(s_{i-1}), V_i)$;
   (b) compute $\boldsymbol{X}_i = \boldsymbol{g}(s_{i-1} + \Delta\tau) + \Omega^{-1/2}\boldsymbol{\xi}(s_{i-1} + \Delta\tau)$;
   (c) set $s_i$ to be such that $\boldsymbol{G}_N(\boldsymbol{X}_i) = \boldsymbol{g}(s_i)$ and $\boldsymbol{\kappa}_i = \Omega^{1/2}(\boldsymbol{X}_i - \boldsymbol{g}(s_i))$.

In the for loop of step 3, $\boldsymbol{C}_i = \boldsymbol{C}(s_{i-1}, s_{i-1} + \Delta\tau)$ and $\boldsymbol{V}_i = \boldsymbol{V}(s_{i-1}, s_{i-1} + \Delta\tau)$ that are derived as solutions of (24) and (25). The simulated sample $\boldsymbol{X}_i$ corresponds to time $t_i = t_0 + i\Delta\tau$, $i = 1, 2, \ldots$, where $t_0$ is the initial time. The time $t_i$ is not necessarily equal to the phase $s_i$, defined by the relation $\boldsymbol{g}(s_i) = \boldsymbol{G}_N(\boldsymbol{X}_i)$, which is stochastic and has variance linearly increasing with the time step $\Delta\tau$.

If one wants to record simulated trajectories at a finer time-scale than $\Delta\tau$ then one can run the algorithm with $\Delta\tau$ replaced by $\Delta\tau/M$ for some integer $M > 1$ and only carry out the phase correction in step 3(c) every $M$th step and at all the other steps just proceeding as in the standard LNA (ignoring step 3(c)). This gives the same distribution as if the intermediate points had not been calculated because of the transitive nature of the LNA i.e. the distribution $P_{\text{LNA}}(\boldsymbol{X}(s+t)|\boldsymbol{X}(0))$ is equal to the distribution $P_{\text{LNA}}(\boldsymbol{X}(t)|\boldsymbol{X}(s) \sim P_{\text{LNA}}(\boldsymbol{X}(s)|\boldsymbol{X}(0)))$. We have noticed that roughly 3-5 corrections per peak-to-peak time typically give good balance between speed and precision.

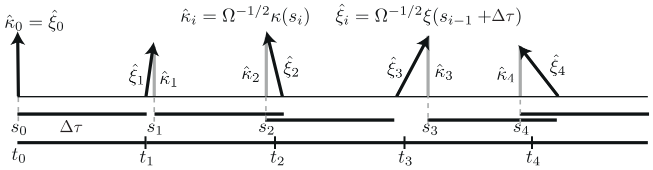

**Figure S6: The main steps in the pcLNA simulation algorithm.** *The solid horizontal bars below the horizontal axis are all of length $\Delta\tau$, the basic time step of the algorithm. The black arrows show $\hat{\boldsymbol{\xi}}_i = \Omega^{-1/2}\boldsymbol{\xi}(s_{i-1} + \Delta\tau)$ and the grey arrows $\hat{\boldsymbol{\kappa}}_i = \Omega^{-1/2}\boldsymbol{\kappa}(s_i)$. Having calculated $\hat{\boldsymbol{\kappa}}(s_{i-1})$ one uses $\boldsymbol{\kappa}(s_{i-1})$ as the initial state and updates it using the LNA and a time-step $\Delta\tau$ to obtain $\boldsymbol{\xi}$ at $s_{i-1} + \Delta t$. Then $\boldsymbol{\xi}(s_{i-1} + \Delta\tau)$ is replaced by $\boldsymbol{\kappa}(s_i)$ so that $\boldsymbol{g}(s_{i-1} + \Delta\tau) + \Omega^{-1/2}\boldsymbol{\xi}(s_{i-1} + \Delta\tau) = \boldsymbol{g}(s_i) + \Omega^{-1/2}\boldsymbol{\kappa}(s_i)$ where $\boldsymbol{\kappa}(s_i)$ is normal to the limit cycle. Therefore, $s_i$ gives the phase of $\boldsymbol{\kappa}(s_i)$ and the corresponding time is $t_i = t_0 + i\,\Delta\tau$.*

#### 4.4 Channel capacity estimation

In our comparisons to the experimental results of the NF-$\kappa$B system response to TNF$\alpha$ stimulus in the section "Mutual Information" and Figure 4(d) of **I**, we allowed the total number of NF-$\kappa$B molecules to vary in different cells-simulated trajectories. Specifically, at the beginning of the simulation of each trajectory, we first generated the total number of NF-$\kappa$B molecules from a log-Normal distribution that is previously used in [16]. This has pdf

$$f(x) = \frac{A}{x\sigma\sqrt{2\pi}} \exp\left(-\frac{(\log(x) - \boldsymbol{\mu})^2}{2\sigma^2}\right), \qquad x > 0$$

and the parameters

$$A = 10^5,\ \sigma = 1/\sqrt{2},\ \boldsymbol{\mu} = -1/4$$

are chosen to give a mean number of molecules equal to $10^5$ that agrees with the fixed value assumed in the deterministic model used in [2] as well as the rest of the models used here. The log-Normal is chosen to give a positive support (i.e. only positive number of the total number of molecules are possible), a single mode value and tunable variance.

After generating the number of NF-$\kappa$B molecules, we used the pcLNA simulation algorithm to obtain a stochastic trajectory. We initiated the stochastic simulation from the equilibrium point under no dose and immediately applied a continuous TNF$\alpha$ dose. We then obtained the nuclear NF-$\kappa$B concentration at $t = 30$min.

The dose was assumed to have a discrete distribution with weights obtained using a constrained optimisation performed by the fmincon function in MATLAB [11].

We show the results assuming that the TNF$\alpha$ dose has only two levels (high and low), but the use of medium doses did not change the estimation substantially.

#### 4.5 Channel capacity under the stochastic model of NF-$\kappa$B in Tay et al. [16]

We used the code provided by [16] to run 500 simulations over a high (10ng/mL) and a low (0.5ng/mL) TNF dose. Similarly to above, we collected the nuclear concentration of NF-$\kappa$B at $t = 30min$ after the initiation of the continuous TNF$\alpha$ stimulus. The histograms are provided in Figure S7. The channel capacity was estimated to be 0.72 assuming that the dose has a discrete distribution. The optimal weights of the dose distribution were derived using a constrained optimisation performed by the fmincon function in MATLAB [11].

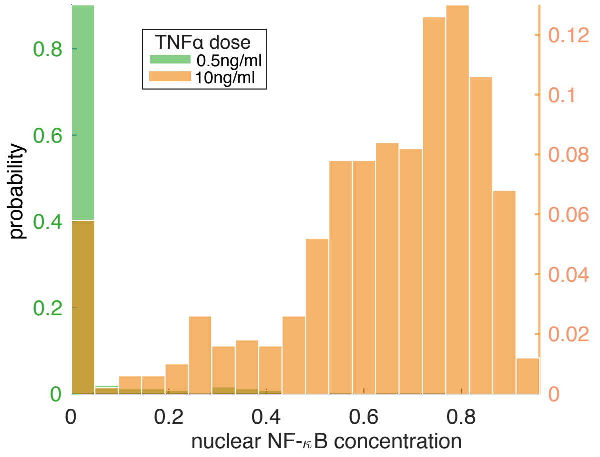

**Figure S7: Simulation of nuclear NF-$\kappa$B concentration using the model in [16]** *Histograms of the $n = 500$ stochastic simulations computed using the code provided in [16] at time $t = 30min$ after the beginning of continuous TNF$\alpha$ stimulation.*

#### 4.6 Simulation of EGR1 and COX-2 genes.

Here we explain the stochastic simulation of copies of mRNA molecules of the EGR1 and COX-2 genes as targets of the NF-$\kappa$B. As explained in **I**, we use a similar model as the one described in SI section 3.4, the difference being that the modification is now promoted by the active state of IKK. More specifically, in equation 12 in table S6 the term $p_{a1}S_2\frac{N_c}{N_c+k_{pnf}}$ is replaced with $p_{a1} \times K_a \times S_2\frac{N_c}{N_c+k_{pnf}}$ and the parameters $p_{a1} = 0.0025$, $S_2 = 1$, $k_{pnf} = 0.01$. The reverse modification parameters are also slightly changed to $k_{pnfp} = 0.01$, $p_{d1} = 1 \times 10^{-5}$. Because the model does not include any feedback of these genes to the NF-$\kappa$B system, they can be simulated separately to the base system, and this makes the hybrid algorithm explained below somewhat simpler than hybrid algorithms such as those discussed for example in [17]. We therefore used the pcLNA simulation algorithm to derive stochastic trajectories of the base system and then applied the SSA to generate stochastic simulations for the EGR1 and COX-2 genes. The reason for using SSA in the latter simulation rather than LNA or pcLNA is that EGR1 and COX-2 activation is highly stochastic and not oscillatory. This method proved to be fairly efficient for the purposes of our demonstration, as we exploited the computational advantages of using pcLNA in the base system, while SSA simulations of only two genes expressed in similar scales proved to also be fairly efficient.

The algorithm proceeds as follows. Let $Y^F = (Y^F(0), Y^F(t_1), \dots, Y^F(t_n))$ be a trajectory of the TF (in molecule numbers) derived using an appropriate stochastic simulation algorithm (e.g. pcLNA). Define the continuous-time function $Y^F(t)$, $t \geq 0$ of the observed at discrete times $Y^F$ using an appropriate (smooth) interpolation method. Also define the degradation, $u_{degr}(\delta, Y^g)$, and transcription, $u_{tran}(\tau, Y^F)$, rate functions respectively. The algorithm proceeds as follows:

1. initialise the gene expression $Y^g := Y^g(0)$ and set $t := 0$,
2. generate the next time of degradation event, $t_1 \sim t + \text{Exp}(u_{degr}(\delta, Y^g))$,
3. generate $u \sim U([0, 1])$ and integrate the transcription rate function up to last time $t_2$ for which $I = \int_t^{t_2} u_{tran}(\tau, Y^F(s))ds + \log(u) \leq 0$.
4. Set $t := (t_1 \wedge t_2)$ and

$$Y^g := \begin{cases} Y^g - 1, & \text{if} \quad t = t_1, \\ Y^g + 1, & \text{otherwise} \end{cases}$$

5. If $t < T_{\max}$ return to step 2.

Note that a Gibson-Bruck type of this algorithm (see [17, 18]) could be used alternatively, but will make little difference for the purposes of our demonstration.

In our simulation, the degradation reaction rate of EGR-1 and COX-2 are

$$u_{degr,g_1} = \delta_1 Y^{(g_1)}, \qquad u_{degr,g_2} = (\Omega \times \delta_2)\frac{Y^{(g_2)}}{Y^{(g_2)} + k_{g_2}}$$

and the transcription rates

$$u_{tr} = (\Omega \times \tau)\frac{Y_A^h}{Y_A^h + k_A^h}\frac{k_R^h}{Y_R^h + k_R^h}$$

where $Y_A$ the state of the activator TF and $Y_R$ the state of the inhibitor TF. As shown in **I** Fig. 4(a), the activator of EGR1 is the (unmodified) nuclear NF-$\kappa$B , $N_n$, and the inhibitor its modified form, $(N_m)_n$ and for COX-2 vice versa. The parameter values used for EGR1 are $\delta_1 = 1 \times 10^{-3}$, $\tau = 8 \times 10^{-7}$, $k_A = 0.03$, $k_R = 0.03$ and for COX-2 are $\delta_2 = 5 \times 10^{-6}$, $\tau = 2 \times 10^{-7}$, $k_A = 0.015$, $k_R = 0.005$, $k_{g_2} = 1 \times 10^{-3}$, also $h = 5$.

##### 4.7 Fisher Information Matrix (FIM) under the pcLNA

To compute FIM under the pcLNA, we need to compute derivatives of the solution $\boldsymbol{x} = \boldsymbol{g}(t)$, and the matrices $\boldsymbol{C}(t_0, t)$, $\boldsymbol{V}(t_0, t)$ with respect to the parameter vector $\boldsymbol{\theta}$.

###### 4.7.1 Derivatives for FIM computation

Consider a solution $\boldsymbol{x} = \boldsymbol{\eta}(t; t^*, \boldsymbol{x}^*, \boldsymbol{\theta})$, $t \geq 0$, of the ode in (19) with initial condition $\boldsymbol{\eta}(t^*; t^*, \boldsymbol{\theta}) = \boldsymbol{x}^*$. Here we write the solution as $\boldsymbol{\eta}(t; t^*, \boldsymbol{x}^*, \boldsymbol{\theta})$ (and the tangent vector as $F(\boldsymbol{\eta}(t); \boldsymbol{\theta})$) to emphasise its dependence to the initial conditions $t^*$, $\boldsymbol{x}^* = \left(x_1^*, x_2^*, \ldots, x_n^*\right)^T$ and the parameter vector $\boldsymbol{\theta} = \left(\theta_1, \theta_2, \ldots, \theta_k\right)^T$.

The Jacobian matrix $\boldsymbol{J}(\boldsymbol{x}) = \boldsymbol{J}(\boldsymbol{x}; \boldsymbol{\theta})$ at $\boldsymbol{x} = \boldsymbol{\eta}(t; t^*, \boldsymbol{x}^*, \boldsymbol{\theta})$ has entries

$$J_{ii'} = \left(\frac{\vartheta \boldsymbol{F}}{\vartheta \boldsymbol{x}}\right)_{i,i'} = \frac{\vartheta \boldsymbol{F}_i}{\vartheta \boldsymbol{x}_{i'}}.$$

Define the first and second derivatives of the solution $\boldsymbol{x} = \boldsymbol{\eta}(t; t^*, \boldsymbol{x}^*, \boldsymbol{\theta})$ with respect to the initial conditions and parameters

$$\boldsymbol{u}_k = \frac{\vartheta \boldsymbol{\eta}}{\vartheta \boldsymbol{x}_k^*}, \qquad \boldsymbol{v}_j = \frac{\vartheta \boldsymbol{\eta}}{\vartheta \theta_j}, \qquad \boldsymbol{w}_{jk} = \frac{\vartheta \boldsymbol{v}_j}{\vartheta \boldsymbol{x}_k^*} = \frac{\vartheta \boldsymbol{u}_k}{\vartheta \theta_j} = \frac{\vartheta^2 \boldsymbol{\eta}}{\vartheta \boldsymbol{x}_k^* \vartheta \theta_j}. \tag{31}$$

As we demonstrate below, the first derivative provides the columns of the fundamental matrix $\boldsymbol{C}(t^*, t)$. The second provides the derivatives of the ode solution wrt to the parameters. The third provides the derivatives of the entries of the fundamental matrix wrt to the $j$-th parameter. All three are useful in computing the derivatives required for the FIM computation.

The derivative $u_k$ in (31) is the solution of the initial value problem

$$\boldsymbol{x}' = \boldsymbol{J}(t)\boldsymbol{x}, \quad \boldsymbol{x}(t^*) = \boldsymbol{e}_k,$$

where $\boldsymbol{e}_l = 0$, for $l \neq k$, $\boldsymbol{e}_l = 1$, for $l = k$.

**Proof.**

$$
\begin{aligned}
\frac{d\boldsymbol{\eta}}{dt} &= F(\boldsymbol{\eta};\boldsymbol{\theta}) \\
\frac{d}{d\boldsymbol{x}_k^*}\left(\frac{d\boldsymbol{\eta}}{dt}\right) &= \frac{dF}{d\boldsymbol{x}_k^*}(\boldsymbol{\eta},\boldsymbol{\theta}) \\
\frac{d}{dt}\left(\frac{\vartheta\boldsymbol{\eta}}{\vartheta\boldsymbol{x}_k^*}\right) &= \frac{\vartheta F}{\vartheta\boldsymbol{\eta}}\frac{\vartheta\boldsymbol{\eta}}{\vartheta\boldsymbol{x}_k^*} \\
\frac{d\boldsymbol{u}_k}{dt} &= \boldsymbol{J}(t)\boldsymbol{u}_k
\end{aligned}
$$

and

$$
\boldsymbol{u}_k(t^*) = \frac{\vartheta\boldsymbol{\eta}}{\vartheta\boldsymbol{x}_k^*}(t^*) = \frac{\vartheta\boldsymbol{x}^*}{\vartheta\boldsymbol{x}_k^*} = \boldsymbol{e}_k.\square
$$

Note that the principal fundamental matrix $\boldsymbol{C}$ is the solution of the latter initial value problem. The matrix can be written as

$$
\boldsymbol{C}(t^*,t) = (u_1(t), u_2(t), \ldots, u_n(t)).
$$

Furthermore, $\boldsymbol{v}_j$ in (31) is the solution of the initial value problem

$$
\boldsymbol{x}' = \boldsymbol{J}(t)\boldsymbol{x} + \boldsymbol{h}_j(t), \quad \boldsymbol{x}(t^*) = \boldsymbol{0} \tag{32}
$$

where $\boldsymbol{h}_j(t) = \frac{\vartheta\boldsymbol{F}}{\vartheta\boldsymbol{\theta}_j}$ (note: this is the partial derivative of $\boldsymbol{F}$ to $\boldsymbol{\theta}$, i.e. $\boldsymbol{x}$ is taken as constant to $\boldsymbol{\theta}$).

**Proof.**

$$
\begin{aligned}
\frac{d\boldsymbol{\eta}}{dt} &= \boldsymbol{F}(\boldsymbol{\eta},\boldsymbol{\theta}) \\
\frac{d}{d\theta_j}\left(\frac{d\boldsymbol{\eta}}{dt}\right) &= \frac{d\boldsymbol{F}}{d\theta_j} \\
\frac{d}{dt}\left(\frac{\vartheta\boldsymbol{\eta}}{\vartheta\theta_j}\right) &= \frac{\vartheta\boldsymbol{F}}{\vartheta\boldsymbol{\eta}}\frac{\vartheta\boldsymbol{\eta}}{\vartheta\theta_j} + \frac{\vartheta\boldsymbol{F}}{\vartheta\theta_j}\frac{\vartheta\theta_j}{\vartheta\theta_j} \\
\frac{d\boldsymbol{v}_j}{dt} &= \boldsymbol{J}\boldsymbol{v}_j + \boldsymbol{h}_j(t).
\end{aligned}
$$

Also,

$$
\boldsymbol{v}_j(t^*) = \frac{\vartheta\boldsymbol{\eta}}{\vartheta\theta_j}(t^*) = \frac{\vartheta\boldsymbol{x}^*}{\vartheta\theta_j} = \boldsymbol{0}\square
$$

Furthermore, $\boldsymbol{w}_{jk}$ in (31) can be derived as the solution of the initial value problem,

$$
\frac{d\boldsymbol{x}}{dt} = \boldsymbol{J}\boldsymbol{x} + \left(\frac{\vartheta J_{kk'}}{\vartheta\boldsymbol{\eta}}v_j + \frac{\vartheta J_{kk'}}{\vartheta\theta_j}\right)_{k,k'}\boldsymbol{u}_k, \qquad \boldsymbol{x}(t^*) = \boldsymbol{0} \tag{33}
$$

**Proof.**

$$
\begin{aligned}
\frac{d}{d\theta_j}\frac{d\boldsymbol{u}_k}{dt} &= \frac{d}{d\theta_j}\left(\boldsymbol{J}(\boldsymbol{\eta},\boldsymbol{\theta})\boldsymbol{u}_k\right) \\
\frac{d}{dt}\frac{d\boldsymbol{u}_k}{d\theta_{j)}} &= \frac{d\boldsymbol{J}(\boldsymbol{\eta},\boldsymbol{\theta})}{d\theta_j}\boldsymbol{u}_k + \boldsymbol{J}(\boldsymbol{\eta},\boldsymbol{\theta})\frac{d\boldsymbol{u}_k}{d\boldsymbol{\theta}_j} \\
\frac{d\boldsymbol{w}_{jk}}{dt} &= \frac{d\boldsymbol{J}(\boldsymbol{\eta},\boldsymbol{\theta})}{d\boldsymbol{\theta}_j}\boldsymbol{u}_k + \boldsymbol{J}(\boldsymbol{\eta},\boldsymbol{\theta})\boldsymbol{w}_{jk}
\end{aligned}
$$

where

$$\frac{d\boldsymbol{J}(\boldsymbol{\eta},\boldsymbol{\theta})}{d\theta_j)} = \left(\frac{dJ_{kk'}(\boldsymbol{\eta},\boldsymbol{\theta})}{d\theta_j}\right)_{k,k'}$$

and

$$\frac{dJ_{kk'}(\boldsymbol{\eta},\boldsymbol{\theta})}{d\theta_j} = \frac{\vartheta J_{kk'}}{\vartheta\theta_j} + \frac{\vartheta J_{kk'}}{\vartheta\boldsymbol{\eta}}\boldsymbol{v}_j. \tag{34}$$

Also,

$$\boldsymbol{w}_{jk}(t^*) = \frac{d}{d\theta_j}\boldsymbol{u}_k(t^*) = \frac{d}{d\theta_j}\boldsymbol{e}_k = \boldsymbol{0}.\square$$

Finally, if $\boldsymbol{V} = \boldsymbol{V}(t^*, t)$ in (25), the $n \times n$ matrix $\boldsymbol{X}_j = d\boldsymbol{V}/d\theta_j$ can be derived as the solution of

$$\frac{d\boldsymbol{X}_j}{dt} = \boldsymbol{X}_j\boldsymbol{J}^T + \boldsymbol{J}\boldsymbol{X}_j + \boldsymbol{V}\left(\frac{\vartheta J_{kk'}}{\vartheta\boldsymbol{\eta}}\boldsymbol{v}_j + \frac{\vartheta J_{kk'}}{\vartheta\theta_j}\right)_{k'k} + \left(\frac{\vartheta J_{kk'}}{\vartheta\boldsymbol{\eta}}\boldsymbol{v}_j + \frac{\vartheta J_{kk'}}{\vartheta\theta_j}\right)_{kk'}\boldsymbol{V}$$

$$+ \boldsymbol{M}\ \mathrm{Diag}\left(\frac{\vartheta\boldsymbol{u}_l}{\vartheta\boldsymbol{\eta}}^T\boldsymbol{v}_j + \frac{\vartheta\boldsymbol{u}_l}{\vartheta\theta_j}\right)_l\boldsymbol{M}^T, \qquad \boldsymbol{X}_j(t^*) = \boldsymbol{0} \tag{35}$$

**Proof.** We differentiate (25) wrt to $\theta_j$ to get

$$\begin{aligned}\frac{d}{d\theta_j}\left(\frac{d\boldsymbol{V}}{dt}\right) &= \frac{d}{d\theta_j}\left(\boldsymbol{V}\boldsymbol{J}^T + \boldsymbol{J}\boldsymbol{V} + \boldsymbol{M}U\boldsymbol{M}^T\right)\\ \frac{d\boldsymbol{X}_j}{dt} &= \boldsymbol{X}_j\boldsymbol{J}^T + \boldsymbol{V}\frac{d\boldsymbol{J}^T}{d\theta_j} + \frac{d\boldsymbol{J}}{d\theta_j}\boldsymbol{V} + \boldsymbol{J}(t)\boldsymbol{X}_j + \boldsymbol{M}\frac{d\boldsymbol{U}}{d\theta_j}\boldsymbol{M}^T\end{aligned}$$

The derivative $d\boldsymbol{J}/d\theta_j$ is derived in (34), while $\frac{\vartheta}{\vartheta\theta_j}\boldsymbol{U}(\boldsymbol{\eta},\boldsymbol{\theta})$ is a diagonal matrix with main diagonal entries,

$$\frac{\vartheta\boldsymbol{u}_l}{\vartheta\boldsymbol{\eta}}^T\frac{\vartheta\boldsymbol{\eta}}{\vartheta\theta_j} + \frac{\vartheta\boldsymbol{u}_l}{\vartheta\theta_j} = \frac{\vartheta r_l}{\vartheta\boldsymbol{\eta}}^T v_j + \frac{\vartheta r_l}{\vartheta\theta_{j)}}.\square$$

#### 4.8 FIM computation for pcLNA joint distributions

The multivariate normality of the pcLNA joint distributions $P_{\text{pcLNA}}(\boldsymbol{Q}_{\boldsymbol{x}_1}, \boldsymbol{Q}_{\boldsymbol{x}_2}, \ldots, \boldsymbol{Q}_{\boldsymbol{x}_k}|\boldsymbol{X}(0) \sim MVN(\boldsymbol{\mu}_0, \boldsymbol{\Sigma}_0))$, for $k = 2, \ldots,$ allows us to compute the corresponding FIM as in (7) by computing the derivatives of its mean vector $\boldsymbol{\mu}$ and precision matrix $\boldsymbol{A}$ in (29) and (30), respectively.

As part of this, the derivatives of $\vartheta\boldsymbol{x}(t_i)/\vartheta\boldsymbol{\theta}$, $\vartheta\boldsymbol{C}(t_i, t_{i+1})/\vartheta\boldsymbol{\theta}$, and $\vartheta\boldsymbol{V}(t_i, t_{i+1})/\vartheta\boldsymbol{\theta}$ are derived by solving the initial value problems in (32), (33) and (35), respectively. The derivatives $\vartheta\boldsymbol{R}_t^{(2)}/\vartheta\boldsymbol{\theta}$ are derived by using the fsolve function of MATLAB to derive a solution of the linear problem with constraints defined by differentiating the equations $\boldsymbol{F}(\boldsymbol{x}(t))^T\boldsymbol{R}_t^{(2)} = \boldsymbol{0}$ and ${\boldsymbol{R}_t^{(2)}}^T\boldsymbol{R}_t^{(2)} = \boldsymbol{I}$.

#### 4.9 Effects of observing less time-points on the singular values of the FIM

The singular values of the FIM in Fig. I3(b) are computed at time-points $t = 12, 22, 110, 142, 198, 244, 286, 344$min. These times include the first four peak times of the nuclear NF-$\kappa$B concentration, $t \approx 22, 142, 244, 344$ min, an earlier and three times between the peaks. If instead the singular values of the FIM are measured at time-points $t = 12, 22$ min, then the singular values are displayed in Fig. S8. The singular values are slightly increased compared to the base model, but the increase is clearly less prominent, particularly for the second singular value, compared to the corresponding singular values for the times $t = 12, 22, 110, 142, 198, 244, 286, 344$ min.

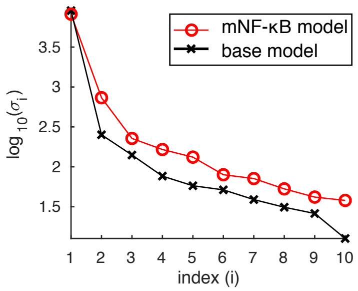

**Figure S8:** *The singular values of the NF-$\kappa$B base model (black -×- line) and the mNF-$\kappa$B model (red -o- line) for observations taken at times $t = 12, 22$ min.*

#### 5 NF-$\kappa$B target genes

We used the list of NF-$\kappa$B target genes provided in [5]. This comprises of the list of target genes taken from the Boston University web site at http://www.bu.edu/nf-kb/gene-resources/target-genes/ [1], combined with target genes found via an extensive ChIP-seq search performed in [15]. There are a total of 1361 genes in this list.

The profile data of the 8 target genes are displayed in Fig. S9 and S10. Fig. S11 illustrates how these 8 genes can be used to distinguish between 14 different experimental conditions. An approximate expression profile alphabet is provided, which includes profiles that can be described as no response, or early/medium/late, sustained or not sustained response. These profiles are identified by clustering [19]. Then we see on the table that each of these 14 different experimental conditions gave a different response combination over these 8 genes.

[1] accessed 8th August 2015

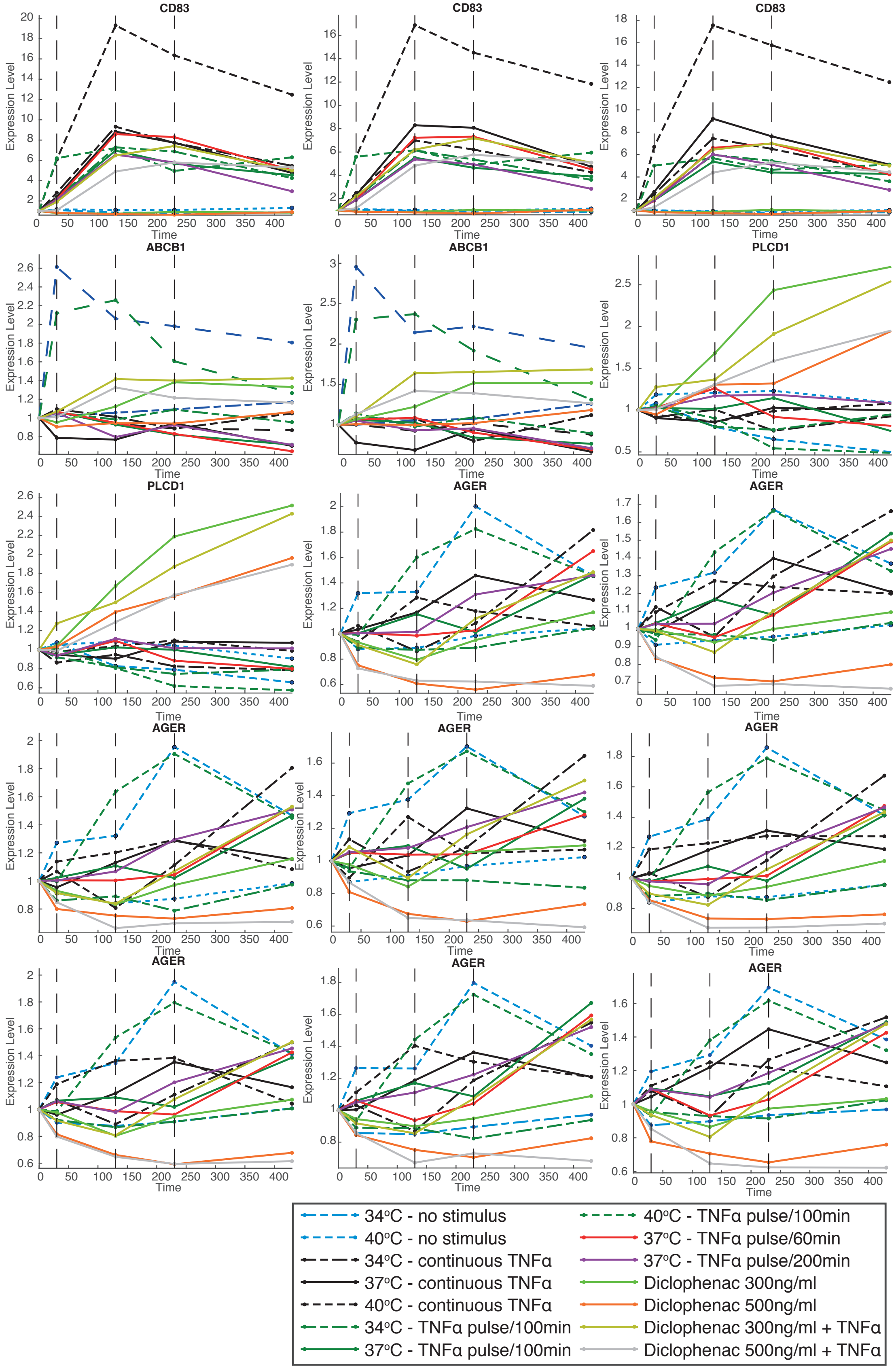

**Figure S9:** *Microarray profiles of the target genes presented in* **I** *Fig. 2. The condition is given in the legend. Some genes have multiple microarray data.*

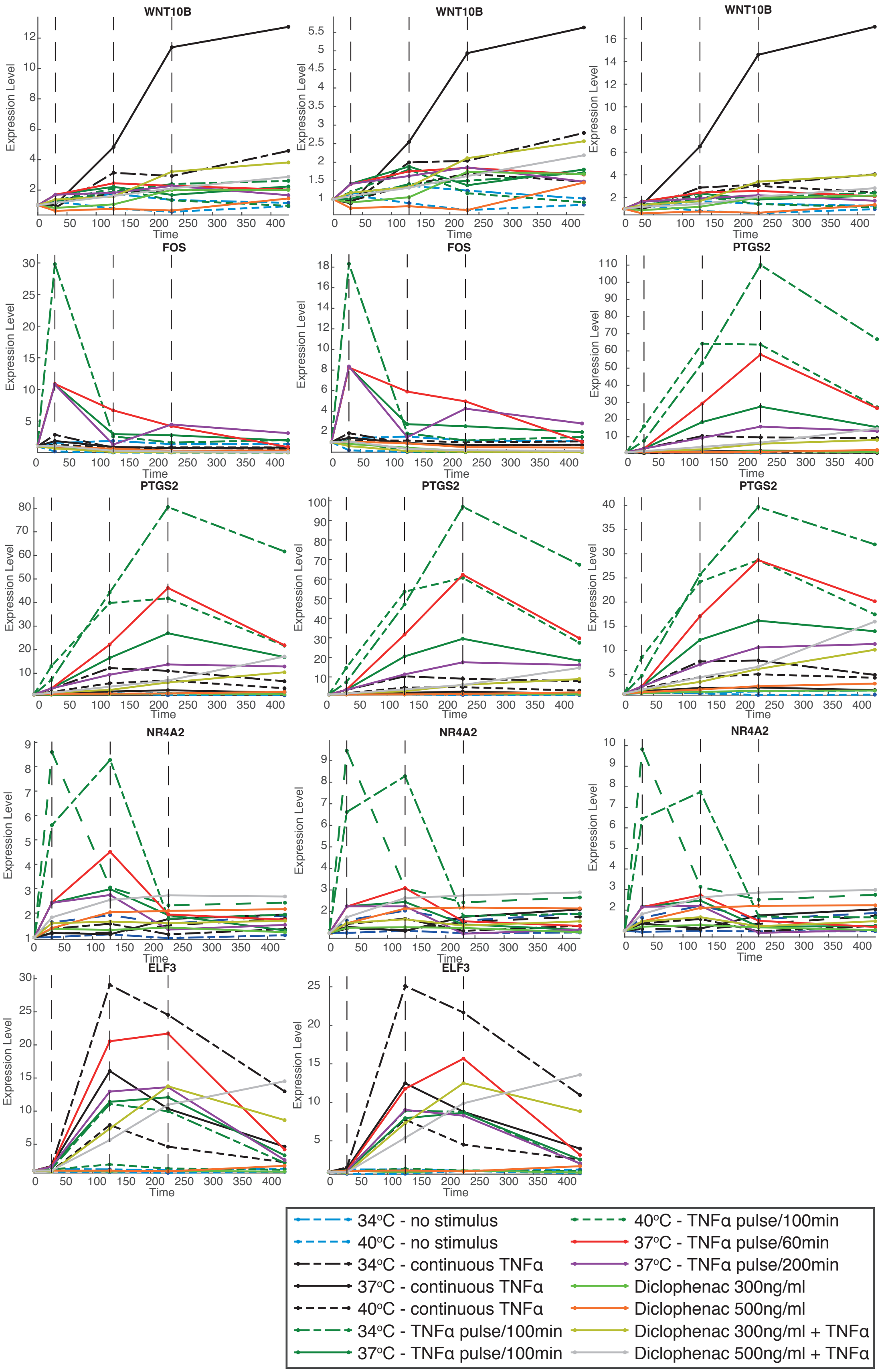

**Figure S10:** *Microarray profiles of the target genes presented in* **I** *Fig. 2. The condition is given in the legend. Some genes have multiple microarray data.*

| Condition | | | Gene | | | | | | | |
|---|---|---|---|---|---|---|---|---|---|---|
| °C | TNFα | Diclofenac | CD | AB | PL | AG | WN | EL | NR | CO |
| 40 | cont. | | a | b | b | e | b | i | b | b |
| 40 | | | b | a | b | e | b | b | b | b |
| 40 | 5/100 mins | | c | a | b | b | b | b | e | e |
| 34 | | | b | b | b | b | b | b | b | b |
| 37 | | 300 | b | b | d | b | b | b | b | b |
| 37 | | 500 | b | b | d | f | b | b | b | b |
| 37 | cont. | 300 | c | b | d | b | b | i | b | b |
| 37 | cont. | 500 | c | b | d | f | b | i | b | b |
| 37 | cont. | | c | b | b | b | g | i | b | b |
| 34 | cont. | | c | b | b | b | b | h | b | b |
| 34 | 5/100 mins | | c | b | b | b | b | i | j | e |
| 37 | 5/100 mins | | c | b | b | b | b | i | b | e |
| 37 | 5/200 mins | | c | b | b | b | b | b | b | b |
| 37 | 5/60 mins | | c | b | b | b | b | b | b | e |

Expression profile alphabet

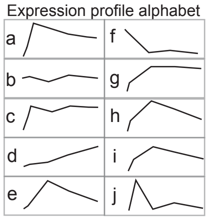

**TNFα treatment**
cont. = continuous
300 = 300ng/ml
500 = 500ng/ml
n/k mins = n mins pulse every k mins

**Genes**
**CD**83 **AG**ER **FOS**
**AB**CB1 **WN**T10B **NR**4A2
**PL**CD1 **EL**F3 **CO**X2

**Figure S11:** *Summary table of the expression of 8 genes (columns) that can distinguish the 14 different experimental conditions (rows). Each letter corresponds to an approximate expression profile given on the right. This includes profiles that can be described as either no response (b), early, medium or late, transient or sustained response. Reading the expression table from left to right and top to bottom, the colours emphasise the gene that identifies each condition. The profile data are given in Fig. S9 and S10*

[10] van Kampen NG. Stochastic Processes in Physics and Chemistry, Third Edition. Amsterdam: Elsevier. Boston and London: Elsevier; 2007.

[11] MATLAB 7.11 (R2017b), The MathWorks, Inc., Natick, Massachusetts, United States.

[17] Wilkinson DJ. Stochastic modelling for systems biology. Boca Raton: CRC Press/Taylor & Francis; 2012.

[18] Gibson MA, Bruck J. Efficient Exact Stochastic Simulation of Chemical Systems with Many Species and Many Channels The Journal of Physical Chemistry A, 104(9): 1876–1889

[19] Neal, R. M. (2000). Markov Chain Sampling Methods for Dirichlet Process Mixture Models. Journal of Computational and Graphical Statistics, 9(2), 249.
